# Oxaliplatin Resistance in Colorectal Cancer Enhances TRAIL Sensitivity Via Death Receptor 4 Upregulation and Lipid Raft Localization

**DOI:** 10.1101/2021.03.05.434100

**Authors:** Joshua D. Greenlee, Maria Lopez-Cavestany, Nerymar Ortiz-Otero, Kevin Liu, Tejas Subramanian, Burt Cagir, Michael R. King

## Abstract

Colorectal cancer (CRC) remains a leading cause of cancer death, and its mortality is associated with metastasis and chemoresistance. We demonstrate that oxaliplatin-resistant CRC cells are sensitized to TRAIL-mediated apoptosis. Oxaliplatin-resistant cells exhibited transcriptional downregulation of caspase-10, but this had minimal effects on TRAIL sensitivity following CRISPR-Cas9 deletion of caspase-10 in parental cells. Sensitization effects in oxaliplatin-resistant cells were found to be a result of increased DR4, as well as significantly enhanced DR4 palmitoylation and translocation into lipid rafts. Raft perturbation via nystatin and resveratrol significantly altered DR4/raft colocalization and TRAIL sensitivity. Blood samples from metastatic CRC patients were treated with TRAIL liposomes, and a 57% reduction of viable CTCs was observed. Increased DR4/lipid raft colocalization in CTCs was found to correspond with increased oxaliplatin resistance and increased efficacy of TRAIL liposomes. To our knowledge, this is the first study to investigate the role of lipid rafts in primary CTCs.

**Impact Statement:** Oxaliplatin-resistant colorectal cancer cells exhibit unregulated death receptor 4 expression with increased receptor palmitoylation and translocation into lipid rafts, increasing their sensitivity to apoptosis via TRAIL.

## Introduction

Colorectal cancer (CRC) is the second leading cause of cancer death and is responsible for over 50,000 deaths annually in the United States (1). The probability of being diagnosed with CRC in one’s lifetime is 1 in 24, and there are over 100,000 new cases diagnosed annually in the United States alone. While the 5-year survival rate of localized and regional disease is 90% and 71%, respectively, patients with metastatic disease have just a 14% 5-year survival rate (2). Dissemination to other organs is the cause of high mortality in most cancers, as nearly 90% of all cancer death is attributed to metastasis (3). The most common sites of CRC metastases include the liver, lungs, and peritoneum (peritoneal carcinomatosis). While surgery and radiation remain curative options for patients with localized disease, the standard of care for CRC patients with advanced metastatic disease is commonly combination front line chemotherapy treatment (4). These chemotherapy regimens typically include fluorouracil (5-FU) and leucovorin (LV) in combination, which work together to inhibit DNA and RNA synthesis and modulate tumor growth, extending median survival in patients from 9 months (with palliative care) to over 12 months (5). Oxaliplatin is a chemotherapeutic agent that upon binding to DNA, forms DNA adducts to cause irreversible transcriptional errors, resulting in cellular apoptosis. When oxaliplatin is administered with 5-FU/LV (FOLFOX), the objective response rate is 50% in previously untreated patients, increasing the median overall survival to 18 - 24 months (5, 6).

While there have been incremental advances in extending survival using FOLFOX and other oxaliplatin-containing chemotherapeutics, patients who eventually succumb to the disease frequently develop chemoresistant subpopulations of cancer cells via intrinsic or acquired mechanisms (6, 7). Mechanisms of oxaliplatin resistance in tumors include alterations in responses to DNA damage, cell death pathways (e.g., apoptosis, necrosis), NF-κB signaling, and cellular transport (7). Despite the robustness of these oxaliplatin-resistant cancer cells, multiple studies suggest that chemoresistant subpopulations may be increasingly susceptible to adjuvant therapies (7–12). Tumor necrosis factor related apoptosis inducing ligand (TRAIL) is a member of the TNF family of proteins and induces apoptosis in cancer cells via binding to transmembrane death receptors (13). The binding of TRAIL to trimerized death receptor 4 (DR4) and 5 (DR5) initiates an intracellular apoptotic cascade beginning with the recruitment of death domains and formation of the death-inducing signaling complex (DISC).

Lipid rafts (LRs) are microdomains in the plasma membrane lipid bilayer that are enriched in cholesterol and sphingolipids, with a propensity to assemble specific transmembrane and GPI-anchored proteins (14). Mounting evidence has demonstrated that lipid rafts play major roles in tumor progression, metastasis and cell death (15). Studies have shown that translocation into lipid rafts can augment signaling for a variety of cancer-implicated receptors, including growth factor receptors (IGFR and EGFR) and death receptors (Fas and Death receptors 4/5) (16–19). Translocation of death receptors into rafts enhances apoptotic signaling through the formation of clusters of apoptotic signaling molecule-enriched rafts (CASMER), which act as scaffolds to facilitate trimerization and supramolecular clustering of receptors (18). It has become increasingly evident that higher order oligomerization of death receptors is necessary for effective apoptotic signaling in cancer cells (20).

Studies have shown that combination treatment of chemotherapeutic agents with TRAIL may sensitize cancer cells to TRAIL-mediated apoptosis through a variety of mechanisms, including death receptor upregulation (21–23), suppression of apoptotic inhibitors within the intrinsic pathway (24), and redistribution of death receptors into lipid rafts (25). However, no study has examined whether surviving oxaliplatin-resistant subpopulations of cancer cells have an enhanced sensitivity to TRAIL. In this study, we demonstrate that oxaliplatin-resistant cells show enhanced sensitivity to TRAIL-mediated apoptosis through lipid raft translocation of DR4. Moreover, we elucidate mechanisms which drive this sensitization using chemoresistant cell lines and blood samples collected from metastatic cancer patients. The response of oxaliplatin-resistant CRC to TRAIL-based therapeutics may prove critical to establishing promising new adjuvants for patients who have exhausted conventional treatment modalities.

## Results

### Oxaliplatin-resistant cell lines show enhanced TRAIL sensitivity

Cell viability of four colorectal cancer cell lines after 24 hr treatment with 0.1-1000 ng/ml of TRAIL was measured and compared to the viability of oxaliplatin-resistant (OxR) cell lines (**Figure 1A**). Briefly, oxaliplatin-resistant cell lines were previously derived from exposure to increasing concentrations of oxaliplatin until a 10-fold increase in IC50 was achieved (26–28). Parental and OxR cells were treated with a range of oxaliplatin concentrations to ensure that chemoresistance was conserved after multiple passages in culture (**Figure 1-figure supplement 1A**). Moreover, oxaliplatin-resistant cells were found to have increased invasion and motility compared to parental cells, consistent with literature reporting their derivation (**Figure 1-figure supplement 1B**). Interestingly, oxaliplatin-resistant HT29, SW620 and HCT116 cell lines showed increased maximum TRAIL sensitization levels compared to their parental counterparts, while SW480 cells showed similar or decreased sensitization levels (**Figure 1-figure supplement 2**). IC50 values demonstrate that a chemoresistant phenotype resulted in augmented TRAIL-mediated apoptosis in 2 cell lines (**Figure 1B**). Importantly, cells were not treated with any oxaliplatin in quantifying the level of TRAIL sensitization, and oxaliplatin was not supplemented in the cell culture media to exclude any possible effects from combination treatment. In SW620 cells, large differences in apoptosis were observed when treated with the highest concentration of TRAIL (1000 ng/ml) (**Figure 1C**). Only 33.3% of parental cells were found to be in late-stage apoptosis after 24 hr, compared to 60.6% for OxR cells.

**Figure 1.**
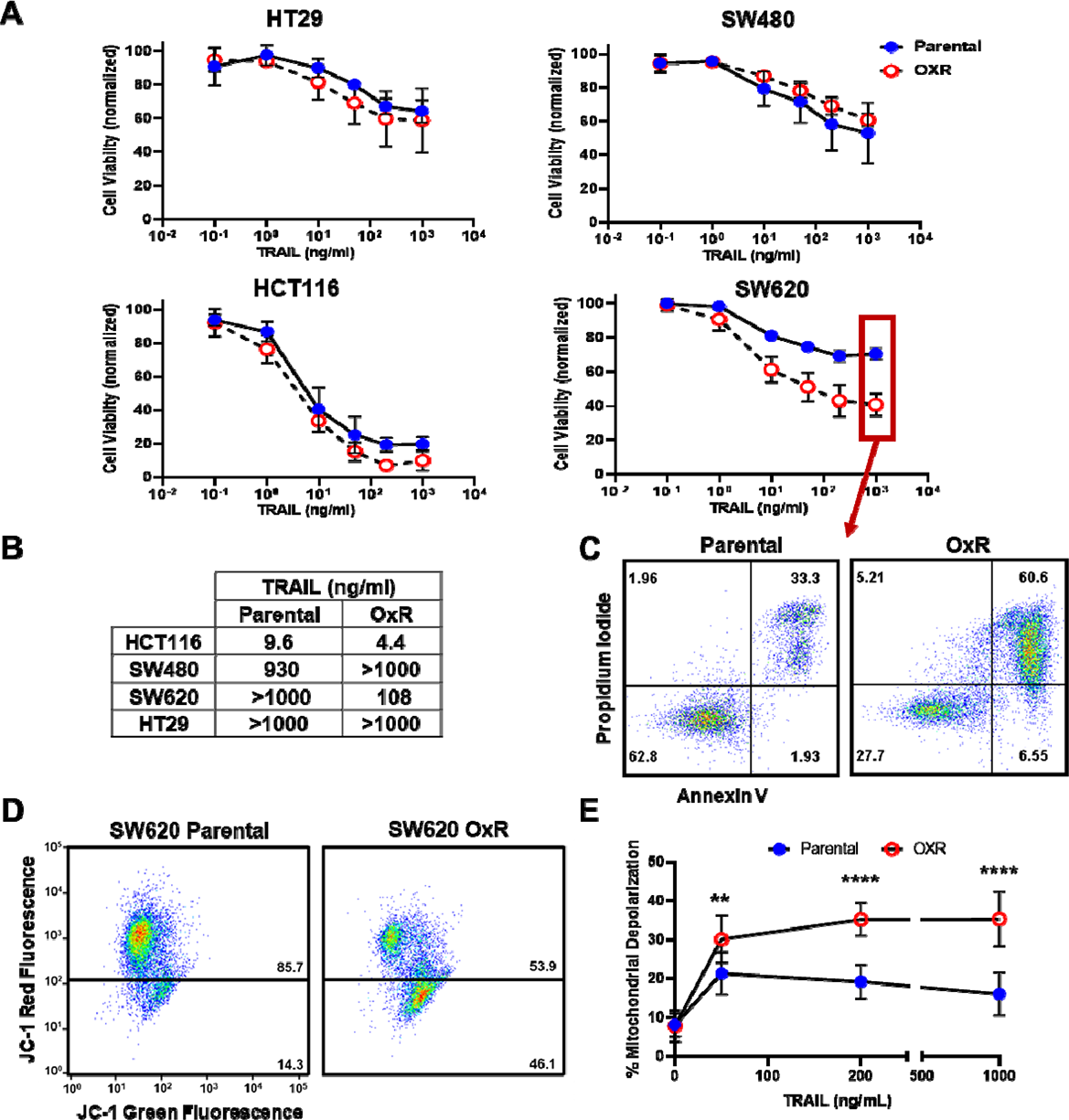
Oxaliplatin-resistant CRC cell lines exhibit enhanced sensitization to TRAIL-mediated apoptosis via the intrinsic pathway and mitochondrial permeabilization. **(A)** Oxaliplatin-resistant SW620, SW480, HCT116 and HT29 colon cancer cell lines demonstrate similar or enhanced sensitivity to TRAIL compared to their parental counterparts after 24 hr of treatment. N=3 (biological replicates); n=9 (technical replicates). **(B)** IC50 values were calculated using a variable slope four parameter nonlinear regression. **(C)** Representative Annexin V/PI flow plots comparing SW620 parental and OxR cell viability after 24 hr of treatment with 1000 ng/ml TRAIL. The four quadrants represent viable cells (bottom left), early apoptosis (bottom right), necrosis (top left) and late apoptosis (top right). **(D)** Representative flow plots of JC-1 assay after treatment with 1000 ng/ml of TRAIL. Mitochondrial depolarization is evidenced by decreased red fluorescence and increased green fluorescence. **(E)** Mitochondrial depolarization as a function of TRAIL concentration for SW620 parental and OxR cell lines. N=3 (n=9). For all graphs, data are presented as mean±SD. **p<0.01 ****p<0.0001 (unpaired two-tailed t-test).

To determine if the observed differences in apoptosis were due to enhanced mitochondrial outer membrane permeability, a JC-1 dye was used. SW620 OxR cells exhibited over a 3-fold increase in the population of JC-1 red (-) cells, indicating increased mitochondrial depolarization (**Figure 1D**). Mitochondrial depolarization was significantly enhanced in OxR cells for TRAIL concentrations of 50 ng/ml and higher (**Figure 1E**). Similar TRAIL-induced mitochondrial effects were observed in HCT116 cells (**Figure 1-figure supplement 3**). These results demonstrate that enhanced TRAIL mediated apoptosis is occurring, at least in part, via the intrinsic pathway and mitochondrial disruption.

### Oxaliplatin-resistant derivatives have decreased CASP10 that has little consequence on TRAIL sensitization

Given that enhanced TRAIL-mediated apoptosis was found to occur via the mitochondrial pathway, gene expression of apoptotic transcripts was compared between the parental and OxR cells. RT-PCR human apoptosis profiler arrays were used to analyze transcripts within the SW620 and HCT116 cell lines, since these cells showed the highest degree of OxR TRAIL sensitization and exhibit different innate sensitivities to TRAIL (HCT116 cells are TRAIL-sensitive whereas SW620 cells are TRAIL-resistant). Interestingly, upon analyzing the RNA expression of 84 apoptotic transcripts, both cell lines shared similar profiles between parental and OxR derivatives. HCT116 OxR cells showed upregulated pro-apoptotic transcripts cytochrome-c and caspase-4 (**Figure 2A**). Cytochrome-c is released from the mitochondria into the cytosol after mitochondrial permeabilization, binding to adaptor molecule apoptosis-protease activating factor 1 (Apaf-1) to form the apoptosome and initiate downstream caspase signaling (29). Caspase-4 is localized to the ER and initiates apoptosis in response to ER stress (30). Interestingly, SW620 OxR cells had upregulated Fas, a cell surface death receptor that acts similarly in apoptotic signaling to DR4/DR5 via binding of Fas ligand (31), and osteoprotegerin, a soluble decoy receptor that sequesters TRAIL and inhibits apoptosis (32) (**Figure 2B**). Upregulated Fas expression in SW620 OxR cells was confirmed via surface staining and flow cytometry, however, receptor neutralization with the ZB4 anti-Fas antibody had no effect on TRAIL sensitization when treated in combination with TRAIL (**Figure 2-figure supplement 1A-C**). Notably, both HCT116 and SW620 OxR cell lines had caspase-10 as the most significantly downregulated transcript. To determine whether this was of consequence to the observed TRAIL sensitization, an SW620 caspase-10 knockout cell line was created using a multi-guide sgRNA CRISPR-Cas9 approach. Knock-out (KO) efficiency was found to be 93% (**Figure 2C**). The TRAIL sensitivity of this caspase-10 KO cell line was compared to a control cell line treated with Cas9 only. Caspase-10 KO cells showed only a slight decrease in cell viability after 24 hr of TRAIL treatment (**Figure 2D**). The number of late-stage apoptotic cells remained similar between cell lines (**Figure 2E**) and the maximum TRAIL sensitization observed was insignificant following caspase-10 KO (**Figure 2F**).

**Figure 2.**
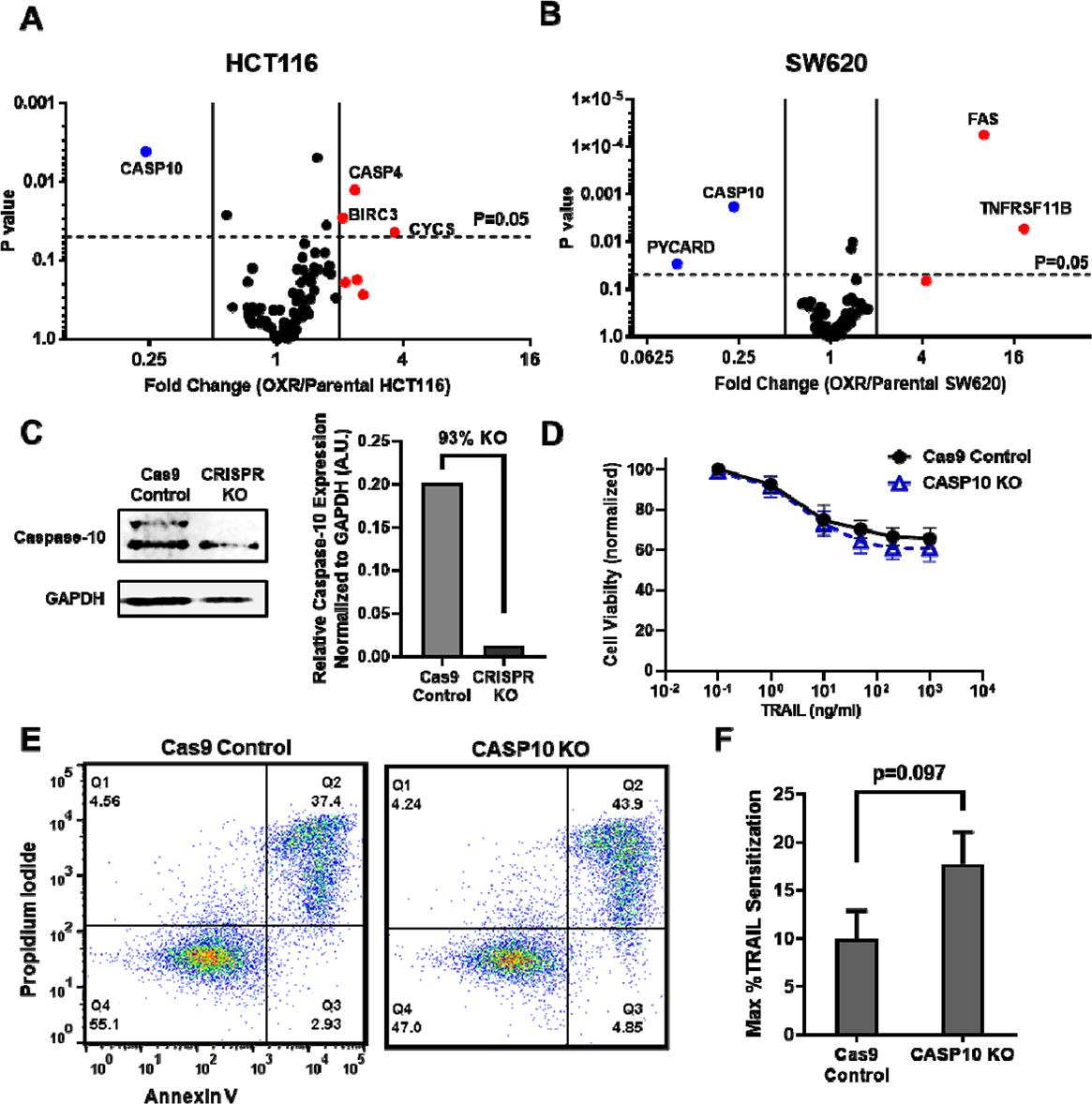
Microarray profiles show that parental and oxaliplatin-resistant CRC cell lines have similar expression of apoptotic transcripts while OxR derivatives have significantly downregulated CASP10. **(A-B)** Volcano plots of RT-PCR Apoptosis Profiler arrays demonstrate downregulation of CASP10 in OxR phenotypes. N=3. **(C)** CRISPR/Cas9 knockout of caspase-10 in SW620 parental cells was confirmed via western blot. sgRNA/Cas9 ribonucleoprotein complexes reduced caspase-10 expression by 93% compared to cells treated with Cas9 alone. **(D)** CASP10 KO cells demonstrate slight decreases in viability when treated with TRAIL compared to Cas9 control. Data are presented as mean±SD. N=3 (n=9). **(E)** Representative Annexin V/PI flow plots comparing SW620 parental (Cas9 only) and CASP10 KO cell viability after 24 hr of treatment with 1000 ng/ml TRAIL. **(F)** Depletion of caspase-10 did not have a significant effect on TRAIL sensitization (unpaired two-tailed t-test). Data are presented as mean+SEM. N=3 (n=9).

### TRAIL-sensitized OxR cell lines have upregulated DR4

While changes in death receptor expression were insignificant at a transcriptional level, studies have demonstrated that chemoresistance can alter receptor abundance via mechanisms of translational regulation (33, 34). Confocal microscopy showed that oxaliplatin-resistant cells have increased DR4 in both HCT116 (**Figure 3A**) and SW620 (**Figure 3B**) cell lines. To quantify receptor expression, total DR4 area per cell was analyzed for at least 70 cells. Analysis showed oxaliplatin-resistant derivative cell lines had significantly increased DR4 area per cell (**Figure 3C**). There were no differences in cell size between parental and OxR derivatives for all four cell lines (**Figure 3-figure supplement 1**). Flow cytometry staining of non-permeabilized cells was used to determine if this death receptor increase was also observed on the cell surface. Both HCT116 OxR and SW620 OxR cells showed significant increases in DR4 surface expression (**Figure 3D**). Total and surface DR4 expression was similar between parental and OxR derivatives in mildly sensitized HT29 cells and unsensitized SW480 cells (**Figure 3-figure supplement 2A-C**). To account for possible thresholding effects in area quantification, raw integrated density counts per cell were also measured and found to be consistent with changes in receptor area (**Figure 3-figure supplement 3**). Increases in DR4 expression of OxR derivatives was also confirmed via western blot but was only significant in SW620 cells (**Figure 3E-F**). Oxaliplatin-resistant HCT116, SW620 and HT29 cells all displayed increases in DR5 area per cell, while SW480 OxR cells had significant decreases in total DR5 expression (**Figure 3-figure supplement 4A-D**). However, total receptor area per cell was considerably lower for DR5 compared to DR4. Additionally, expression of surface DR5, analyzed via flow cytometry, was only significantly upregulated in SW620 OxR cells (**Figure 3-figure supplement 4E**). This is further confounded by western blot data which show no change in DR5 expression in HCT116 OxR cells, and a significant decrease in SW620 OxR cells (**Figure 3-figure supplement 5A-B**). Decoy receptors are surface receptors that, like death receptors, can bind to exogenous TRAIL. However, decoy receptor 1 (DcR1) and decoy receptor 2 (DcR2) are unable to activate the apoptotic pathway, making these receptors sequestering agents that competitively bind to TRAIL. While some studies have shown that chemotherapy-induced changes in TRAIL sensitivity have been linked to modulation or augmentation of decoy receptors (35), all cell lines exhibited no meaningful difference in surface DcR1 and DcR2 expression between parental and OxR derivatives (**Figure 3-figure supplement 6**). Despite statistical significance in SW480 and HT29 cells, decoy receptor expression, especially DcR2, was expressed in negligible quantities in these cell lines.

**Figure 3.**
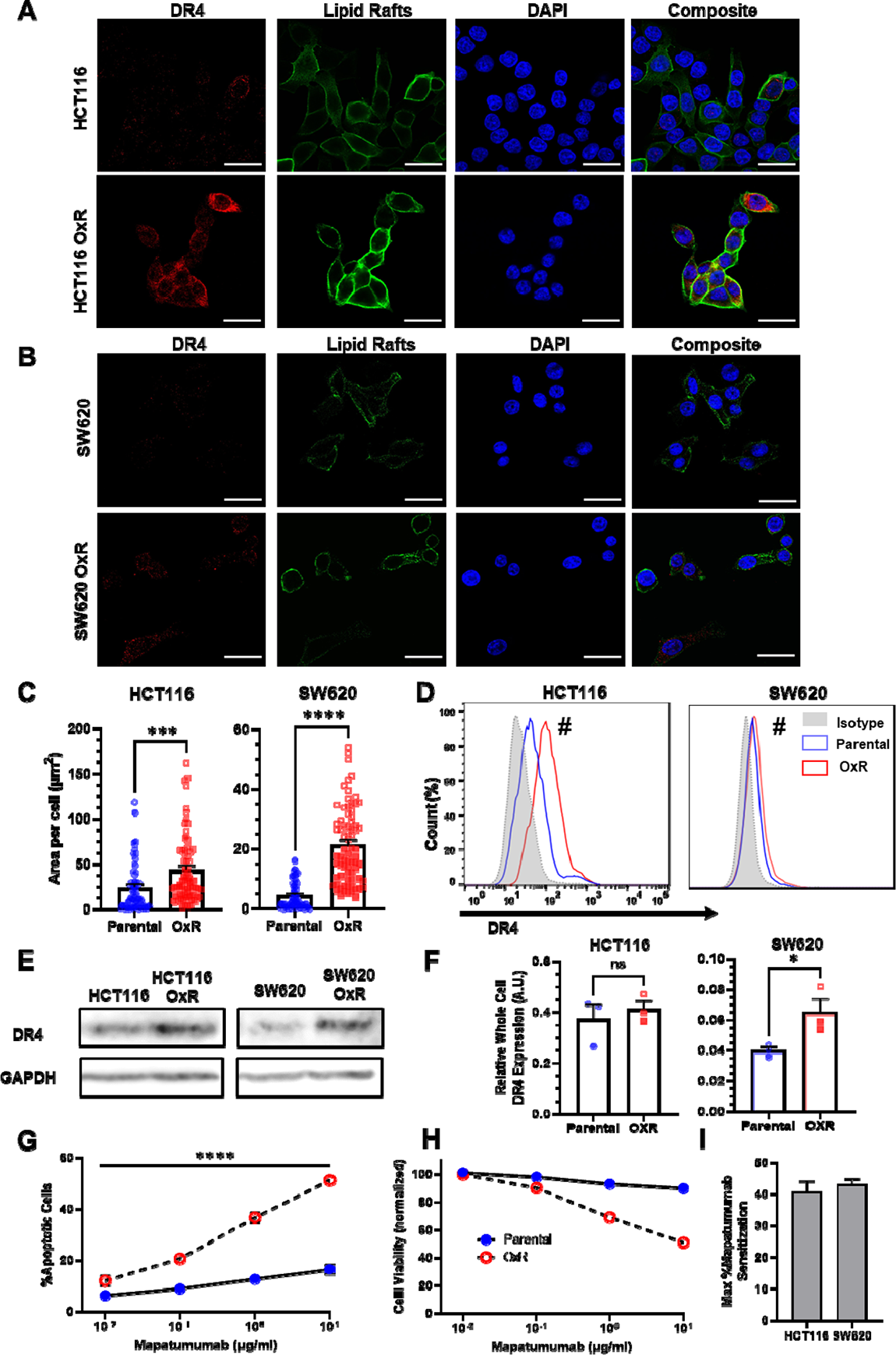
Oxaliplatin-resistant colon cancer cell lines have upregulated DR4 expression. **(A-B)** Confocal micrographs of HCT116 and SW620 cells, respectively. Red channel represents DR4, green is lipid rafts and blue is DAPI (nuclei). Scale bar = 30 μm. **(C)** Quantification of DR4 area per cell in HCT116 and SW620 cells. For each cell line, N=^μ^75 cells were analyzed. Data are presented as mean+SEM from N=3 independent experiments. ***p<0.001 ****p<0.0001 (unpaired two-tailed t-test). **(D)** OxR cells had increased surface expression of DR4 in non-permeabilized cells analyzed via flow cytometry. ^#^Significant according to a chi-squared test (see Supplementary File 1). **(E)** Western blots for DR4 in whole cell lysates of parental and OxR cells. **(F)** Quantification of western blots from three independent experiments (N=3). Data are presented as mean+SEM. *p<0.05 (unpaired two-tailed t-test). **(G)** Percentage of apoptotic SW620 cells after treatment with 0.01-10 µg/ml Mapatumumab (sum of early and late-stage apoptotic cells from Annexin/PI staining). Data are presented as mean+SD. N=3 (n=9). ****p<0.0001 (multiple unpaired two-tailed t-tests). **(H)** Cell viability of SW620 cells after Mapatumumab treatment, determined by AnnexinV/PI staining. Data are presented as mean±SD. N=3 (n=6). **(I)** Maximum Mapatumumab sensitization within OxR cell lines compared to their parental counterparts. Data are presented as mean+SEM.

Given the consistency in data suggesting DR4 upregulation in TRAIL-sensitized OxR cell lines, cells were treated with the DR4-agonist monoclonal antibody Mapatumumab to determine DR4 specificity. SW620 OxR cells exhibited significant increases in the number of apoptotic cells after 24 hours of treatment, including a 3-fold increase in apoptosis at concentrations of 10 µg/ml (**Figure 3G**). Cell viability closely paralleled TRAIL treatments: parental cells remained resistant at high doses while OxR cells exhibited a dose-responsive decrease in cell viability (**Figure 3H**). HCT116 cell lines exhibited similar results, as OxR cells were significantly more apoptotic at concentrations exceeding 0.1 µg/ml (**Figure 3-figure supplement 7A-B**). The maximum Mapatumumab sensitization was calculated to be greater than 40% for oxaliplatin-resistant HCT116 and SW620 cells (**Figure 3I**), providing more causal evidence for a DR4-associated mechanism.

### TRAIL-sensitized OxR cell lines have enhanced colocalization of DR4 into lipid rafts

Binary projections of colocalization events between DR4 and lipid rafts demonstrate that OxR phenotypes had enhanced DR4 translocation into lipid rafts in HCT116 and SW620 cells (**Figure 4A**). Quantification of total area of colocalization events showed that HCT116 OxR and SW620 OxR cells have significantly enhanced DR4 localized into lipid rafts, each with an over four-fold increase (**Figure 4B**). The areas of DR4/LR colocalized events per cell were not significantly different in HT29 and SW480 cells (**Figure 3-figure supplement 2D**). Other methods of colocalization analysis, including calculation of the Manders’ Correlation Coefficient (MCC), supported these results, specifically in HCT116 and SW620 cell lines where the Manders’ overlap was significantly greater in OxR cells (**Figure 4-figure supplement 1**). The fold change in DR4/LR colocalization area between OxR and parental cells exhibited a strong linear correlation (0.86) with TRAIL sensitization (**Figure 4C**). Colocalization of lipid rafts with DR5 was significantly enhanced only in HCT116 and HT29 cells, and analysis of the correlation between DR5 colocalization and TRAIL sensitization resulted in a weaker correlation of 0.48 (**Figure 4-figure supplement 2A-C**). Quantification of lipid raft area per cell revealed insignificant changes between parental and OxR derivatives in all cell lines except for HT29 cells, where parental cells showed significantly more rafts (**Figure 4-figure supplement 3**).

**Figure 4.**
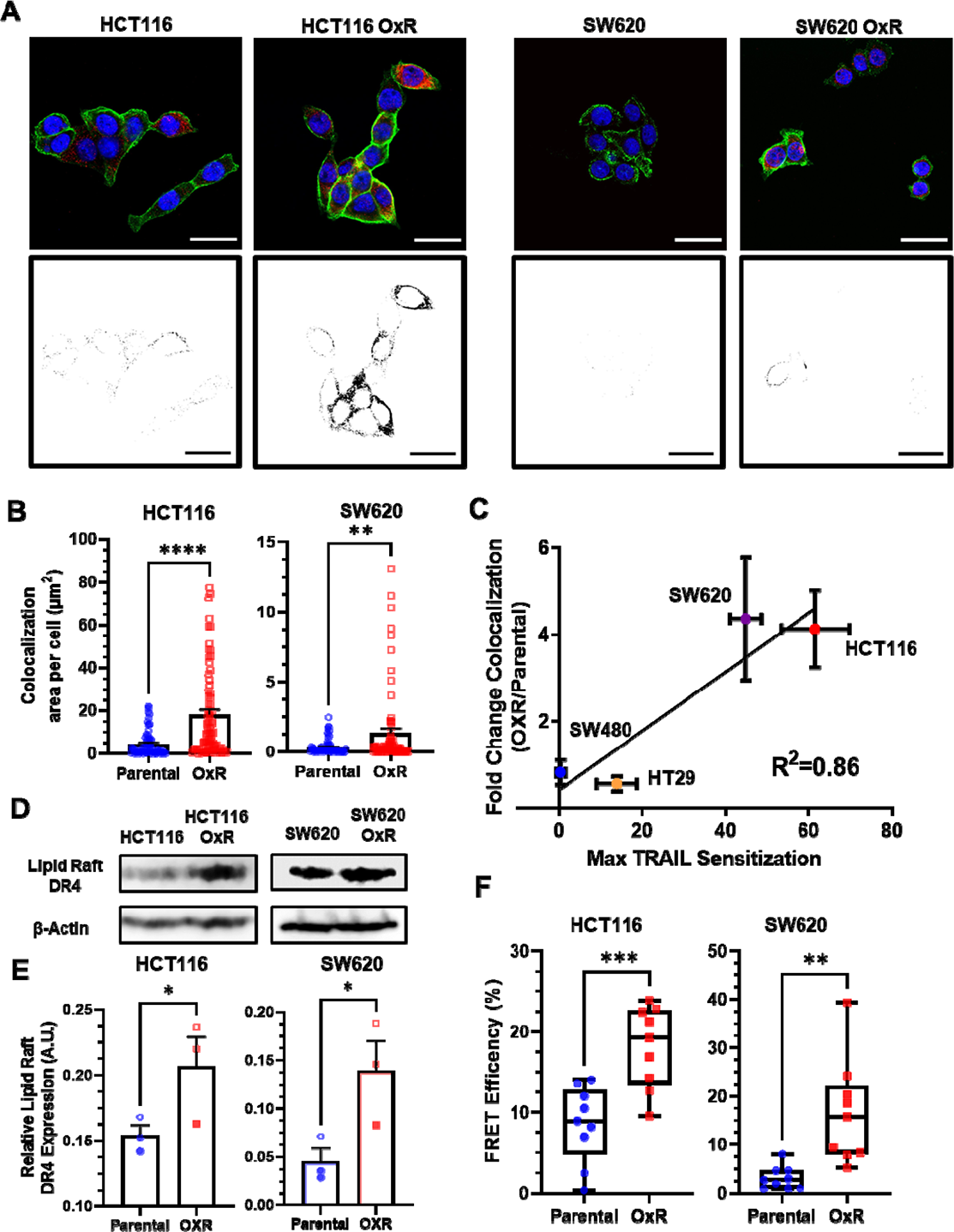
Oxaliplatin-resistant colon cancer cell lines have enhanced colocalization of DR4 into lipid rafts. **(A)** Composite images and binary projections of DR4/LR colocalization areas in HCT116 and SW620 cell lines. Lipid raft and DR4 binary images were generated for a specified threshold, then multiplied by one another to generate images with positive pixels in double positive areas. Red is DR4, green is lipid rafts and blue is DAPI. Scale bar = 30 μ Quantification of DR4 and lipid raft colocalization area per cell in HCT116 and SW620 cells. For each cell line, N=75 cells were analyzed. **p<0.01 ****p<0.0001 (unpaired two-tailed t-test). (**C)** Correlation between the fold change in DR4/LR colocalization (OxR phenotype/parental) and maximum TRAIL sensitization observed by the OxR phenotype for each of the four cell lines (simple linear regression analysis). **(D)** Lipid raft fractions were isolated and analyzed for DR4 via western blot in parental and OxR cells. **(E)** Quantification of lipid raft DR4 blots in (D). *p<0.05 (unpaired two-tailed t-test). For all graphs, data are presented as mean+SEM. **(F)** FRET efficiencies of FITC-labeled DR4 (donor) and Alexa 555-labeled lipid rafts (acceptor) in parental and OxR cells analyzed via flow cytometry. **p<0.01 ***p<0.001 (unpaired two-tailed t-test).

To confirm DR4 redistribution into rafts, western blots for DR4 were run on plasma membrane-derived lipid raft fractions, isolated using non-ionic detergent and centrifugation. Isolated lipid raft fractions exhibited significant increases in DR4 for both HCT116 OxR and SW620 OxR cells (**Figure 4D-E**). -actin was used as a loading control to compare relative DR4 expression between parental and OxR cell lines (36). There were no detectable levels of DR5 in western blots from lipid raft isolated fractions (**Figure 4-figure supplement 2D**). This is consistent with studies in hematological cancers that demonstrate raft localization of DR4 but not DR5 (17, 37). To further examine the proximity of DR4 and lipid rafts between phenotypes, Förster resonance energy transfer (FRET) efficiency was measured using a previously described flow cytometry protocol and calculated via a donor quenching method (38). Both HCT116 and SW620 OxR cells had significantly increased FRET efficiencies compared to their parental counterparts, with an over two-fold and five-fold increase, respectively (**Figure 4F**).

### Altering lipid raft composition affects DR4/LR colocalization and has consequential effects on TRAIL sensitization

To probe the effects of LR modulation on DR4 clustering and TRAIL sensitization, SW620 OxR and HCT 116 OxR cells were treated with 5 µM of nystatin, a cholesterol-sequestering agent that inhibits LR formation, in combination with TRAIL for 24 hr (**Figure 5A, G**). Nystatin inhibited TRAIL-mediated apoptosis in SW620 OxR cells, significantly decreasing the maximum TRAIL sensitization from 55% to 23%. (**Figure 5B**). Nystatin treatment was found to decrease DR4/LR colocalization by over 20-fold (**Figure 5C, M).** Similar results were found in HCT116 OxR cells, as nystatin treatment decreased TRAIL sensitization from 62% to 1% (**Figure 5H**), and decreased DR4/LR colocalization by over nine-fold (**Figure 5I**). To demonstrate that enhancing LR formation would have pro-apoptotic effects, parental cells were treated with 70 μ of resveratrol in combination with TRAIL for 24 hr (**Figure 5D, J**). Resveratrol has been shown to stabilize liquid-ordered domains in the plasma membrane and promote cholesterol/sphingolipid enriched lipid rafts (39). Resveratrol significantly sensitized parental SW620 cells to TRAIL irrespective of TRAIL concentration with a maximum TRAIL sensitization of 68% (**Figure 5E**). Treatment with resveratrol coincided with significant augmentation of DR4/LR colocalization area, an increase of over six-fold **(Figure 5F, N)**. Similarly, parental HCT116 cells treated with resveratrol were sensitized 59% (**Figure 5K**), corresponding with a nearly seven-fold increase in DR4/LR colocalization area per cell (**Figure 5L**). Resveratrol and nystatin had no significant effects on DR5 lipid raft colocalization, except in SW620 OxR cells where nystatin treatment surprisingly resulted in a slight increase in colocalization (**Figure 5-figure supplement 1A-B**).

**Figure 5.**
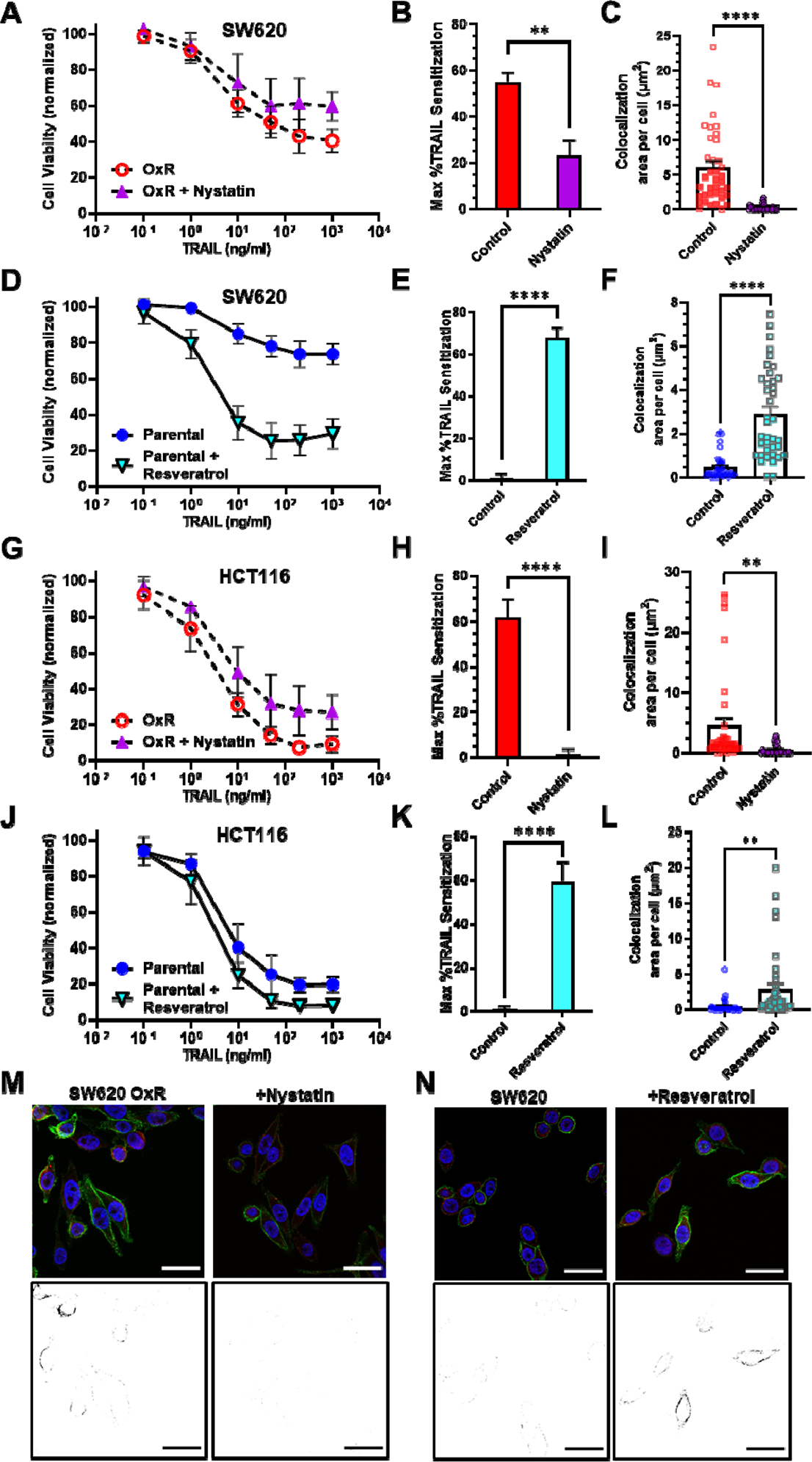
Pharmacological perturbation of DR4 localization in lipid rafts significantly alters cellular apoptosis in response to TRAIL. **(A, G)** SW620 OxR and HCT116 OxR cells, respectively, treated for 24 hr with a combination of TRAIL and 5 µM nystatin. **(B, H)** SW620 OxR and HCT116 OxR cells, respectively, showed a significant decrease in TRAIL sensitization when treated in combination with nystatin. N=3 (n=9). **(C, I)** Treatment with 5 µM nystatin significantly decreased DR4/LR colocalization area in SW620 OxR and HCT116 OxR cells, respectively. For each cell line, N=40 cells were analyzed. **(D, J)** SW620 Par and HCT116 Par cells, respectively, treated for 24 hr with a combination of TRAIL and 70 µM resveratrol. N=3 (n=9). **(E, K)** SW620 Par and HCT116 Par cells, respectively, showed a significant increase in TRAIL sensitization when treated in combination with resveratrol. N=3 (n=9). **(F, L)** Treatment with 70 µM nystatin significantly increased DR4/LR colocalization area in SW620 Par and HCT116 Par cells, respectively. For each cell line, N=40 cells were analyzed. **(M)** Representative composite images and binary projections of DR4/LR colocalization in SW620 OxR cells before and after nystatin treatment. **(N)** Representative composite images and binary projections of DR4/LR colocalization in parental SW620 cells before and after resveratrol treatment. Red represents DR4, green is lipid rafts and blue is DAPI. Scale bar = 30 **p<0.01 ****p<0.0001 (unpaired two-tailed t-test for all graphs). **(A, D, G, J)** Data are presented as mean±SD. **(B, C, E, F, H, I, K, L)** Data are presented as mean+SEM.

### S-Palmitoylation of DR4 is enhanced in oxaliplatin-resistant cells

Palmitoylation is the reversible, post-translational addition of the saturated fatty acid palmitate to the cystine residue of proteins. Palmitoylation of DR4 has proven to be critical for receptor oligomerization and lipid raft translocation, both obligatory for effective TRAIL-mediated apoptotic signaling (40). S-palmitoylation of DR4 in SW620 parental and OxR cells was analyzed via protein precipitation, free thiol blocking, thioester cleavage of palmitate linkages, and exchange with a mass tag label to quantify the degree of palmitoylated protein. We discovered that DR4 has four distinct palmitoylated sites, the degree of which was enhanced in the oxaliplatin-resistant phenotype (**Figure 6A**). Quantifying the percentage of palmitoylated protein in relation to input fraction (IFC) and non-mass tag preserved controls (APC-) validated that oxaliplatin-resistant cells had a significantly higher percentage of DR4 that was palmitoylated (55% compared to 43%) (**Figure 6B**). To determine whether enhanced palmitoylation was specific to DR4 and not a ubiquitous characteristic of the OxR phenotype, total cellular protein palmitoylation was measured and analyzed via flow cytometry (**Figure 6-figure supplement 1A**). Fluorescent azide labeling of palmitic acid confirmed that total cellular palmitoylation was unchanged between parental and OxR cells (**Figure 6-figure supplement 1B**).

**Figure 6.**
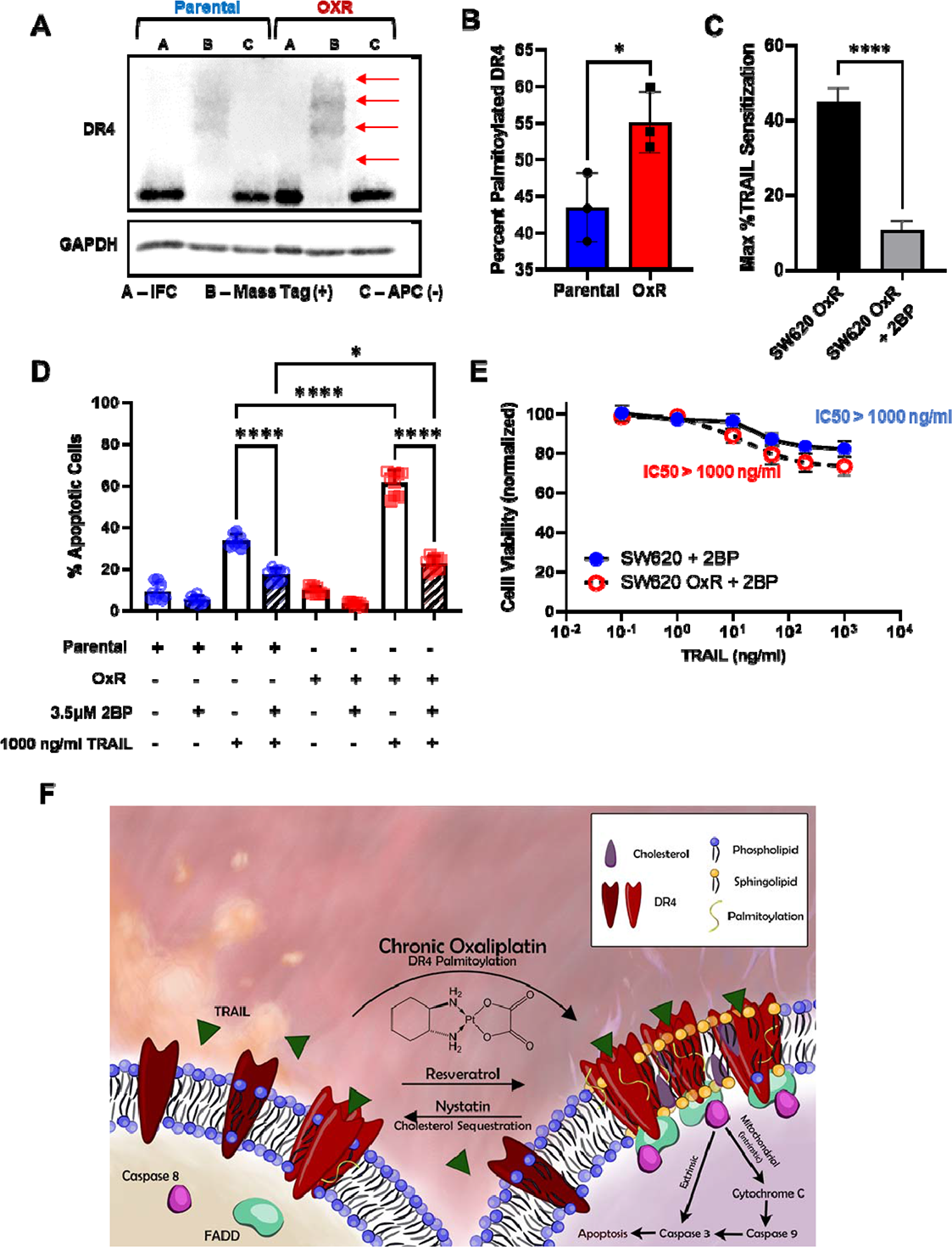
Oxaliplatin resistance enhances palmitoylation of DR4, selectively. **(A)** Death receptor palmitoylation was determined by protein precipitation, thioester cleavage, and conjugation of a mass tag to enumerate and quantify the degree of S-palmitoylation between cellular phenotypes. Samples with a mass tag “B” have distinct bands of equivalent increasing mass, with each mass shift indicating a palmitoylated site. Input fraction control (IFC) samples “A” were collected before thioester cleavage, while the acyl preservation negative control (APC) samples were incubated with an acyl-preservation reagent to block free thiols in place of the mass tag reagent. Arrows show palmitoylation bands. **(B)** Quantification of the percentage of palmitoylated DR4, calculated by dividing the total palmitoylated mass shift intensity by the average intensity of IFC and APC for each sample. Data are presented as mean±SD (N=3). *p<0.05 (unpaired two-tailed t-test). **(C)** Treatment with the irreversible palmitoylation inhibitor 2BP in combination with TRAIL significantly reduced TRAIL sensitization in SW620 OxR cells. Data are presented as mean+SEM. N=3 (n=9). *p<0.0001 (unpaired two-tailed t-test). **(D)** Percentage of apoptotic SW620 parental and OxR cells after treating with 1000 ng/ml TRAIL and 3.5 μ 2BP in combination (sum of early and late-stage apoptotic cells from Annexin/PI staining). Data are presented as mean+SD. N=3 (n=9). *p<0.05 ****p<0.0001 (ordinary one-way ANOVA–Tukey’s multiple comparison test). **(E)** Cell viability determined by AnnexinV/PI staining for cells treated with 0.1-1000 ng/ml TRAIL and 3.5 μ 2BP. IC50 values were calculated using a variable slope four parameter nonlinear regression. Data are presented as mean±SD. N=3 (n=9). **(F)** Proposed mechanism of enhanced TRAIL-mediated apoptosis in oxaliplatin-resistant cells.

To further examine the relationship between DR4 palmitoylation and TRAIL sensitization in OxR cells, the irreversible palmitoylation inhibitor 2-bromopalmitate (2BP) was used. 2BP is a commonly used palmitate analog that is thought to bind to palmitoyl acyl transferase, forming an inhibitory enzyme complex (41). Treating SW620 OxR cells with 3.5 μ 2BP in combination with TRAIL significantly reduced TRAIL sensitization and increased IC50 to over 1000 ng/ml (**Figure 6C, E**). 2BP significantly reduced the number of apoptotic cells in both parental and OxR cells, demonstrating the importance of DR4 palmitoylation in TRAIL signaling, particularly in chemoresistant cells (**Figure 6D**). These data suggest a novel mechanism for enhanced lipid raft-DR4 colocalization in OxR cells via enhanced DR4 palmitoylation (**Figure 6F**).

### Metastatic CRC patients show sensitivity to TRAIL liposomes despite chemoresistance

Despite promising specificity for cancer cells and low off-target toxicity, TRAIL’s translational relevance has been confounded by a short half-life and ineffective delivery modalities (42). In recent studies, our lab has demonstrated that TRAIL-coated leukocytes via the administration of liposomal TRAIL can be effective in eradicating circulating tumor cells (CTCs) in the blood of metastatic cancer patients (43). Briefly, liposomes were synthesized as previously described using a thin film hydration method, stepwise extrusion to 100 nm in diameter, and decoration with E-selectin and TRAIL via his-tag conjugation (44) (**Figure 7A**). Undecorated “control” liposomes, soluble TRAIL (290 ng/ml; at equivalent concentrations as TRAIL liposomes) and oxaliplatin (at peak plasma concentrations of 5 μ were used as controls. Blood was collected from 13 metastatic CRC patients who had previously undergone or were currently undergoing an oxaliplatin chemotherapy regimen (**Table 1**). Of these, five patients were analyzed at two to three timepoints over their respective treatment regimens, representing 21 total samples. Blood samples were treated with TRAIL liposomes or control treatments under hematogenous circulatory shear conditions in a cone-and-plate viscometer. TRAIL liposomes significantly decreased the average percentage of viable CTCs in patient blood to 43%, compared to just 86% when treated with oxaliplatin (**Figure 7B**). Interpatient variation was dominant in response to TRAIL liposome treatment, as the between-patient coefficient of variation was twice as high (CoV=0.55) as the average within-patient variation (CoV=0.28). Viable CTCs were categorized as cells that were cytokeratin(+), DAPI(+), CD45(-) and propidium iodide(-) (**Figure 7C**). TRAIL liposomal therapy reduced total viable CTC counts by 58% compared to control liposomes, and over 32% compared to oxaliplatin after just 4 hr in circulation (**Figure 7-figure supplement 1**). Notably, in two patients (P10 and P11), there were no detectable viable CTCs in blood samples treated with TRAIL liposomes. When categorizing patients by location of metastasis, patients that presented with metastases in the liver or bone showed a greater reduction in viable CTCs (69% and 71%, respectively) than patients with both lung and liver metastases (32%) (**Figure 7D**). Patients had similar CTC reductions regardless of their treatment at the time of blood draw, while those undergoing FOLFOX or capecitabine + oxaliplatin had the highest reduction in CTCs (65% and 60%, respectively) (**Figure 7E**). When categorizing patients as either oxaliplatin-sensitive or resistant, based on their response to 5 μM oxaliplatin under hematogenous circulatory-shear conditions (threshold 80% CTC viability), there was no significant difference in CTC response to TRAIL liposomes (**Figure 7-figure supplement 2A**). Likewise, grouping patients by those undergoing oxaliplatin chemotherapy and those who had failed oxaliplatin previously, there was no significant difference in reduction of viable CTCs from the administration of liposomal TRAIL (**Figure 7-figure supplement 2B**). This demonstrates the utility of TRAIL liposomes to eradicate CTCs in both oxaliplatin-sensitive and oxaliplatin-resistant patients.

**Figure 7.**
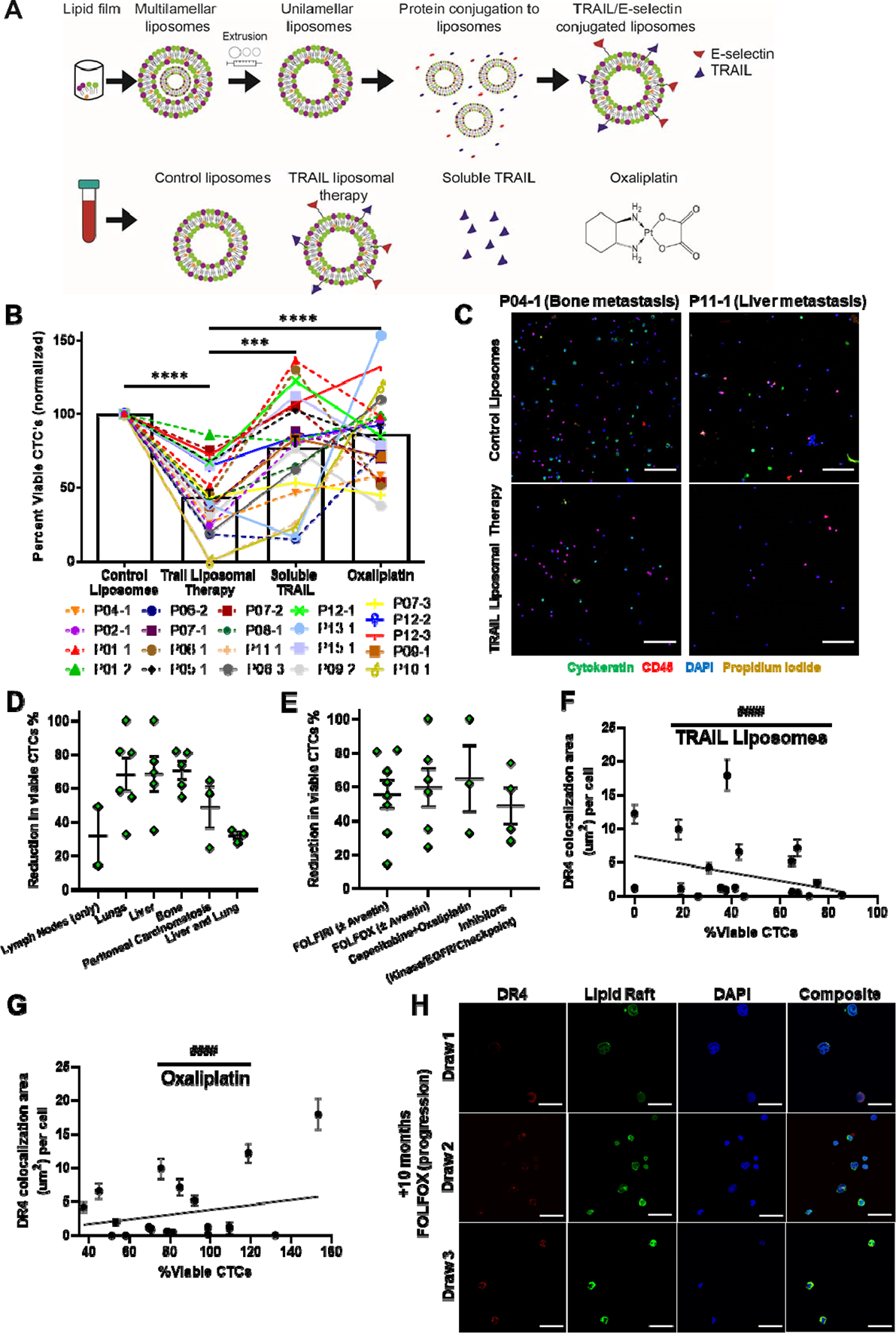
TRAIL-conjugated liposomes neutralize CTCs from the blood of patients with metastatic, oxaliplatin-resistant colorectal cancer. **(A)** Liposomes were synthesized using a thin film hydration method, followed by extrusion and his-tag conjugation of TRAIL and E-selectin protein. Patient blood samples were treated in a cone-and-plate viscometer under circulatory shear conditions with either control liposomes, TRAIL liposomes, soluble TRAIL, or oxaliplatin. **(B)** Effects of TRAIL liposomes and control treatments on the number of viable CTCs, normalized to control liposome treatment. Bars represent the average of all patients and time points. (N=21) **p<0.001 ****p<0.0001 (ordinary one-way ANOVA–Tukey’s multiple comparison test). **(C)** Representative micrographs of 2 patients showing neutralization of CTCs in TRAIL liposomes compared to control liposomes, stained for cytokeratin (green), DAPI (blue), CD45 (red), and propidium iodide (yellow). Scale bar = 100 µm. **(D, E)** Reduction in viable CTCs categorized by location of metastasis and treatment administered at the time of blood draw, respectively. **(F)** DR4/LR colocalization area of patient CTCs plotted against the percentage of viable CTCs following TRAIL liposome treatments. Each point corresponds with one patient draw. ^####^p<0.0001 (simple linear regression to confirm significant deviation from zero). **(G)** DR4/LR colocalization area of patient CTCs plotted against the normalized percentage of viable CTCs following oxaliplatin treatment. Each point corresponds with one patient draw. ^####^p<0.0001 (simple linear regression to confirm significant deviation from zero). **(H)** CTCs of Patient 7, stained for DR4 (red) and lipid rafts (green), demonstrating increased DR4/LR colocalization over the course of 10 months of FOLFOX treatment (with progressive disease despite treatment). Scale bar = 30 µm. For all graphs, data are presented as mean±SEM.

**Table 1.**
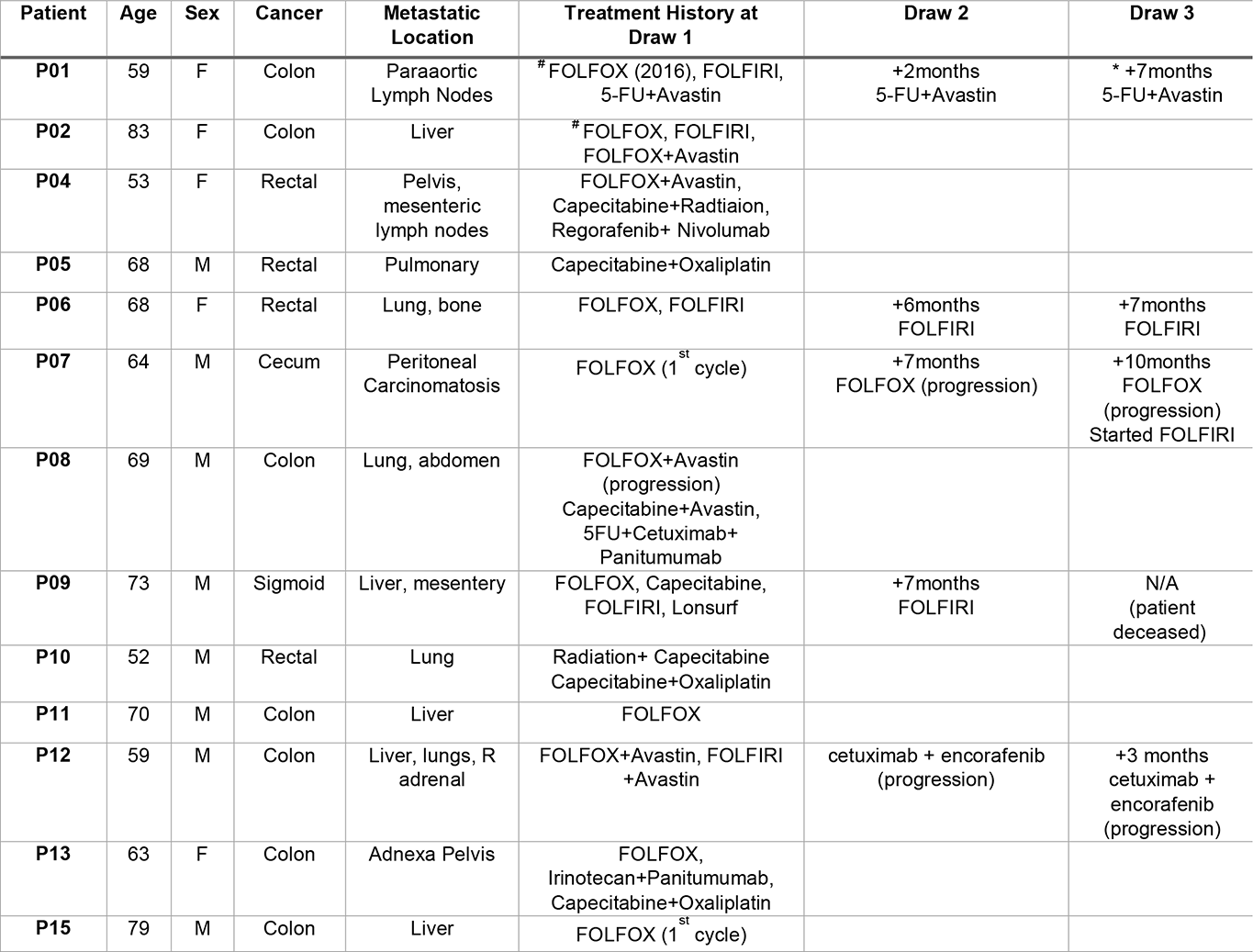
Demographic and clinical information of metastatic CRC patients enrolled in this study. ^#^Denotes missing DR4/LR analysis for this sample. *Denotes missing treatment analysis for this sample.

### CTC DR4-lipid raft colocalization corresponds with TRAIL liposome treatment efficacy and oxaliplatin resistance

Patient CTCs were also stained for DR4 and lipid rafts to examine the relationships between raft colocalization, treatment efficacy and oxaliplatin resistance. Decreasing lipid raft colocalization with DR4 coincided with reduced efficacy of TRAIL liposomes (higher percentage of viable CTCs after treatment), with a negative slope that significantly deviated from zero (**Figure 7F**). Additionally, increasing lipid raft DR4 corresponded with increasing resistance to oxaliplatin (higher percentage of viable CTCs after oxaliplatin treatment), with a positive slope that significantly deviated from zero (**Figure 7G**). These same trends were observed for total DR4 area (**Figure 7-figure supplement 2C-D**). Despite the small size of the patient cohort, these results are encouraging and support our *in vitro* data in OxR cell lines. Five patients provided multiple blood samples over the course of their treatment, as shown in **Table 1.** Of these, P07 was the only patient being treated with oxaliplatin (FOLFOX) over the course of all three blood draws. Patient 7 was undergoing the 1^st^ cycle of FOLFOX at the time of draw one and progressed while on FOLFOX for draws two and three. However, DR4 and lipid raft staining of CTCs revealed increased DR4/LR colocalization despite progression (**Figure 7H**). This same trend of enhanced CTC DR4/LR colocalization with treatment was observed in patients undergoing 5FU+Avastin (P01) and FOLFIRI (P09), while P06 (FOLFIRI) exhibited a bimodal response (**Figure 7-figure supplement 2E**). Interestingly, P12 exhibited decreased colocalization in CTCs over the course of treatment. This is hypothesized to be a result of a switch in treatment (FOLFIRI + Avastin to cetuximab + encorafenib) due to progression after the first draw.

## Discussion

Our lab has demonstrated the utility of TRAIL nanoparticles to treat a variety of cancer types *in vitro* (44), *in vivo* (45), and in clinical samples (43). While frontline chemotherapy remains a viable option for patients with metastatic CRC, long term treatment frequently leads to chemoresistance, consequently yielding a more aggressive, robust phenotype that is unresponsive to many systemic treatments (7). Our results demonstrate that OxR colorectal cancer cells are particularly susceptible to TRAIL-mediated apoptosis. Additionally, the ability to eradicate over 57% of oxaliplatin-resistant CTCs in patient blood demonstrates the utility of TRAIL liposomes clinically. Moreover, two patient samples exhibited 100% neutralization of all viable CTCs following *ex vivo* TRAIL liposomal treatment. This demonstrates the natural cancer cell targeting ability and low toxicity of TRAIL-based therapeutics, presenting a promising cancer management strategy for patients who have exhausted traditional treatment modalities. TRAIL’s apoptotic affect has been shown to be sensitized by circulatory shear stress, further supporting its use as an anti-metastatic therapy in the blood of patients (46). Multiple other studies have demonstrated that platin-based chemotherapeutics, including oxaliplatin, are able to sensitize cancer cells to TRAIL-mediated apoptosis when treated in combination (12,21,22,35). However, no study has investigated the effects of oxaliplatin resistance on TRAIL-mediated apoptosis, and importantly, no study has demonstrated that oxaliplatin-resistant cancers can be exploited with TRAIL therapies.

Elucidating the mechanisms that drive OxR TRAIL sensitization will be key in establishing personalized treatment strategies in patients. Interestingly, genetic analysis of TRAIL-sensitized OXR cells demonstrated that OxR cells consistently exhibited downregulated caspase-10. This may seem counterintuitive generally, since caspase-10 is a caspase-8 analog that initiates the apoptotic pathway after binding to FADD. However, studies have demonstrated the potential anti-apoptotic effects of high caspase-10 expression (47, 48). One recent study in particular demonstrated that upon activation with Fas ligand, caspase-10 reduced DISC association and activation of caspase-8, rewiring DISC signaling toward the NF-κB pathway and cell survival (49). However, this non-canonical caspase-10 signaling was found to have an insignificant effect on TRAIL sensitization as evidenced in experiments where caspase-10 was depleted in parental cells. This establishes that the observed augmentation of TRAIL sensitivity is likely a result of a translational or post-translational effect induced by oxaliplatin resistance, rather than a transcriptional change within the apoptotic pathway.

We have demonstrated that augmentation of death receptors, particularly DR4, in oxaliplatin resistant cells is one of the drivers of enhanced sensitization. One study found that cisplatin and 5-FU resistant side populations of colon cancer cells had upregulated DR4, consistent with our results (8). While microscopy data suggest that DR5 is upregulated in TRAIL-sensitized oxaliplatin-resistant cell lines, DR5 area per cell was considerably lower than DR4. Additionally, conflicting western blot and flow cytometry data make the case for DR5 augmentation in OxR cells less convincing. Treatment with the DR4 agonist antibody Mapatumumab validated the role of DR4 in the TRAIL-sensitization of OxR cells, as the differential treatment responses were analogous to that observed from TRAIL treatment. Interestingly, DR4 augmentation appears to be independent from transcriptional upregulation, as there was no significant change in mRNA expression between OxR and parental cell lines. Increasing evidence demonstrates that chemoresistance affects small non-coding microRNA (miRNA) expression, which modulates transcriptional and translational processes (33, 34). More specifically, studies have shown that oxaliplatin treatment and subsequent resistance in colorectal cancer cells alter miRNA expression, affecting signaling pathways within p53, epithelial-to-mesenchymal transition, and cell migration (28,50,51). Moving forward, future studies should examine the role of miRNA attenuation post-oxaliplatin resistance on the expression of death receptors, particularly DR4.

While sufficient DR4 expression is important for sustained apoptotic signaling, DR4 localization and compartmentalization within lipid rafts is unequivocally vital. Lipid rafts enhance the signaling capacity of surface receptors through a multitude of mechanisms (15). For example, LRs promote death receptor trimerization which is needed for signal transduction, act as concentrating platforms for DISC assembly and the recruitment of death domains, and protect DRs from internalization or enzymatic degradation (14). Additionally, juxtaposition of multiple DR trimers forms supramolecular entities, recently termed “CASMER” (18), capable of multivalent TRAIL signaling via extracellular pre-ligand assembly domains (PLADs) (20). Altering raft integrity via cholesterol sequestration using nystatin had profound impacts on reducing TRAIL sensitization within the OxR phenotype. Moreover, raft stabilization with resveratrol was able to enhance TRAIL sensitization within the parental phenotype, mirroring that observed in OxR cells. These changes in sensitivity were confirmed to coincide specifically with enhanced clustering of DR4 within rafts. These results are consistent with other studies which have shown that pharmacological alterations of lipid rafts have profound impacts on Fas and TRAIL toxicity (19, 52). Other studies have demonstrated that DR4 localization into lipid rafts is obligatory for TRAIL-induced apoptosis in hematological malignancies and non-small cell lung cancer, whereas DR5 has no dependence on raft translocation (17,20,53,54), consistent with our correlative data and receptor contents from lipid raft isolated membrane fractions. Additionally, one study found that oxaliplatin combination treatment with TRAIL in gastric cancer cells enhances apoptotic signaling through casitas B-lineage lymphoma (CBL) regulation and death receptor redistribution into lipid rafts (25). While it is evident that rafts promote CASMER formation, death receptor oligomerization, and TRAIL-mediated apoptosis, the mechanism linking the oxaliplatin-resistant phenotype and enhanced DR4 localization within rafts has yet to be studied.

We have demonstrated that a mechanism for this phenomenon is via enhanced DR4 palmitoylation. Palmitoylation is the post-translational covalent attachment of a fatty acid tail to cysteine residues in the protein transmembrane domain, influencing protein trafficking and signaling. There is evidence that both Fas receptor and DR4 are palmitoylated, while DR5 is not (40, 55). Furthermore, this post-translational modification has proven to be mandatory for DR4 oligomerization, lipid raft localization, and TRAIL-mediated apoptotic signaling (40). Interestingly, in a sensory neuron study in rats, palmitoylation of δ-Catenin in dorsal root ganglion was significantly increased after chronic oxaliplatin treatment (56). This is analogous to our results, as oxaliplatin-resistant colorectal cancer cells that have undergone chronic oxaliplatin treatment exhibited a higher percentage of palmitoylated DR4. Inhibiting palmitoylation with 2BP abrogated the TRAIL sensitizing effects within OxR cells, demonstrating the mandatory role palmitoylation has on DR4-mediated TRAIL signaling. Additionally, the fact that palmitoylation is inherent to DR4 and not DR5 explains why TRAIL sensitization of oxaliplatin-resistant cells strongly correlated with lipid raft translocation of DR4, but not DR5. Further studies probing the differences in palmitoylation between parental and OxR phenotypes are warranted to provide a more detailed understanding of oxaliplatin-induced palmitoylation of specific membrane proteins.

We have also shown that these results translate clinically, as DR4 expression and lipid raft colocalization of patient CTCs coincided with increased oxaliplatin resistance and increased neutralization of CTCs from TRAIL liposome treatment. Additionally, some metastatic CRC patients exhibited increased DR4/LR colocalization with ongoing chemotherapy cycles despite metastatic progression and worsening prognosis. To our knowledge, this is the first study investigating lipid raft/protein interactions in primary CTCs (15). Overall, our results demonstrate a novel mechanism for TRAIL sensitization in chemoresistant colorectal cancer cells via death receptor upregulation and localization within lipid rafts. However, since this sensitization was only observed in two of the four CRC cell lines tested, future studies should investigate genetic and phenotypic differences between these cell lines that may make some more susceptible than others to DR4 palmitoylation, augmentation, and localization. For the scope of this study, we chose to focus on the use of TRAIL treatment alone given its low toxicity and given our previous work in engineering TRAIL-conjugated delivery vehicles. However, since patients are treated with combination therapies, it would be valuable to investigate other therapeutics, such as curcumin or oxaliplatin, that synergize with TRAIL to treat oxaliplatin-resistant cancer cells (11, 24). Additionally, future studies should examine the TRAIL sensitization of oxaliplatin-resistant cells *in vivo* in orthotopic models of colorectal cancer metastasis (57). Examining the efficacy of TRAIL and TRAIL-conjugated nanoparticles to curb metastasis of oxaliplatin-resistant cells in humanized mouse models will provide translational evidence to support the mechanisms elucidated in this study. Moving forward, leveraging the enhanced signaling of death receptors in lipid rafts through mechanisms of drug delivery and lipid raft antagonization will be instrumental in therapeutic development for chemoresistant cancers.

## Acknowledgements

This work was funded by the National Institutes of Health, Grant No. R01CA203991 to M.R.K. We thank all the cancer patients who donated blood samples for this study. We also thank the Oncology Research Coordinator at Guthrie Clinic, Michelle Hunter, for supervising blood sample collection and shipment. Finally, the authors thank Dr. Mika Hosokawa and Dr. Lee Ellis for providing us with the oxaliplatin-resistant cell lines, as well as Matthew R. Zanotelli for assistance writing the colocalization macro in FIJI.

## Methods

### Key Resource Table

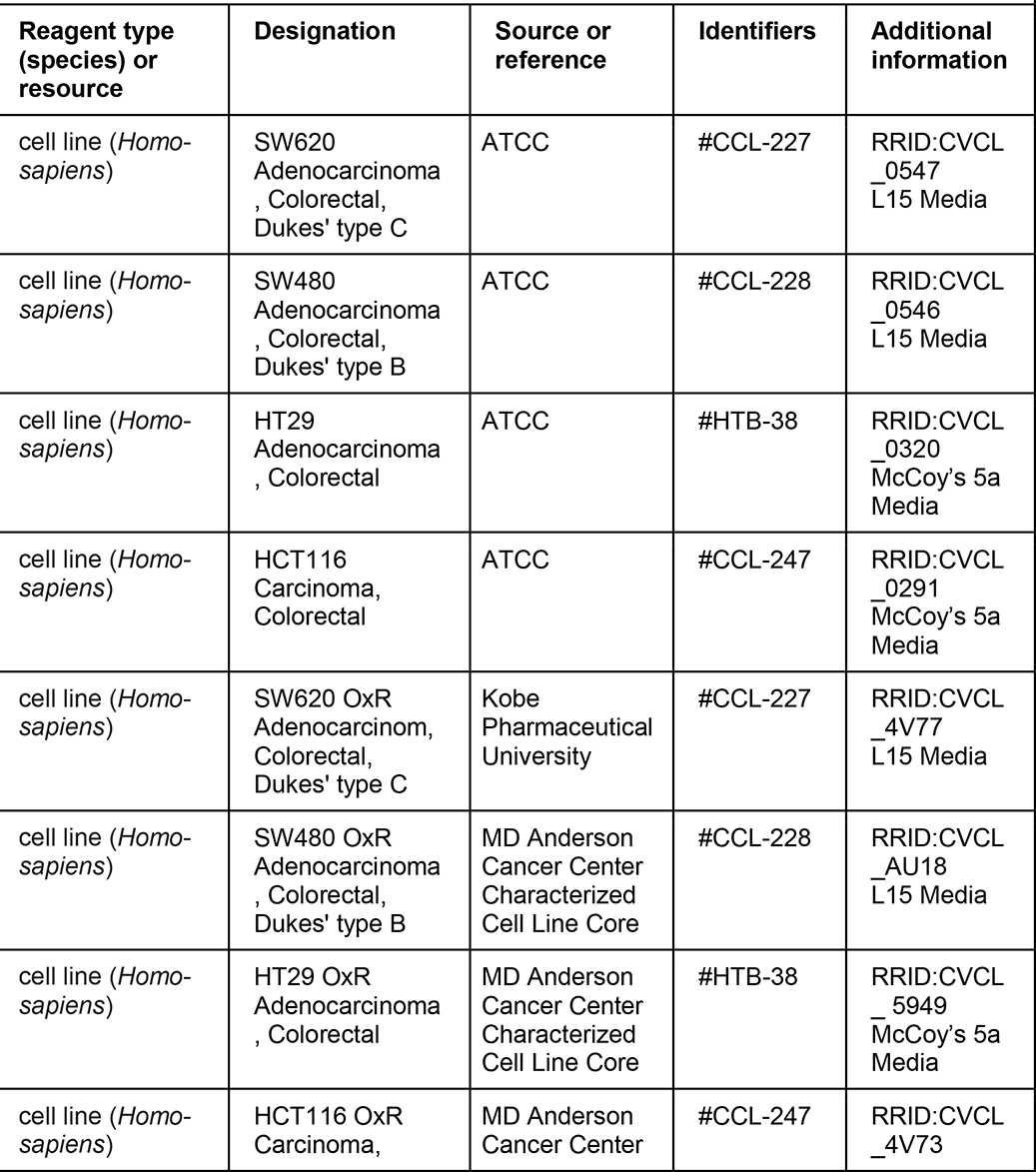
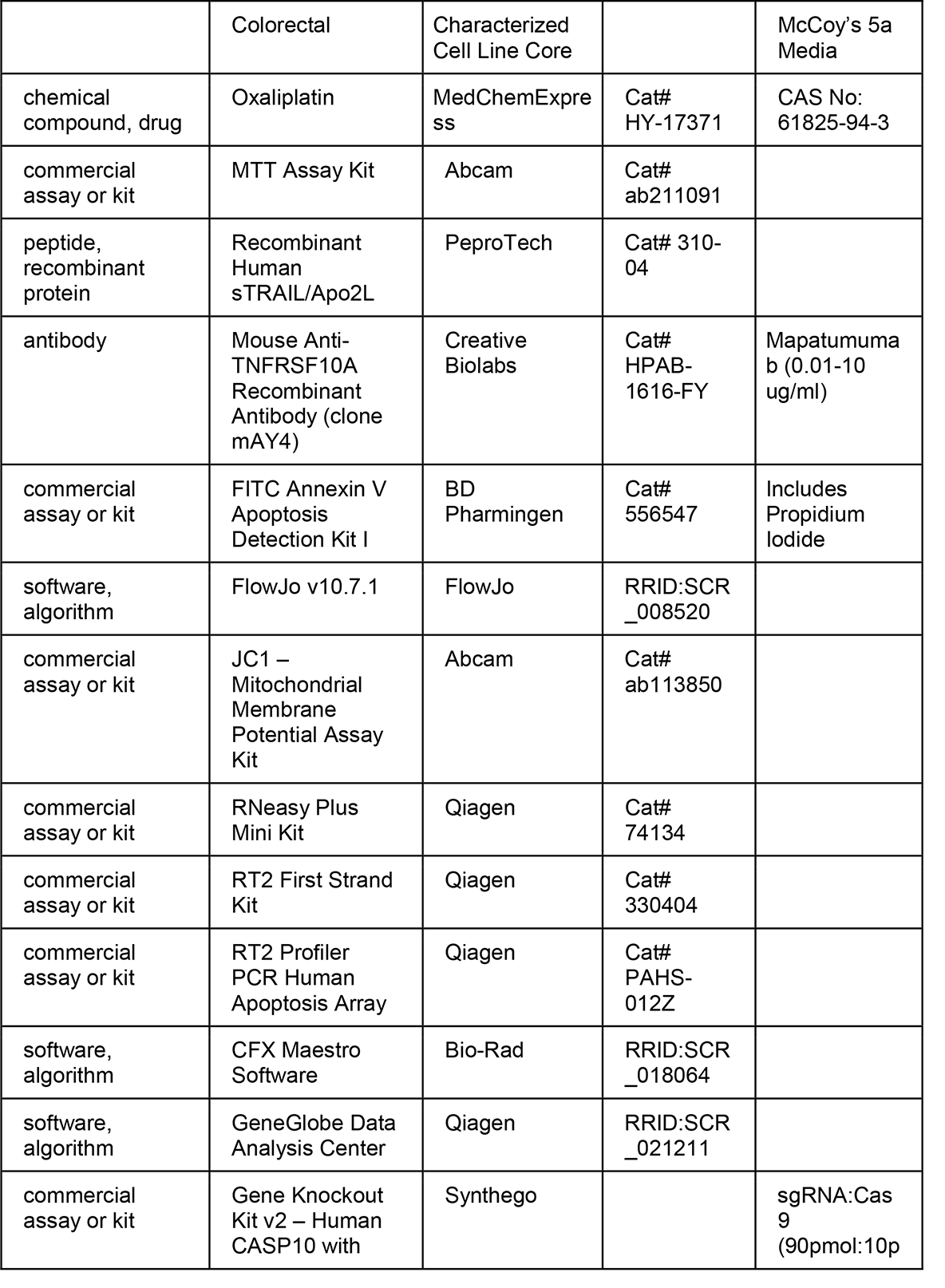
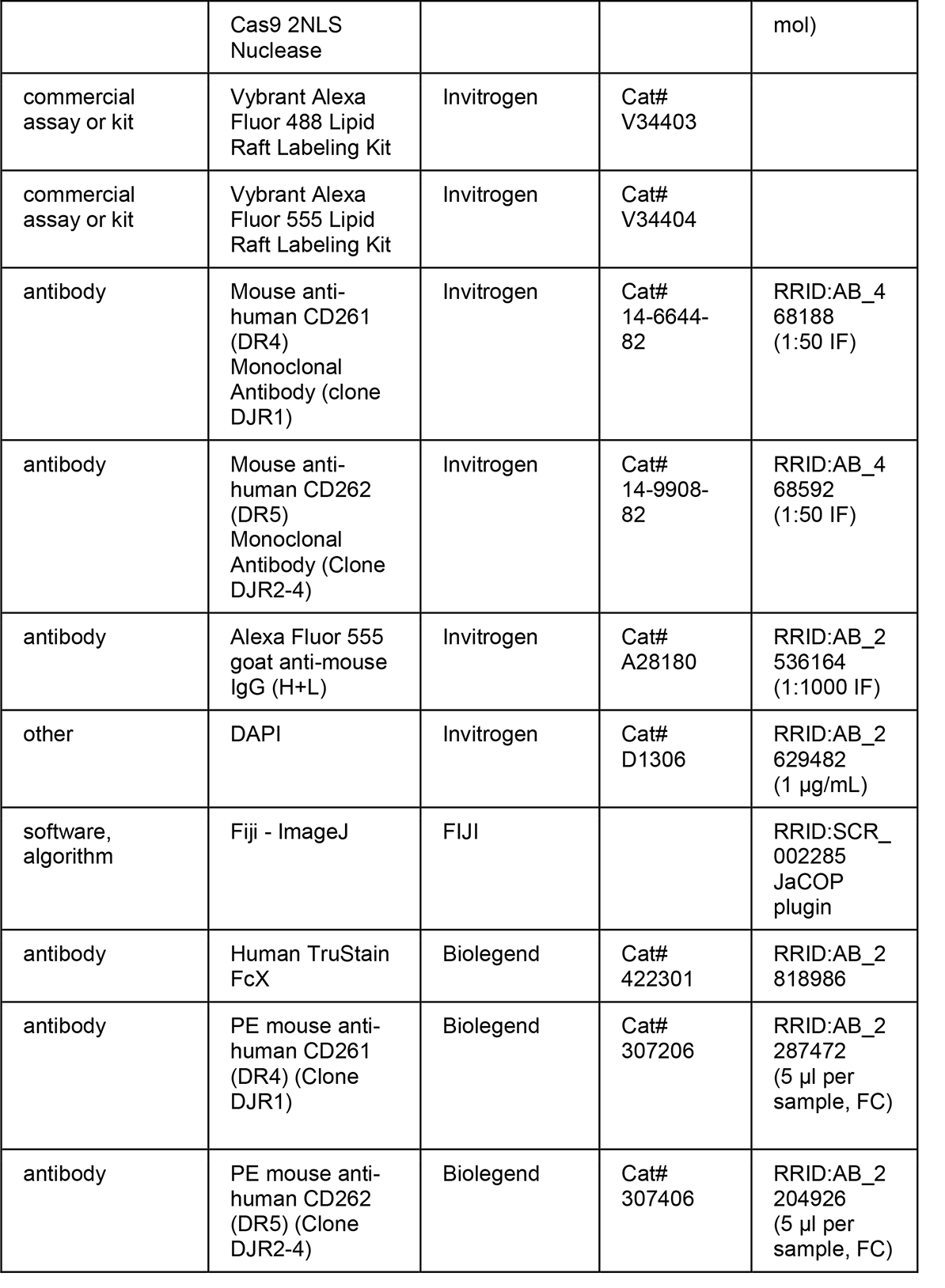
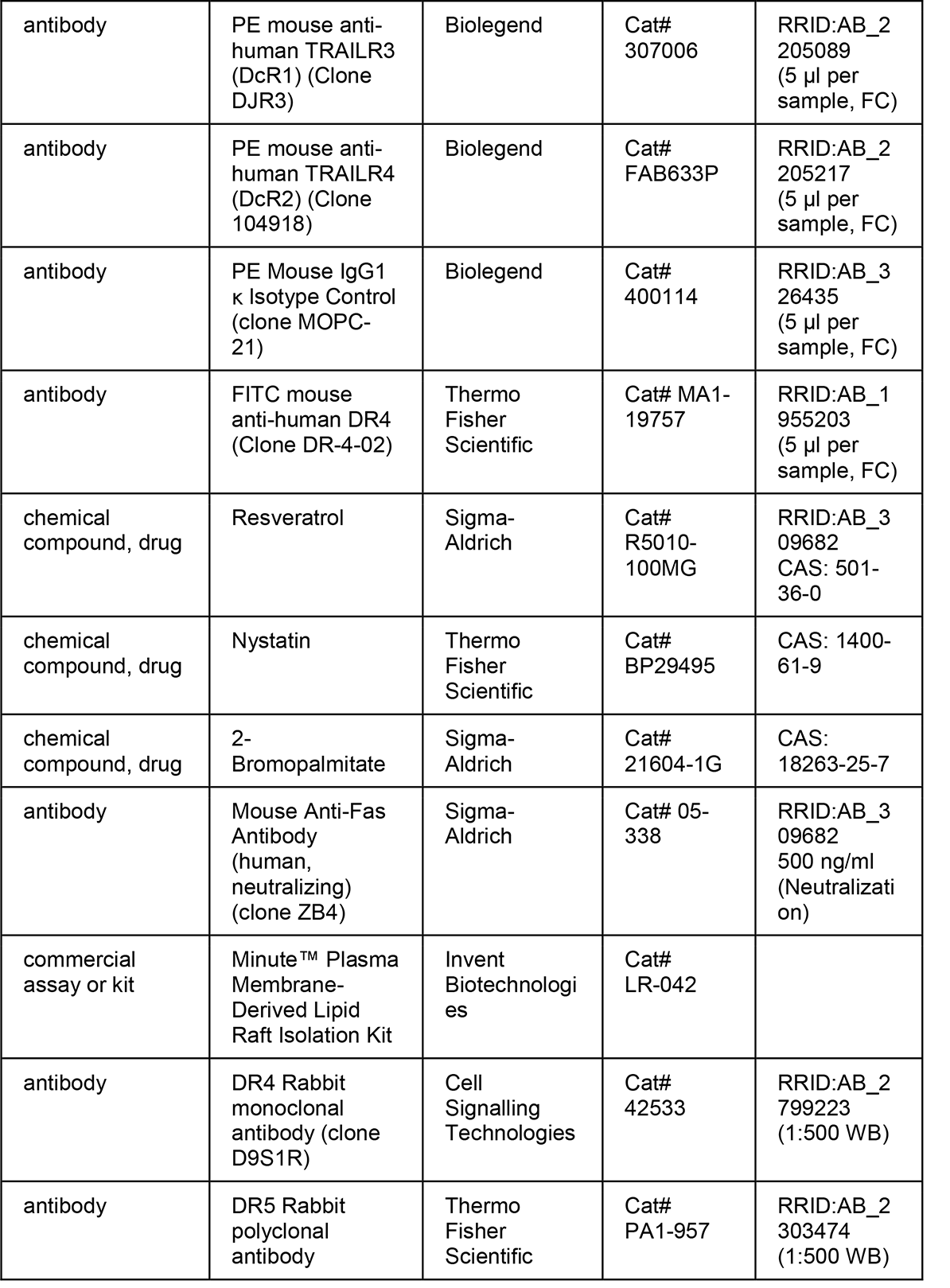
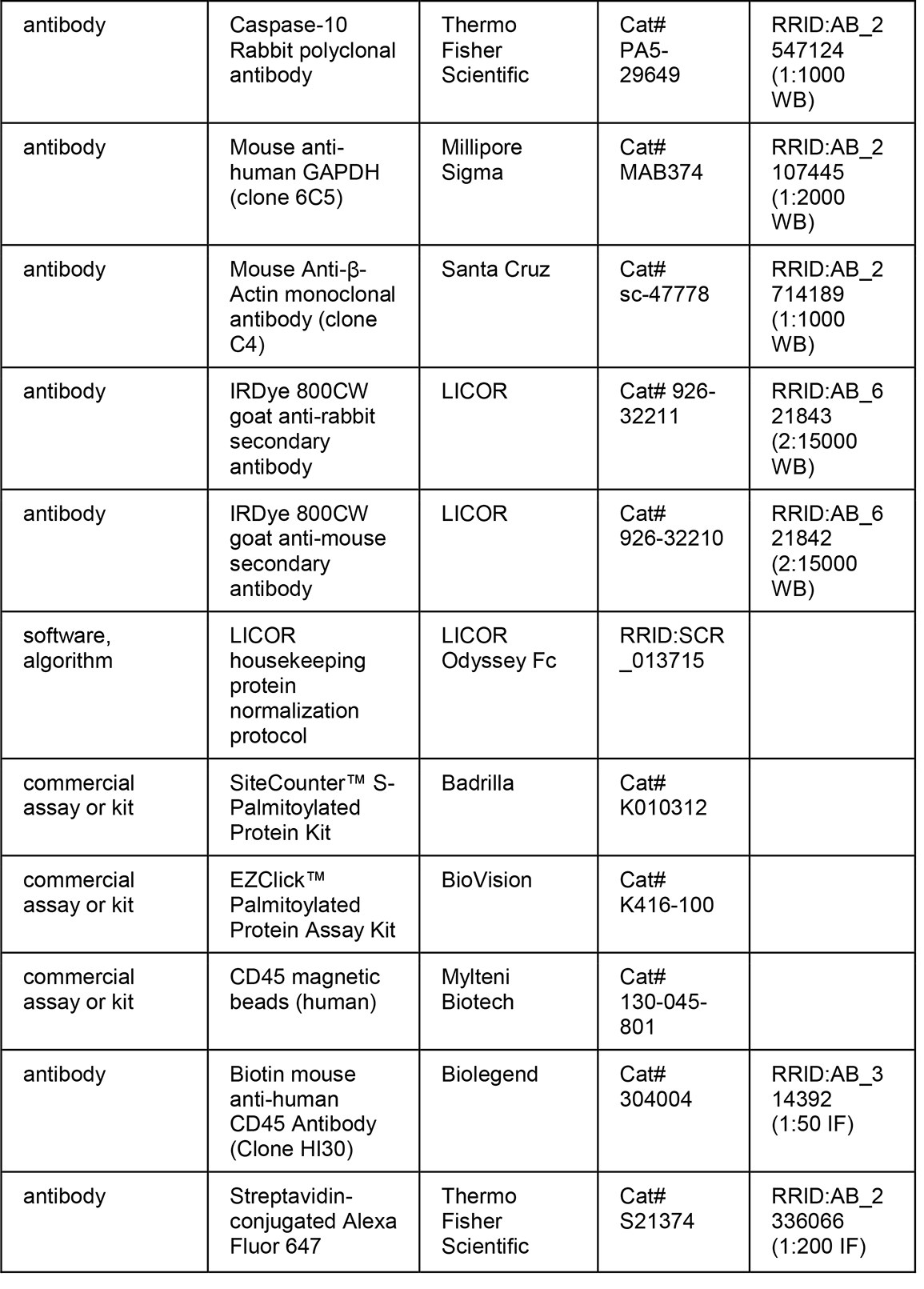
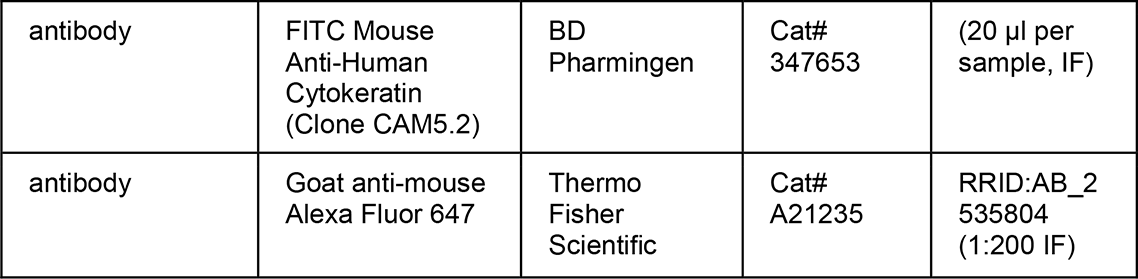

### Cell Culture

Colorectal cancer cell lines SW620 (ATCC, #CCL-227), SW480 (ATCC, #CCL-228), HCT116 (ATCC, #CCL-247), and HT29 (ATCC, #HTB-38), were purchased from American Type Culture Collection. SW620 and SW480 cells were cultured in Leibovitz’s L-15 cell culture medium (Gibco). HCT116 and HT29 cells were cultured in McCoy’s 5A cell culture medium (Gibco). Media was supplemented with 10% (v/v) fetal bovine serum and 1% (v/v) PenStrep, all purchased from Invitrogen. SW480 OxR, HCT116 OxR and HT29 OxR cells were obtained from MD Anderson Cancer Center Characterized Cell Line Core, supplied and generated by the Dr. Lee Ellis laboratory. SW620 OxR cells were obtained from Dr. Mika Hosokawa at Kobe Pharmaceutical University in Japan. Oxaliplatin-resistant derivative cell lines were cultured in the same medium as their parental counterparts. To prevent phenotypic drift of OxR lines, cells were used within 6 passages from the time they were received. To prevent chemotherapy-induced cytotoxicity in downstream experiments, oxaliplatin was not supplemented in OxR cell culture media. All cell lines were maintained in a humidified incubation chamber at 37°C and 5% CO_2_. All cell lines were screened for mycoplasma contamination and tested negative.

### MTT Assay

SW620, SW620 OxR, HCT116, HCT116 OxR, HT29, HT29 OxR, SW480 and SW480 OxR cell lines were plated into tissue culture 96-well black-walled plates, at a concentration of 3,000 cells/well and incubated overnight at 37°C. A 10mM stock oxaliplatin suspension was created by dissolving oxaliplatin (MedChemExpress) in molecular grade water via sonication and heating. Cell culture media was replaced with oxaliplatin treatments ranging from 0-1000µM and incubated for 72 hr. Following treatments, an MTT assay (Abcam) was carried out according to the manufacturer’s protocol. The plates were then read using a plate reader (BioTek µQuant) at 590nm absorbance using gen5 software. Control wells containing the MTT solution without cells were used for background subtraction.

### Transwell Assay

Transwell inserts (6-well with 8 µm pores) (Greiner Bio-one) were evenly coated with 75 µL of a 1 mg/mL collagen solution composed of 3 mg/mL rat tail collagen (Gibco), serum free media, and 0.2% 1N NaOH for crosslinking. Inserts were incubated for 20 min at 37°C. After crosslinking, 2.5 mL of 10% FBS media was added to the bottom of the well plate while the top was filled with 1 mL of serum free media until cells were ready for seeding. Parental and oxaliplatin resistant SW480, SW620, HT29, HCT116 cells were seeded in the collagen-coated inserts at a concentration of 200,000 cells/mL in serum-free media. The transwell inserts were replaced with new serum-free media after two days. On day four, the number of cells that had migrated into the bottom plate were counted using a Thermo Fisher Countess II Automated Cell Counter.

### Annexin-V/PI Apoptosis Assay

Parental and OxR cell lines were plated at 100,000 cells/well onto 24-well plates and incubated overnight at 37°C. Wells were treated in triplicate with soluble human TRAIL (PeproTech), or treated with the anti-DR4 agonist antibody Mapatumumab (Creative Biolabs, clone mAY4) and incubated for 24 hr. All cells were collected by recovering the supernatant and lifting the remaining adhered cells using 0.25% Trypsin-EDTA (Gibco). Cells were washed thoroughly with HBSS with calcium and magnesium (Gibco). Cells were incubated for 15 min with FITC-conjugated Annexin-V and propidium iodide (PI) (BD Pharmingen) at room temperature (RT) in the absence of light and immediately analyzed using a Guava easyCyte 12HT benchtop flow cytometer (MilliporeSigma). Viable cells were identified as being negative for both Annexin-V and PI, early apoptotic cells as positive for Annexin-V only, late apoptotic cells were positive for both Annexin-V and PI, and necrotic cells were positive for PI only. Flow cytometry plots were analyzed using FlowJo v10.7.1 software. Control samples included: unstained negative control with no Annexin-V/PI to adjust for background and autofluorescence, and Annexin V only and PI only samples for gating apoptotic and necrotic populations.

The change in cell viability in response to TRAIL treatments between parental and OxR cells for each of the four CRC cell lines was calculated using the following TRAIL Sensitization equation:

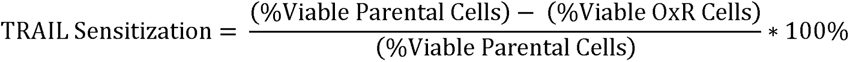

where the percentage of viable cells was normalized to untreated controls for each trial. TRAIL sensitization was calculated for each concentration of TRAIL, where the “Maximum TRAIL Sensitization” was the highest sensitization observed among all concentrations. Since this sensitization equation is based on a percent reduction formula, small changes in viability can yield large TRAIL sensitizations when cell viability is low. To account for this, both cell viability and TRAIL sensitization are reported to provide a complete perspective on treatment responses between cell lines.

### JC-1 (Mitochondrial Membrane Potential) Assay

SW620 and HCT116 cells (parental and OxR) were plated at 100,000 cells/well onto 24-well plates and incubated overnight. Cells were treated in triplicate with TRAIL for 24 hr. Following treatment, cells were collected, washed thoroughly with HBSS without calcium and magnesium, and incubated for 15 min with JC-1 dye (Abcam) in accordance with the manufacturer’s protocol. JC-1 fluorescence was assessed via flow cytometry. Cells with healthy mitochondria were identified as having higher red fluorescence while those with depolarized mitochondria had lower red JC-1 fluorescence.

### RT-PCR Profiler Array

2x10^6^ SW620 and HCT116 (parental and OxR) cells were seeded into a 100 mm diameter cell culture dish for 24 hr. Cells were lifted using a cell scraper and washed with HBSS with calcium and magnesium. RNA was isolated using the RNeasy Plus Mini Kit (Qiagen) according to the manufacturer’s protocol. RNA yield following isolation was determined using a UV5Nano spectrophotometer (Mettler Toledo). cDNA synthesis was completed using the RT^2^ First Strand Kit (Qiagen, 330404) using 0.5μg RNA per sample. RNA expression of 84 apoptotic genes was analyzed using the RT2 Profil^μ^er PCR Human Apoptosis Array (Qiagen, PAHS-012Z). Arrays were prepared according to the manufacturer’s protocols applied to the prepared cDNA samples. Profiler array plates were run on a CFX96 Touch Real Time PCR (Bio-Rad) using the following protocol: 1 cycle for 10 min at 95°C, 40 cycles of 95 °C for 15 s followed by 60°C for 60 s at a rate of 1°C/s. Melt curves were generated immedi^L^ately following the PCR protocol. Cycle threshold (Ct) values were calculated using CFX Maestro Software (Bio-Rad). Data analysis was completed using the GeneGlobe Data Analysis Center (Qiagen). Volcano plots were generated in GraphPad Prism using calculated fold changes in gene expression between OxR and parental cells and their corresponding p-values.

### CRISPR-Cas9 Knockout

Knockout of the CASP10 gene in SW620 cells was completed using the Gene Knockout Kit v2 – Human CASP10 kit with Cas9 2NLS Nuclease (Synthego). Ribonucleoprotein (RNP) complexes were made at a 9:1 ratio of sgRNA:Cas9 (90pmol:10pmol) in Gene Pulser® Electroporation Buffer (Bio-Rad, 1652677) and incubated for 10 min at RT. Cas9 control samples consisted of 10 pmol Cas9 with no sgRNA. RNP complexes were added to 200,000 cells in 200µL electroporation buffer (0.2 cm cuvette) and electroporated via the Gene Pulser Xcell™ Electroporation System (Bio-Rad) using exponential decay pulses (145V, 500µF, 1000ohm). Cells were immediately cultured in 12-well plates and allowed to recover for 7 days before measuring knockout efficiency.

### Confocal Microscopy and Image Analysis

Parental and OxR cells were seeded onto polystyrene cell culture slides (Thermo Fisher Scientific). Cells were allowed to grow for 48 hr at 37°C. In samples treated with nystatin or resveratrol, cells were plated for 24 hr then treated for 24 hr before staining. Cells were washed and lipid rafts were stained using the Vybrant Alexa Fluor 488 Lipid Raft Labeling Kit (Invitrogen, V34403) according to the manufacturer’s protocol. Briefly, cells were incubated with Alexa488-conjugated cholera toxin subunit B (CT-B) followed by an anti-CT-B antibody to crosslink CT-B labeled rafts. Slides were fixed for 15 min with 4% paraformaldehyde (PFA) (Electron Microscopy Sciences) in PBS (Gibco) and then permeabilized using 1% Triton X-100 (MilliporeSigma) in PBS at RT. Slides were blocked for 2 hr with 5% goat serum (Thermo Fisher Scientific) and 5% bovine serum albumin (BSA, Sigma) in HBSS. Primary staining was done overnight at 4°C with either DR4 monoclonal antibody (Invitrogen, Clone DJR1) or DR5

monoclonal antibody (Invitrogen, Clone DJR2-4) in the blocking serum at a ratio of 1:50. Secondary staining was carried out with Alexa Fluor 555 goat anti-mouse IgG (H+L) (Invitrogen, A28180) for 30 min at RT (1:1000). Slides were stained with DAPI (Invitrogen, D1306) for 30 min at RT in the blocking solution at 1:1000. Washes were done twice between each step for 5 min each using 0.02% Tween20 in PBS. Slides were assembled using 10 μl of Vectrashield antifade mounting media (Vector Laboratories). Confocal imaging was performed using an LSM 880 (Carl Zeiss) with a 63x/1.40 Plan-Apochromat Oil, WD = 0.19 mm objective. At least five images were taken per sample

Image analysis was performed in FIJI using a macro to automate quantification of raft and DR contents per cell. Briefly, all images were adjusted for background using the same thresholding specifications. The “analyze particles” feature was used to quantify the total area of lipid rafts and DR per outlined cell. Colocalization events were quantified by creating binary masks of DR and lipid raft events. For each gated cell, the lipid raft and DR binary masks were multiplied to create a binary projection of colocalized events. Raw integrated density and cell area (ROI area) were also measured. Cells with areas outside of three times the standard deviation from the mean were considered outliers and not included in the analysis. Colocalization analysis was also performed using the JACoP plugin in FIJI (58). The Manders’ Correlation Coefficient was calculated as the fraction of lipid raft colocalized DR4.

### Flow Cytometry

#### Surface DR Expression

Parental and OxR cell lines were cultured to 70% confluency upon collection and split into 250,000 cells per sample. Cells were fixed in 4% PFA in HBSS for 15 min at RT, then blocked in a 100 μL 1% BSA solution for 30 min at 4°C, with 2X HBSS washes between each step. Cells suspensions of 100 μL were incubated for 15 min at RT with 2 μL Human TruStain FcX (Biolegend, 422301) to prevent nonspecific Fc receptor binding. Samples were immediately stained with 5μL of either PE anti-human CD261 (DR4) (Biolegend, Clone DJR1), PE antihuman CD262 (DR5) (Biolegend, Clone DJR2-4), PE anti-human TRAILR3 (DcR1) (Biolegend, Clone DJR3), PE anti-human TRAILR4 (DcR2) (R&DSystems, Clone 104918) or PE Mouse IgG1 κ Isotype Control (Biolegend, Clone MOPC-21) for 30 min at 4°C. Samples were washed twice with HBSS and analyzed using a Guava easyCyte flow cytometer. A chi-squared test was performed using FlowJo v10.7.1, where significance in histogram distribution was confirmed if T(x) between parental and OxR stained samples was greater than T(x) between background (unstained) parental and OxR samples (see **Supplementary File 1**).

#### FRET

Cells were prepared as described above, but without fixation or permeabilization. Samples were stained for lipid rafts using the Vybrant Alexa Fluor 555 Lipid Raft Labeling Kit (Invitrogen, V34404). Samples were then stained with 5 µL FITC anti-human DR4 (Thermo Fisher Scientific, Clone DR-4-02) for 30 min at 4°C. Samples were washed twice with HBSS and analyzed using a Guava easyCyte flow cytometer. Donor quenching FRET efficiency was calculated using the following formula:

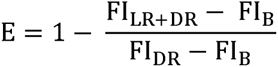

Where E is the FRET efficiency, FI_LR+DR_ is the mean fluorescence intensity of the double stained lipid raft/DR4 sample (acceptor + donor), FI_DR_ is the mean fluorescence intensity of the DR4 only stain (donor only) and FI_B_ is the flurescence intensity of an unstained sample (background). Fluorescence intensit^F^y as recorded in the donor (FITC) channel.

### TRAIL Combination Treatments

#### Resveratrol

Parental SW620 and HCT116 cells were plated at 100,000 cells/well onto 24-well plates and incubated overnight at 37°C. Cells were treated with 70 µM resveratrol (Sigma) in combination with 0.1-1000 ng/ml of TRAIL for 24 hr. Following treatment, cells were collected for Annexin-V/PI apoptosis assay. TRAIL sensitization was calculated using the following equation:

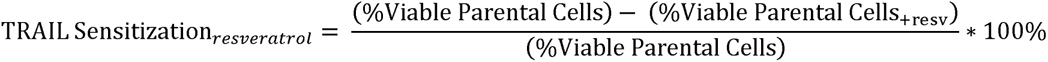

where TRAIL + resv treatments were normalized to resveratrol treatment in the absence of TRAIL to account for any resveratrol-associated cytotoxicity.

#### Nystatin

SW620 OxR and HCT116 OxR cells were plated at 100,000 cells/well onto 24-well plates and incubated overnight at 37°C. Cells were treated with 5 µM nystatin (Thermo Fisher Scientific) in combination with 0.1-1000 ng/ml of TRAIL. Following treatment, cells were collected for Annexin-V/PI apoptosis assay. TRAIL sensitization was calculated using the following equation:

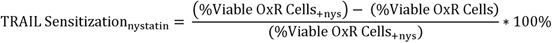

where TRAIL + nys treatments were normalized to nystatin treatment in the absence of TRAIL to account for any nystatin associated cytotoxicity.

#### 2-Bromopalmitate

SW620 parental and OxR cells were plated at 100,000 cells/well onto 24-well plates and incubated overnight at 37°C. Cells were treated with 3.5 µM 2BP (Millipore Sigma) in combination with 0.1-1000 ng/ml of TRAIL. Following treatment, cells were collected for Annexin-V/PI apoptosis assay. TRAIL sensitization was calculated as described above.

#### Anti-Fas (ZB4)

SW620 OxR cells were plated at 100,000 cells/well onto 24-well plates and incubated overnight at 37°C. Cells were treated with 500 ng/ml human anti-Fas (Millipore Sigma, Clone ZB4) with and without 1000 ng/ml of TRAIL. Following treatment, cells were collected for Annexin-V/PI apoptosis assay. TRAIL sensitization was calculated as described above.

### Western Blot

Lipid rafts were isolated according to manufacturer’s protocol using the Minute™ Plasma Membrane-Derived Lipid Raft Isolation Kit (Invent Biotech, LR-042). Cell lysates and lipid raft protein isolates were prepared by sonication in 4x Laemmli sample buffer (Bio-Rad, 1610747) and then loaded into 10% SDS-polyacrylamide gels for electrophoresis. Protein transfer onto a PVDF membrane was carried out overnight, and then blocked with Intercept (TBS) Blocking Buffer (LICOR, 927-60001) at RT for an hour. Primary antibody incubation occurred overnight at 4°C for DR4 (Cell Signaling Technology, 42533) and DR5 (Thermo Fisher Scientific, PA1-957) at 1:500 dilution and caspase-10 (Thermo Fisher Scientific, PA5-29649) at a 1:1000 dilution in LICOR buffer. Cell lysate protein bands were normalized to GAPDH (EMD Millipore, MAB347) at 1:2000 dilution, while lipid raft isolates were normalized to β-actin (Santa Cruz, 47778) at 1:1000 dilution in LICOR blocking buffer. Western blots were quantified using the Licor Odyssey Fc with IRDye 800CW goat anti-rabbit secondary antibody (LICOR, 926-32211) and IRDye 800CW goat anti-mouse secondary antibody (LICOR, 926-32210) at a dilution of 2:15000. Quantification was done following the LICOR housekeeping protein normalization protocol.

### Palmitoylation Assay

Cells were grown to 70% confluency in a 100 mm tissue culture dish. Palmitoylation of DR4 was measured using the SiteCounter™ S-Palmitoylated Protein Kit (Badrilla, K010312) according to the manufacturer’s protocol. Input fraction controls (IFC) were obtained prior to thioester cleavage. Acyl preservation negative controls (APC-) were obtained by using an acyl preserving reagent instead of mass-tag conjugation. Western blots were run for DR4 following the “Western Blot” protocol described above. The percentage of DR4 palmitoylation was calculated by dividing the total intensity of all palmitoylated bands (mass tag) divided by the average intensity of the IFC and APC(-) bands for that sample.

To measure the amount of total palmitoylated protein, cells were cultured in 96-well plates at a concentration of 20,000 cells/well. The EZClick™ Palmitoylated Protein Assay Kit (BioVision, K416-100) was used in accordance with the manufacturer’s protocol. Cells were incubated overnight with either 1x EZClick™ Palmitic Acid label in media or culture media with no label (background control). Cells were recovered and stained using EZClick™ Fluorescent Azide, then analyzed via flow cytometry for shifts in FL2-H intensity. Median fluorescence intensity (MFI) was calculated by subtracting the background intensity from each sample (Palmitic Acid label [-]/ Fluorescent Azide [**+**]).

### Patient Blood Samples

Peripheral whole blood samples of 10 mL were collected from 13 metastatic CRC patients after informed consent. Patient criteria for this study included the following: presenting with metastatic CRC at the time of blood draw and undergone (or undergoing) oxaliplatin-containing chemotherapy (i.e. FOLFOX). Additionally, 5 patients had samples collected through their respective chemotherapy regimens. De-identified blood samples were transported from the Guthrie Clinic to Vanderbilt University and processed within 24 hr. Blood samples were split for treatment (8 ml) and death receptor/lipid raft staining (1-2 ml).

#### Ex-vivo treatment of colorectal cancer patient blood samples

For the treated samples, 2 mL of blood were treated with either 40 µL of control liposomes, 40µl TRAIL/E-selectin conjugated liposomes (290 ng/mL of TRAIL), 6 µL (290 ng/mL) of soluble TRAIL or 2 µL (5 µM) of oxaliplatin. Liposomes were synthesized using a thin film hydration method followed by extrusion and his-tag conjugation as described previously (44). The aliquots were sheared for 4 hr in a cone-and-plate viscometer (Brookfield LVDVII) at a shear rate of 120 s^-1^. Prior to incubation, the cone-and-plate viscometers were blocked using 5% BSA for 30 min. After 4 hr, the blood aliquots were washed from the viscometer’s spindle and cup by using twice the volume of HBSS without calcium and magnesium. Blood aliquots were placed over twice the volume of Ficoll (GE Healthcare) to separate out mononuclear cells within the buffy coat. CTCs were enriched using a negative selection kit with CD45 magnetic beads (Mylteni Biotech, 130-045-801) following the manufacturer’s protocol (43).

The resulting isolated CTCs were placed in cell culture overnight using RPMI media supplemented with 10% FBS. After 1 day in culture, the cells were recovered from the tissue culture plate and stained with 100 µL of propidium iodide for 15 min. Cells were washed, fixed with 4% PFA and cytospun onto microscope slides using a Cytospin 3 (Shandon). Samples were then permeabilized and blocked with 100 µL of 0.25% Triton-X (Sigma) for 15 min and 100 µL of blocking solution (5% BSA and 5% goat serum) for 1 hr, respectively. Cells were stained with anti-CD45 conjugated with biotin (Biolegend, Clone HI30) for 45 min at 1:50 dilution. Finally, cells were stained with 100 µL of streptavidin-conjugated Alexa Fluor 647 (Thermo Fisher Scientific, S21374) at 1:200 dilution and 20 µl per sample of anti-cytokeratin conjugated with FITC (BD Pharmingen, Clone CAM5.2) for 45 min. Cells were washed 3X after each staining incubation using 200 µL of 0.02% Tween20 in PBS. Cells were stained with 10 µL of DAPI mounting media (Vector Laboratories), covered with a coverslip (No. 1.5, VWR), and sealed with nail polish.

Five images per sample were taken at random locations using an LSM 710 (Carl Zeiss) with a 20x/0.8 objective. The cell number in the sample was scaled up by multiplying by the relative area (slide area/frame area). Viable tumor cells were identified using the following criteria: (i) positive for DAPI, (ii) negative for CD45, (iii) positive for cytokeratin and (iv) negative for propidium iodide.

#### Staining of lipid raft and death receptors in primary CTCs

CTCs from the remainder of the patient blood were isolated and cytospun onto slides as described above. Death receptors and lipid rafts were stained and analyzed as detailed above in “Confocal Microscopy and Image Analysis”. Lipid rafts were stained using the Vybrant Alexa Fluor 555 Lipid Raft Labeling Kit (Invitrogen, V34404) after CTCs were spun onto slides. Secondary staining for DR4 and DR5 was completed using goat anti-mouse Alexa Fluor 647 (Thermo Fisher Scientific, A21235) at a 1:200 dilution. Cells were also stained with FITC-conjugated cytokeratin, as described above, to positively identify CTCs for analysis.

### Statistical Analysis

Data sets were plotted and analyzed using GraphPad Prism 9. When comparing two groups, a symmetric unpaired t-test was used with p < 0.05 considered significant. One-way ANOVA with multiple comparisons was used for multiple groups with p < 0.05 considered significant. At least three independent biological replicates were used for each experiment unless otherwise stated.

## Supplementary

**Figure 1-figure supplement 1.**
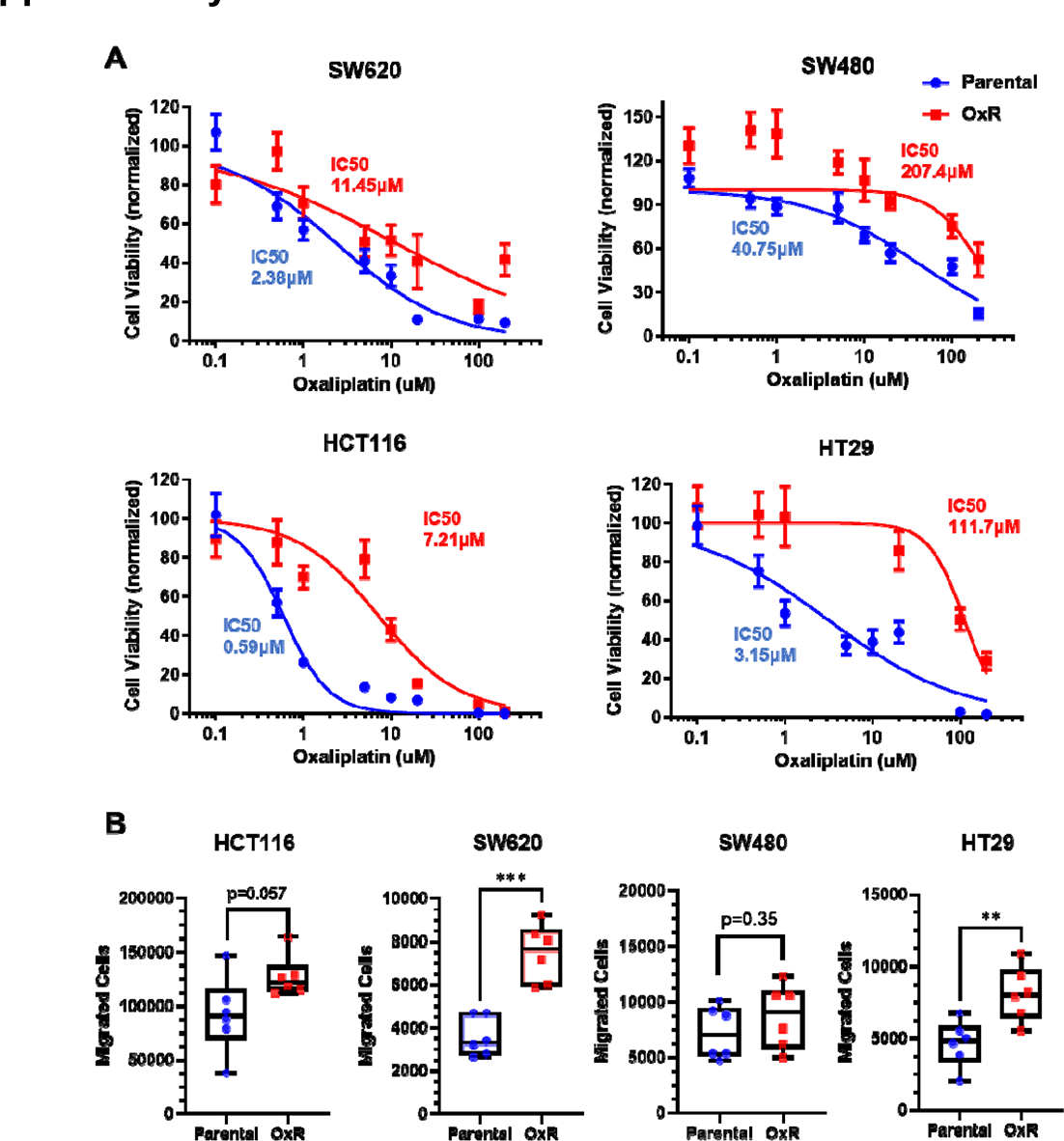
Oxaliplatin-resistant CRC cells retain their increasingly chemoresistant and invasive phenotypes in culture. **(A)** SW620, SW480, HCT116, and HT29 cells were treated with various concentrations of oxaliplatin for 72 hr and cell viability was measured using an MTT assay. IC50 values were calculated using a variable slope four parameter nonlinear regression. Data are presented as mean±SEM. N=2 (n=12). **(B)** Counts of successfully invasive cells after a four-day Transwell assay with an initial seeding of 200,000 cells. N=2 (n=6). * p<0.05 **p<0.01 (unpaired two-tailed t-test).

**Figure 1-figure supplement 2.**
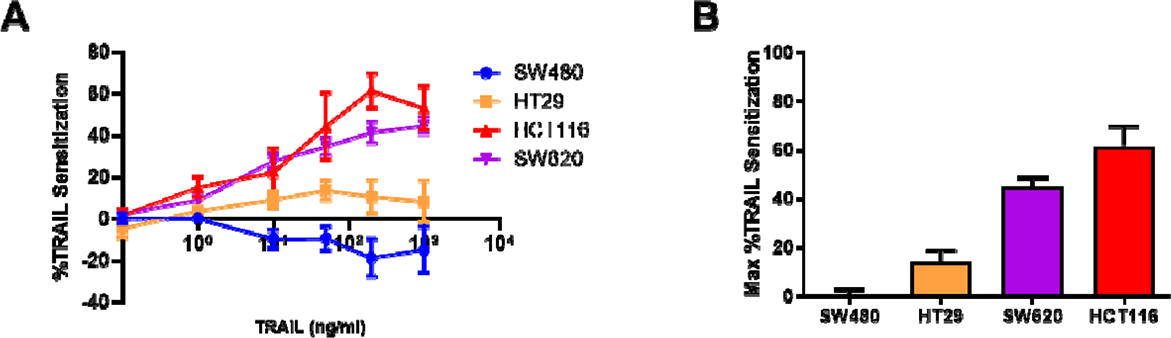
Sensitization of oxaliplatin-resistant CRC cell lines to TRAIL. **(A)** Sensitization of oxaliplatin-resistant SW620, HCT116, HT29 and SW480 cell lines compared to their parental counterparts as a function of TRAIL concentration. **(B)** Maximum TRAIL sensitization for each cell line between the tested concentrations of 0.1-1000 ng/ml. Data are presented as mean±SEM.

**Figure 1-figure supplement 3.**
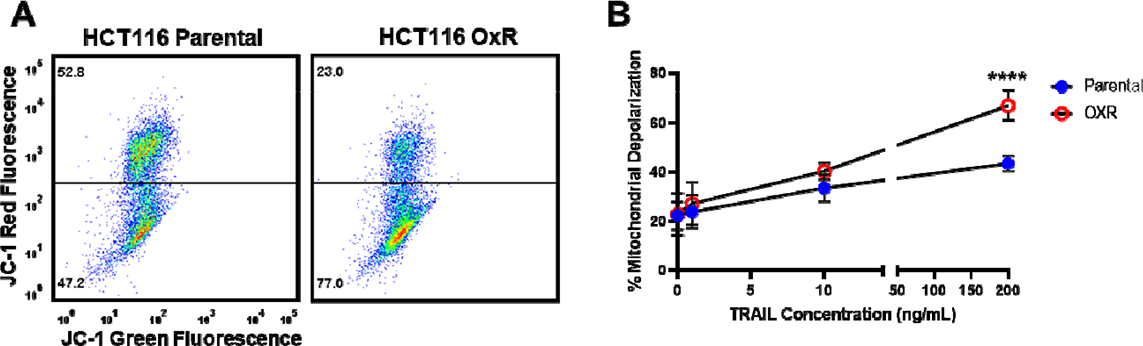
HCT116 OxR cells have increased mitochondrial depolarization and activation of the intrinsic apoptotic pathway when treated with TRAIL. **(A)** Representative flow plots of JC-1 assay after treatment with 200 ng/ml of TRAIL. Mitochondrial depolarization is evidenced by decreased red fluorescence and increased green fluorescence. **(B)** Mitochondrial depolarization as a function of TRAIL concentration for HCT116 parental and OxR cell lines. Data are presented as mean±SD. N=3 (n=9). ****p<0.0001 (unpaired two-tailed t-test).

**Figure 2-figure supplement 1.**
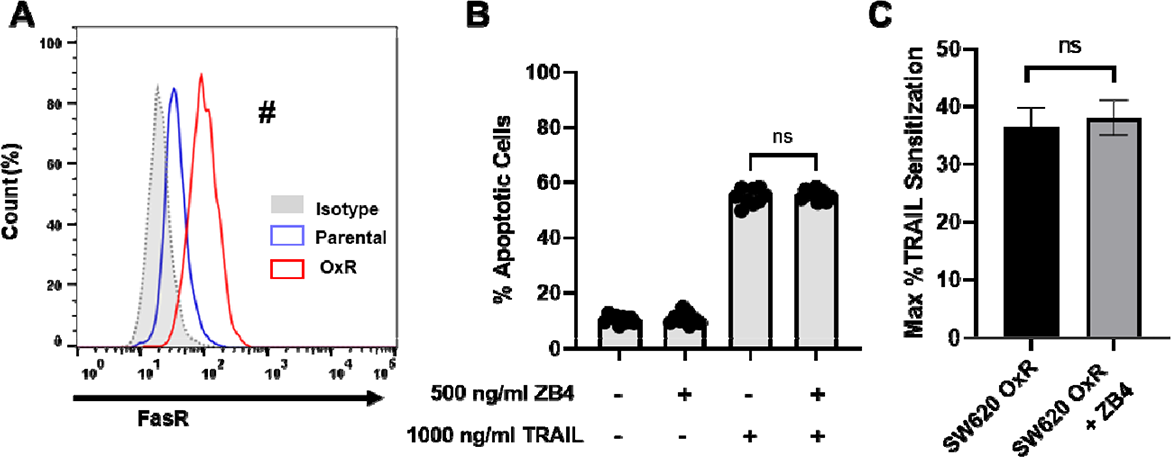
Upregulated FasR in SW620 OxR cells has no effect on TRAIL sensitivity. **(A)** Flow cytometry staining confirms increased surface expression of Fas receptor. ^#^Significant according to a chi-squared test (see Supplementary File 1). **(B)** Percentage of apoptotic SW620 OxR cells after treating with 1000 ng/ml TRAIL and 500 ng/ml of the anti-FasR neutralizing antibody ZB4 (sum of early and late-stage apoptotic cells from Annexin/PI staining). Data are presented as mean+SD. N=3 n=9. Significance was measured using an ordinary one-way ANOVA–Tukey’s multiple comparison test. **(C)** Neutralizing FasR has no effect on TRAIL sensitization. Data are presented as mean+SEM. N=3 (n=9). Significance was measured using an unpaired two-tailed t-test.

**Figure 3-figure supplement 1.**
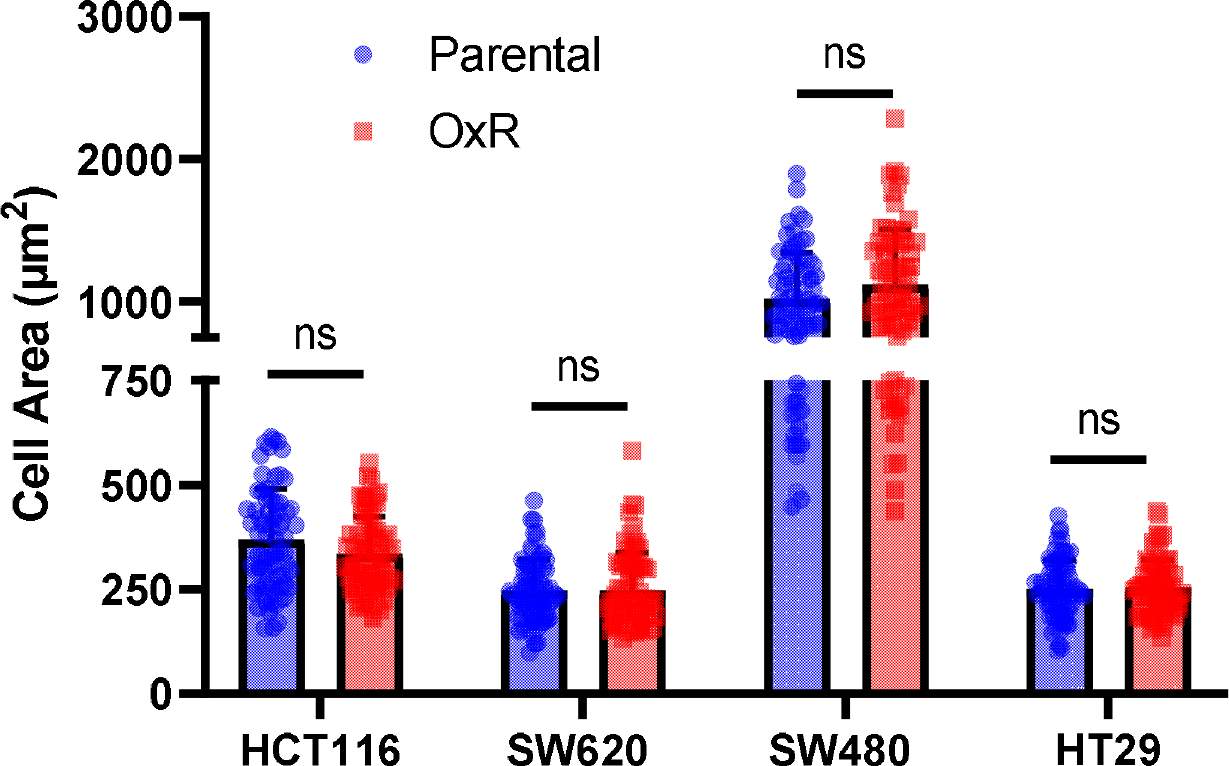
Parental and OxR cell lines have no significant changes in cell area, analyzed from confocal microscopy images. Data are presented as mean+SD. For each cell line, N=75 cells were analyzed. An unpaired two-tailed t-test was used to measure significance between groups.

**Figure 3-figure supplement 2.**
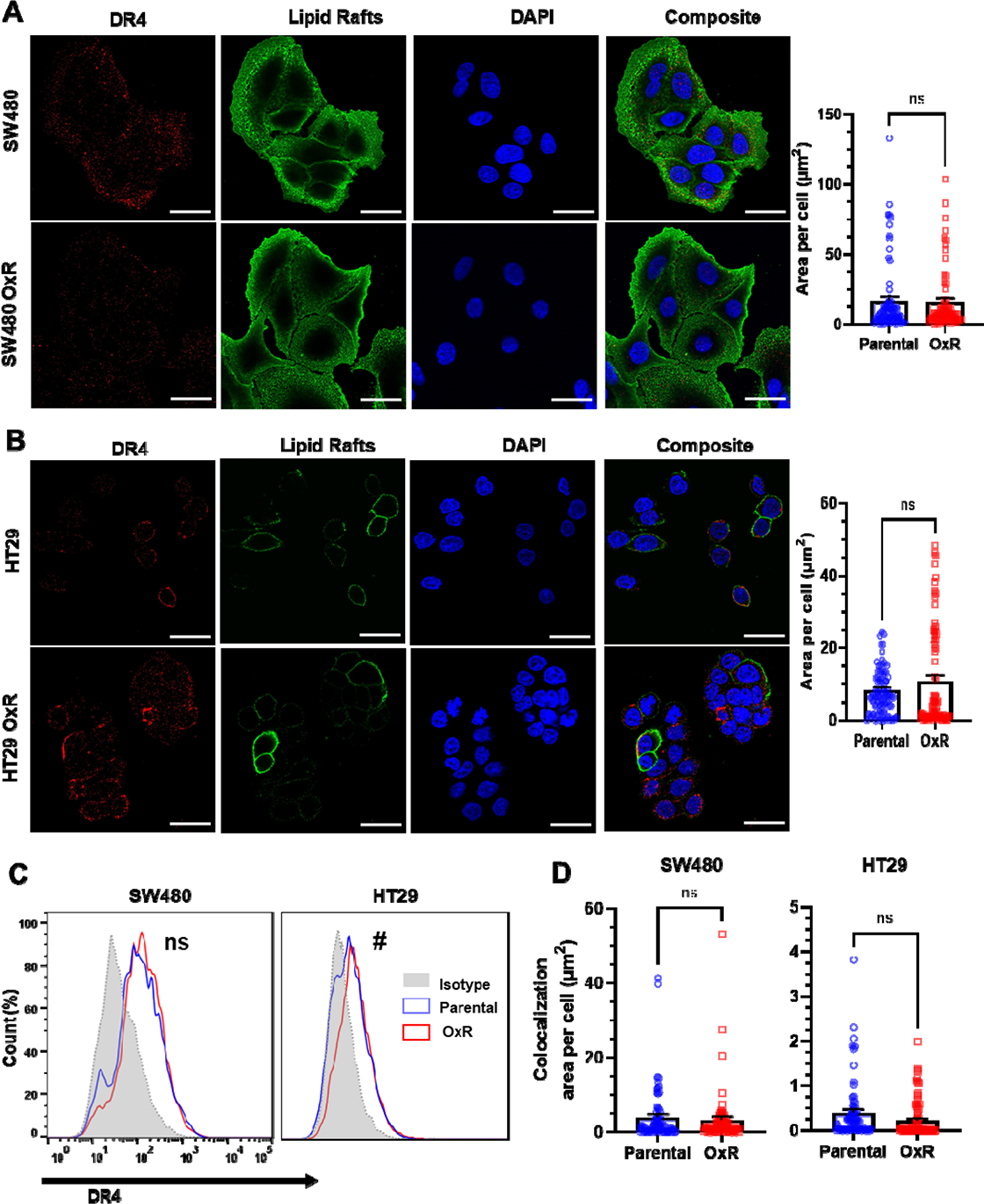
Unsensitized SW480 and HT29 OxR cell lines show no significant changes in DR4 expression or lipid raft colocalization relative to their parental counterparts. **(A-B)** Confocal micrographs and DR4 quantification of SW480 and HT29 cells, respectively. Red channel is DR4, green is lipid rafts and blue is DAPI (nuclei). Scale bar = 30 μm. **(C**) Surface expression of DR4 in non-permeabilized cells analyzed via flow cytometry. Significant according to a chi-squared test (see Supplementary File 1). **(D)** DR4/LR colocalization area per cell for SW480 and HT29 was found to be insignificant between parental and OxR phenotypes. Data are presented as mean+SEM. An unpaired two-tailed t-test was performed for panels A, B and D. For each cell line, N=75 cells were analyzed.

**Figure 3-figure supplement 3.**
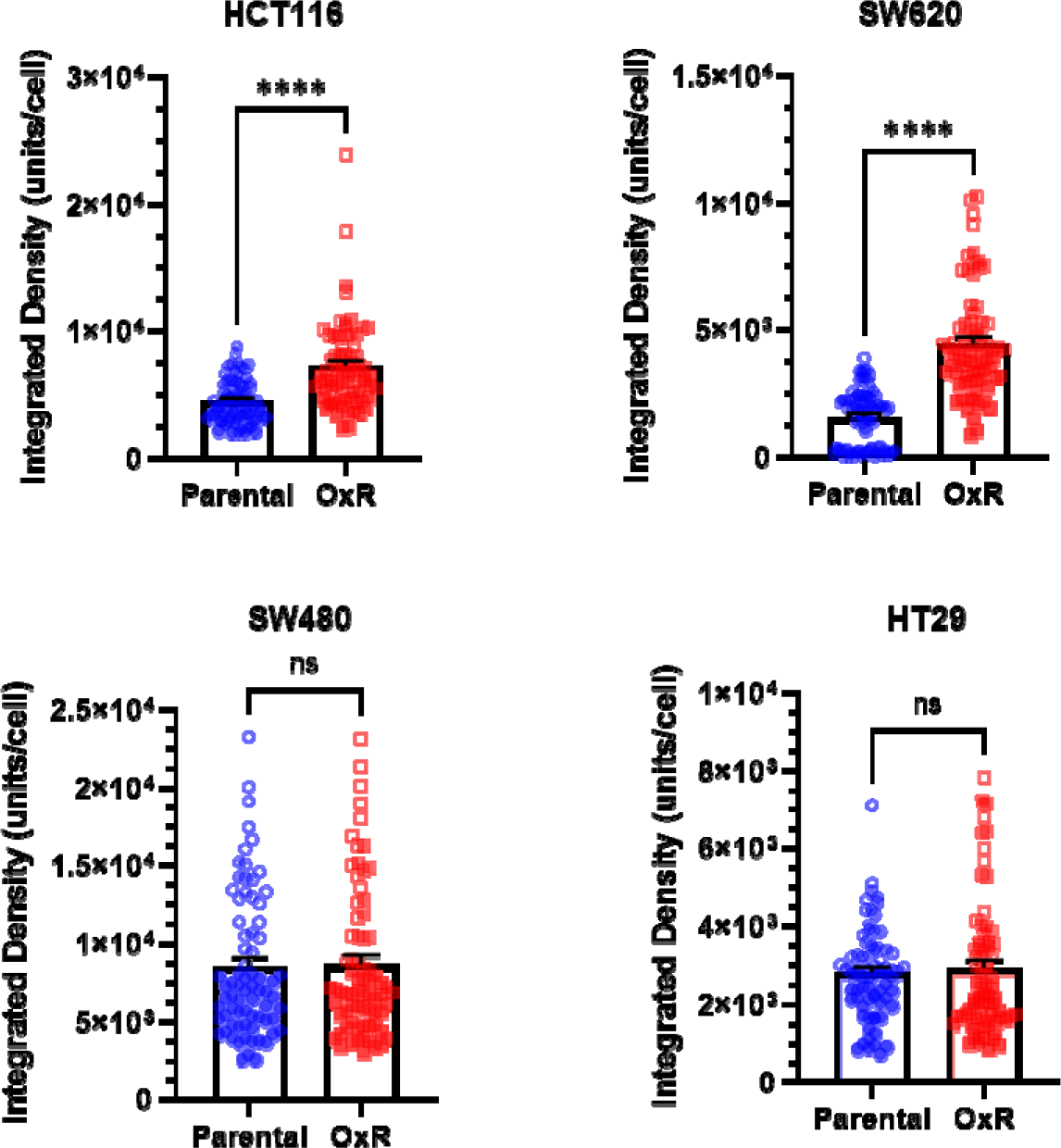
Sensitized oxaliplatin-resistant cell lines have significantly increased DR4 integrated density per cell. Data are presented as mean+SEM. For each cell line, N=75 cells were analyzed. ****p<0.0001 (unpaired two-tailed t-test).

**Figure 3-figure supplement 4.**
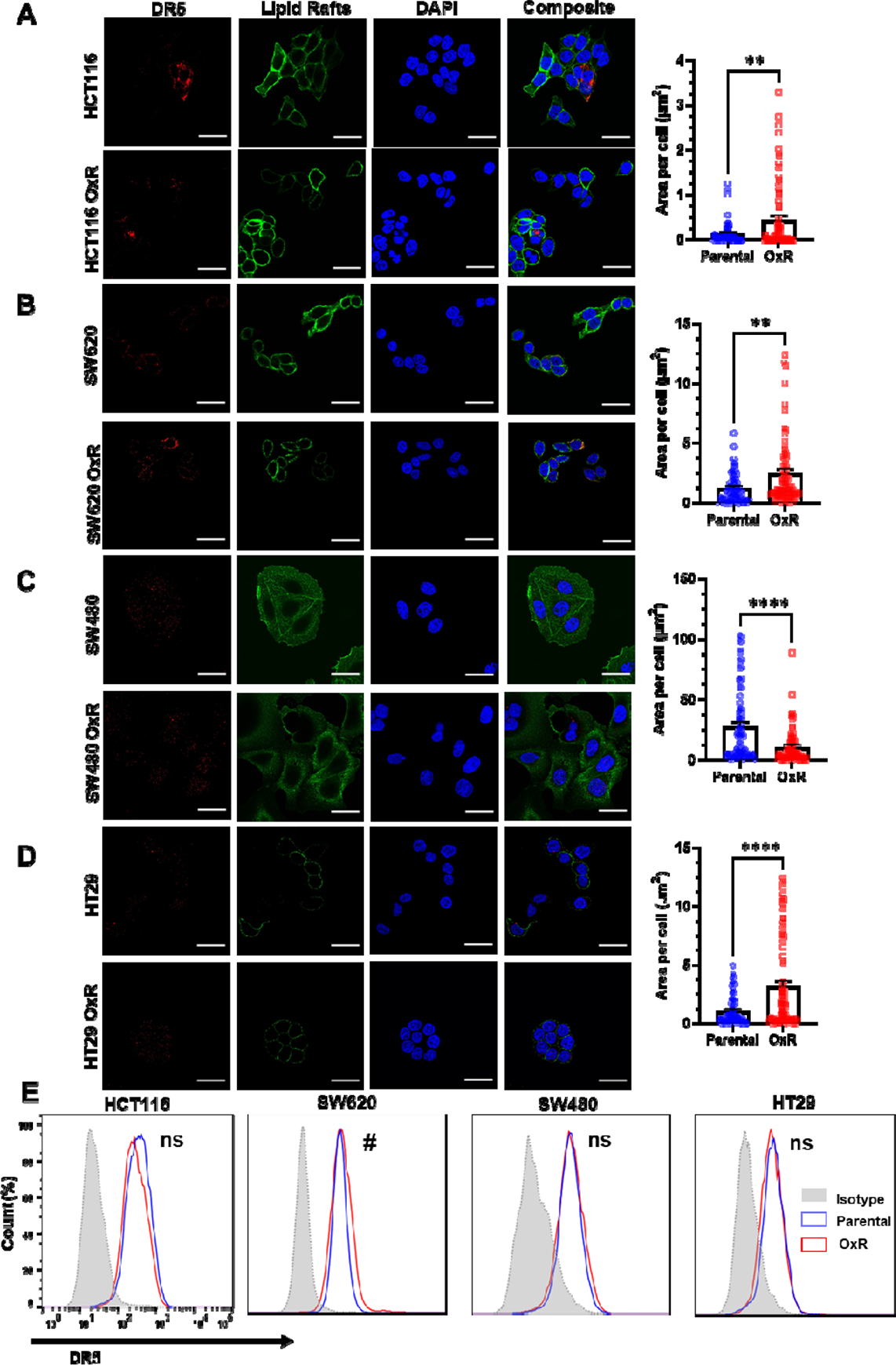
Chemoresistant HCT116, SW620 and HT29 cells have upregulated DR5 while in chemoresistant SW480 cells, DR5 is decreased. **(A-D)** Confocal micrographs and DR5 quantification of HCT116, SW620, SW480 and HT29 cells, respectively. Red channel is death receptor 5, green is lipid rafts and blue is DAPI (nuclei). Scale bar = 30m. ** p<0.01 ****p<0.0001 (unpaired two-tailed t-test). Data are presented as mean+ SEM. For each cell line, N=75 cells were analyzed. **(E**) OxR cells only demonstrate increased surface expression of DR5 in non-permeabilized SW620 cells. ^#^Significant according to a chi-squared test (see Supplementary File 1).

**Figure 3-figure supplement 5.**
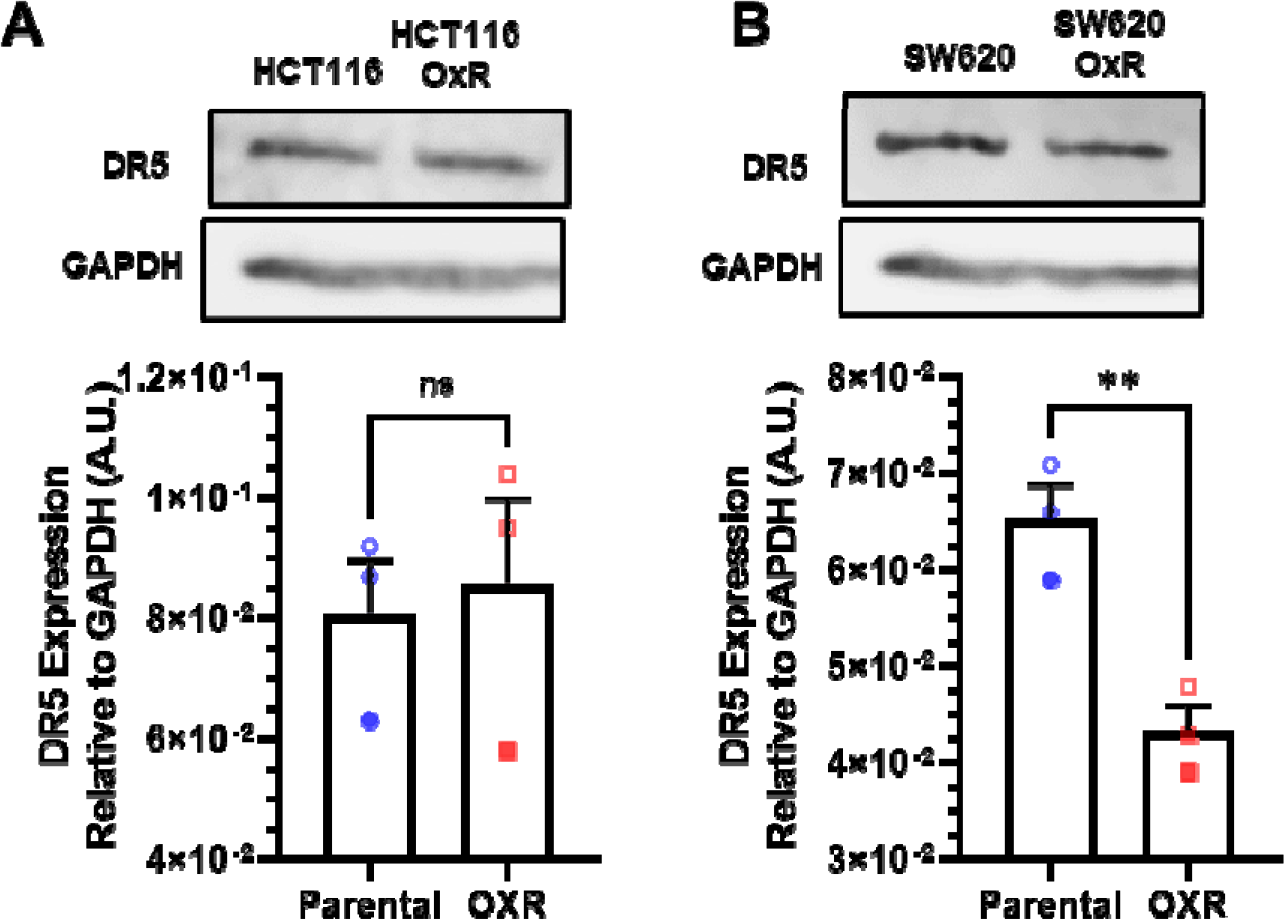
Western blots show TRAIL-sensitized HCT116 (A) and SW620 (B) OxR cells have no increases in DR5. Data are presented as mean+SEM. N=3. ** p<0.01 (unpaired two-tailed t-test).

**Figure 3-figure supplement 6.**
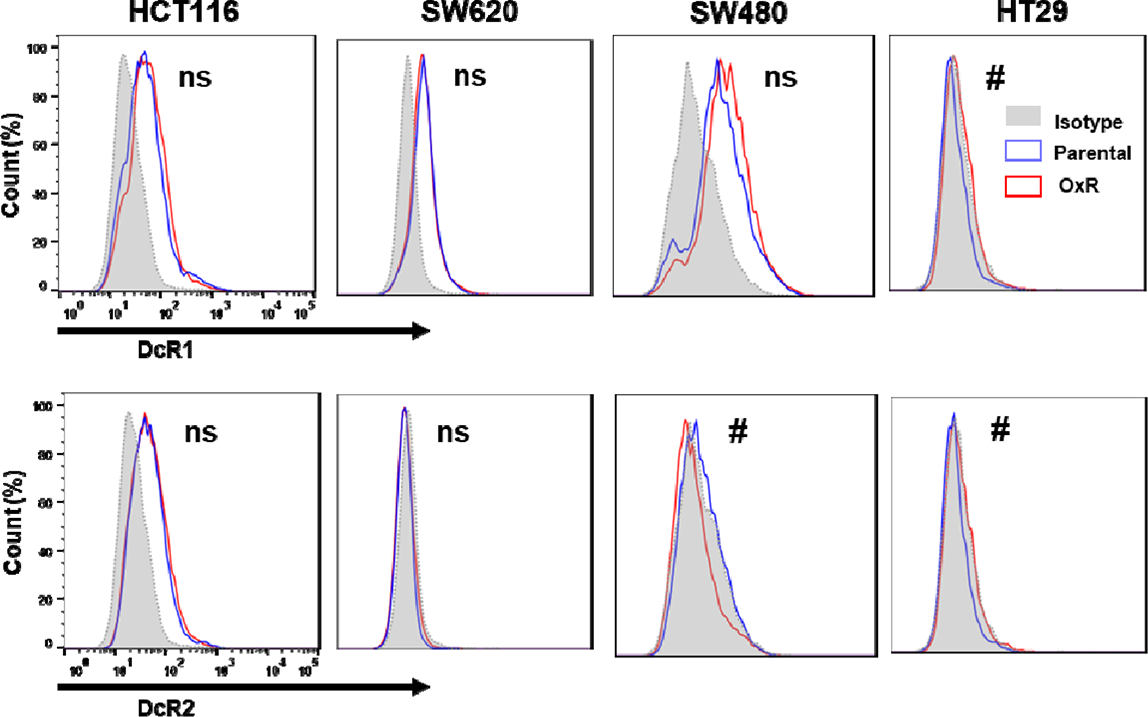
Flow cytometry analysis of the surface expression of decoy death receptors 1 (DcR1) and 2 (DcR2) on nonpermeabilized parental and OxR cell lines. ^#^Significant according to a chi-squared test (see Supplementary File 1).

**Figure 3-figure supplement 7.**
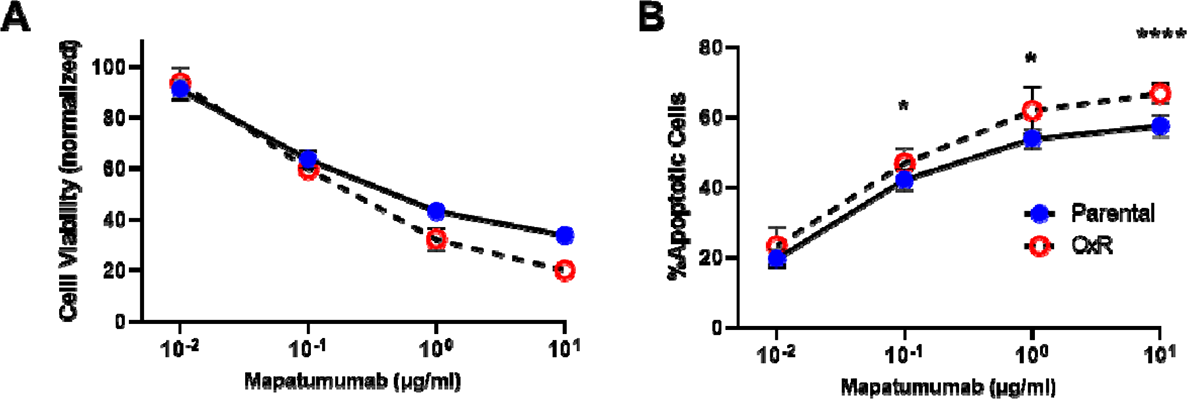
HCT116 OxR cells are increasingly sensitive to DR4 agonist antibody treatment. **(A)** Cell viability of HCT116 cells after 0.01-10 µg/ml Mapatumumab treatment, determined by AnnexinV/PI staining. **(B)** Percentage of apoptotic SW620 cells after Mapatumumab treatment (sum of early and late-stage apoptotic cells from Annexin/PI staining). For all graphs, data are presented as mean+SD. N=3 (n=6). *p<0.05****p<0.0001 (multiple unpaired two-tailed t-tests).

**Figure 4-figure supplement 1.**
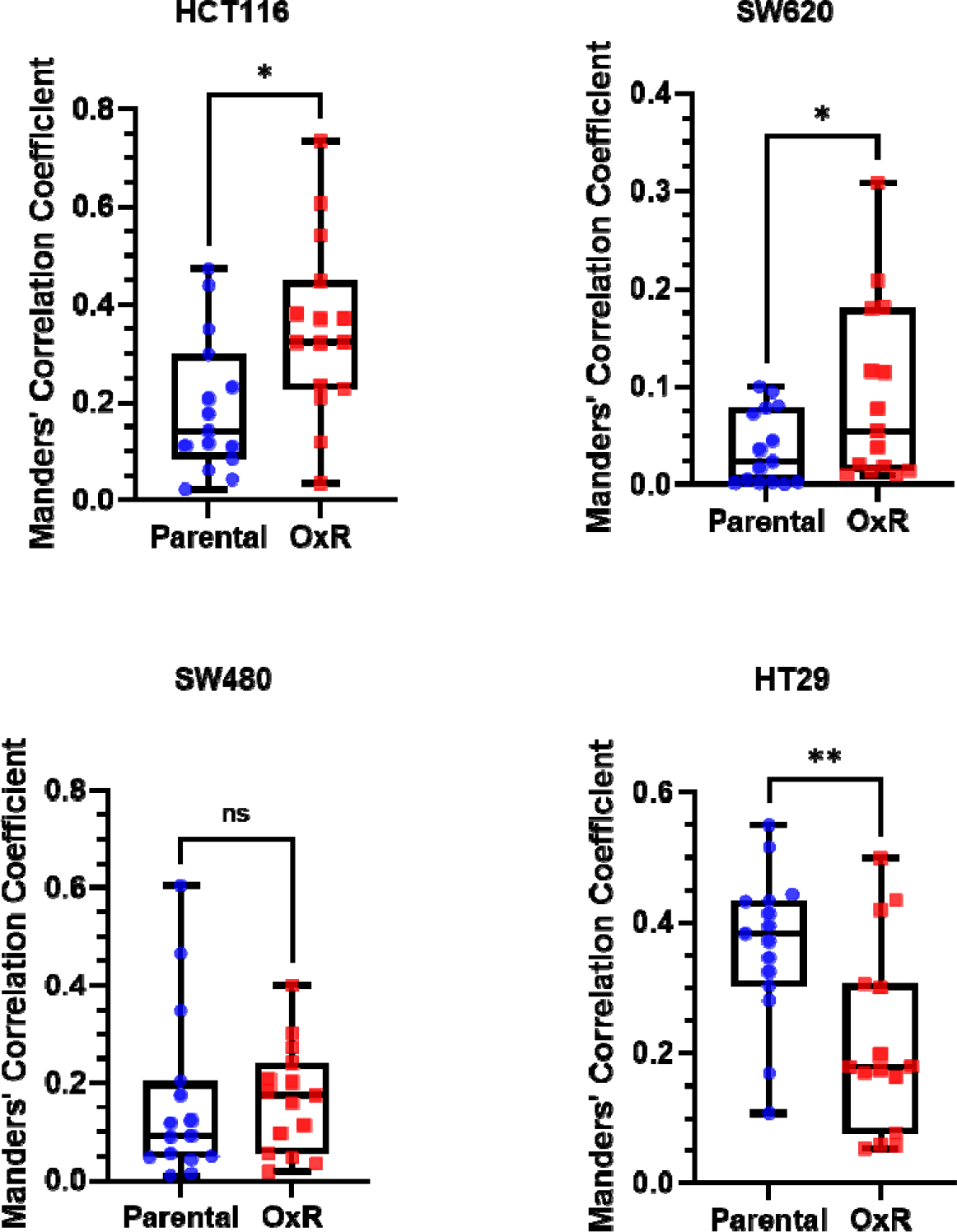
TRAIL-sensitized OxR cells have significantly increased colocalization of DR4 with lipid rafts according to Manders’ Correlation Coefficient. N=15 analyzed micrographs across three independent trials. *p<0.05 *p<0.001 (unpaired two-tailed t-test).

**Figure 4-figure supplement 2.**
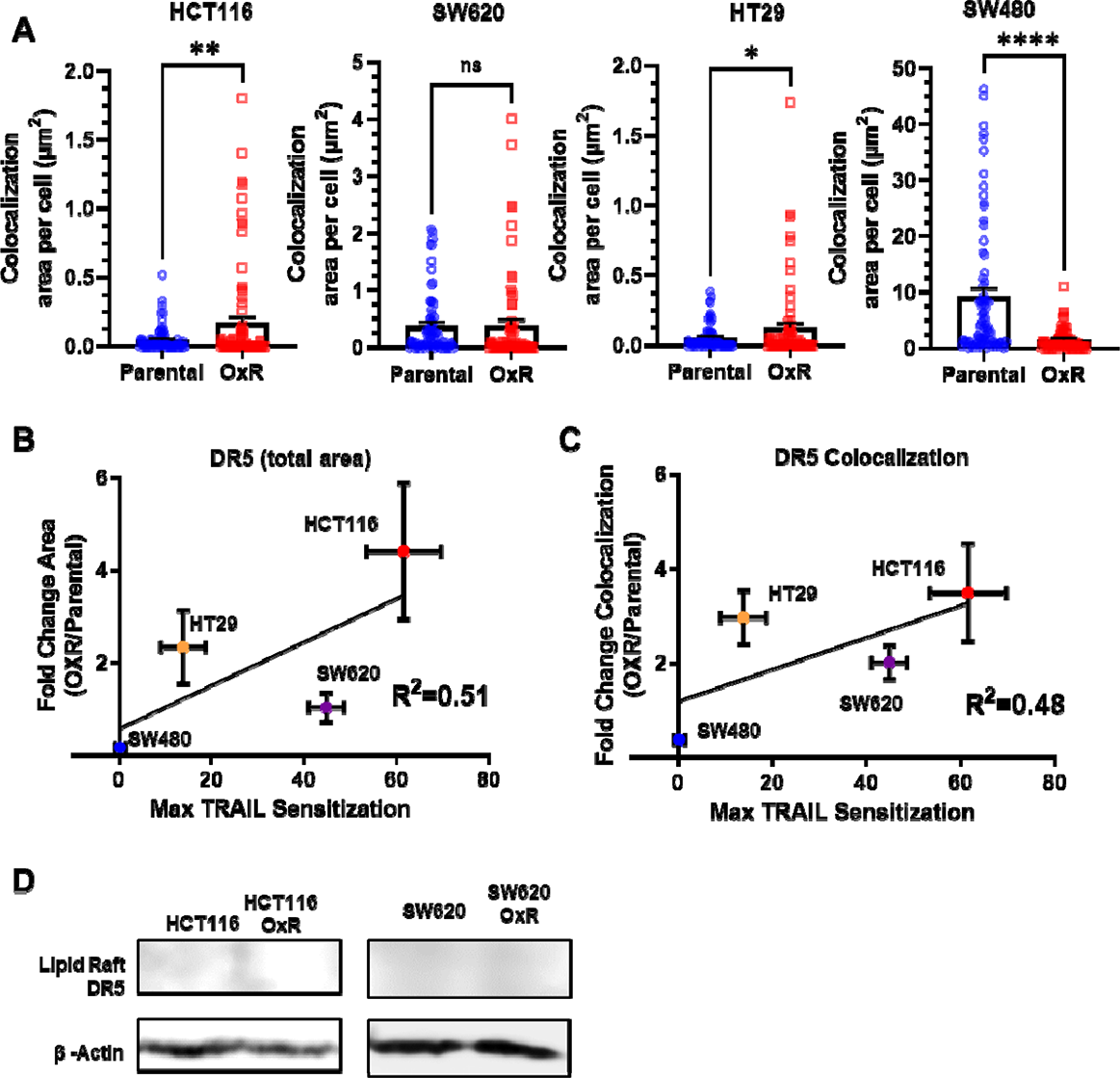
Sensitization to TRAIL in OxR cell lines poorly correlates with DR5 expression while lipid raft fractions have no detectable DR5. **(A)** Quantification of DR5/LR colocalization in HCT116, SW620, SW480 and HT29 cells. *p<0.05 **p<0.01****p<0.0001 (unpaired two-tailed t-test). For each cell line, N=75 cells were analyzed. **(B)** Correlation of total DR5 area per cell and **(C)** DR5/LR colocalization with maximum TRAIL sensitization observed in OxR cells (linear regression analysis). For all graphs, data are presented as mean+SEM. **(D)** Western blots show DR5 was undetectable in lipid raft isolated fractions.

**Figure 4-figure supplement 3.**
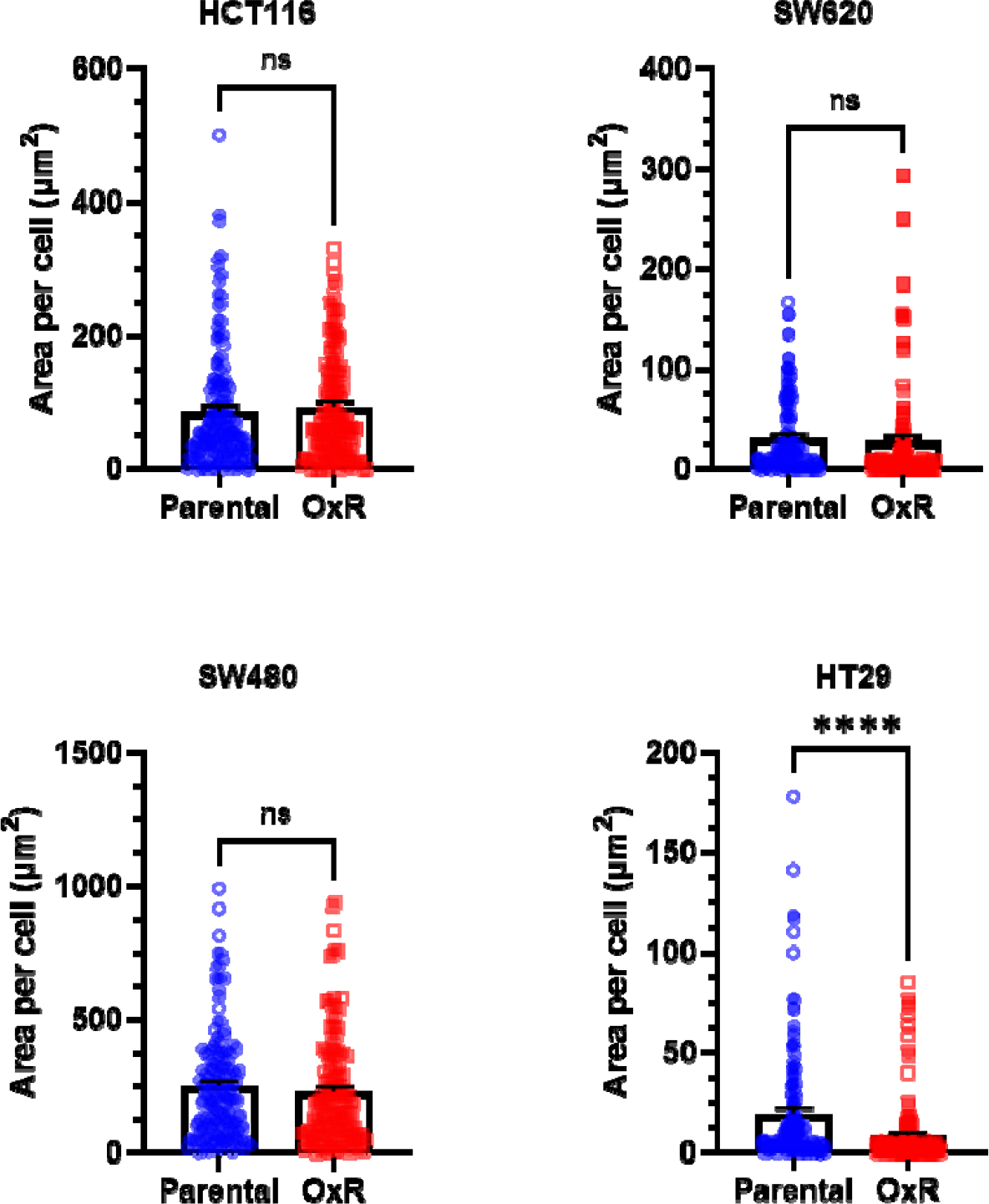
Quantification of lipid raft area per cell. For each cell line, N=150 cells were analyzed. Data are presented as mean+SEM. ****p<0.0001 (unpaired two-tailed t-test).

**Figure 5-figure supplement 1.**
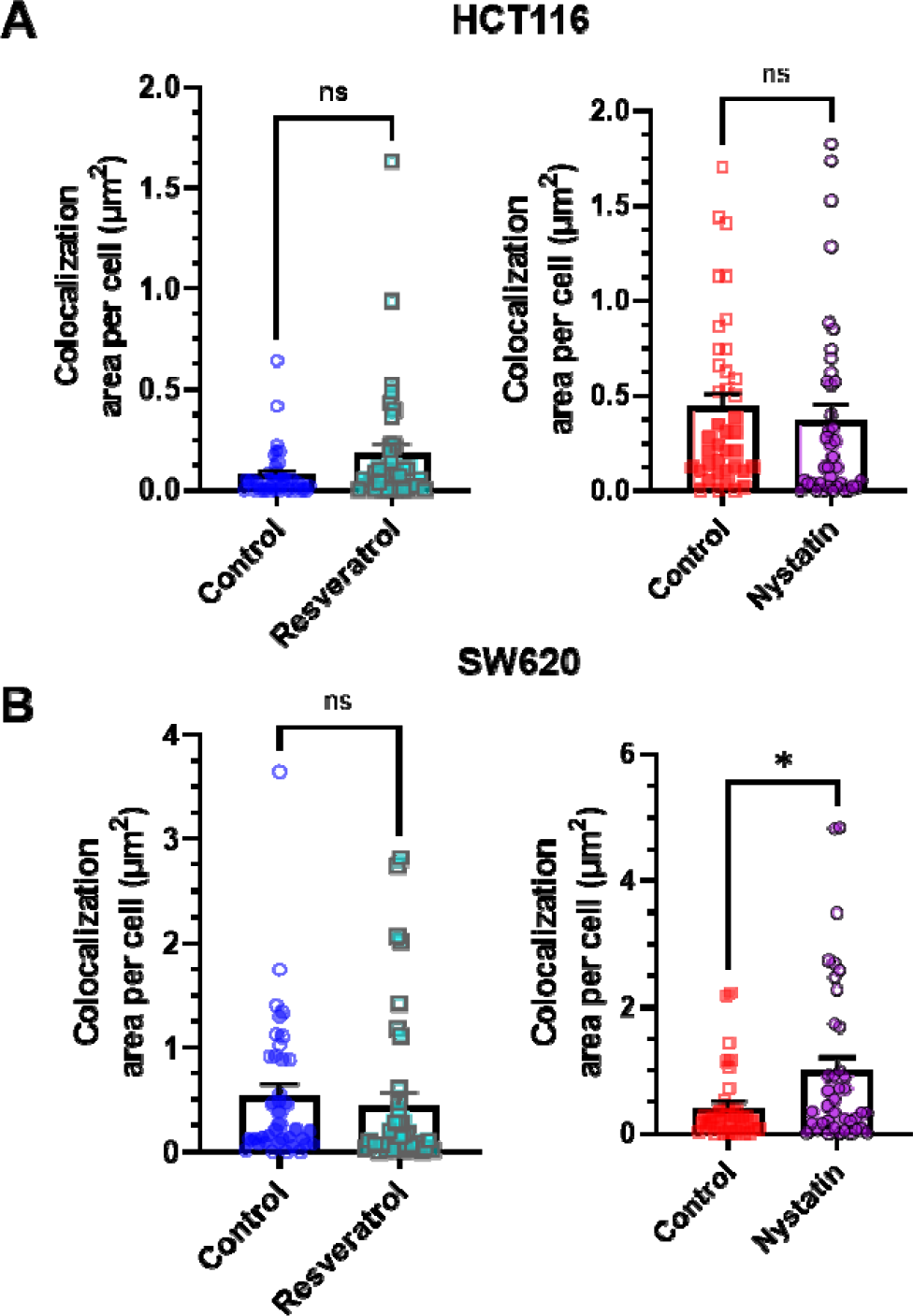
Quantification of the effects of resveratrol and nystatin on DR5 colocalization with lipid rafts in HCT116 (A) and SW620 cells (B). For each cell line, N=40 cells were analyzed. Data are presented as mean+SEM. *p<0.05 (unpaired two-tailed t-test).

**Figure 6-figure supplement 1.**
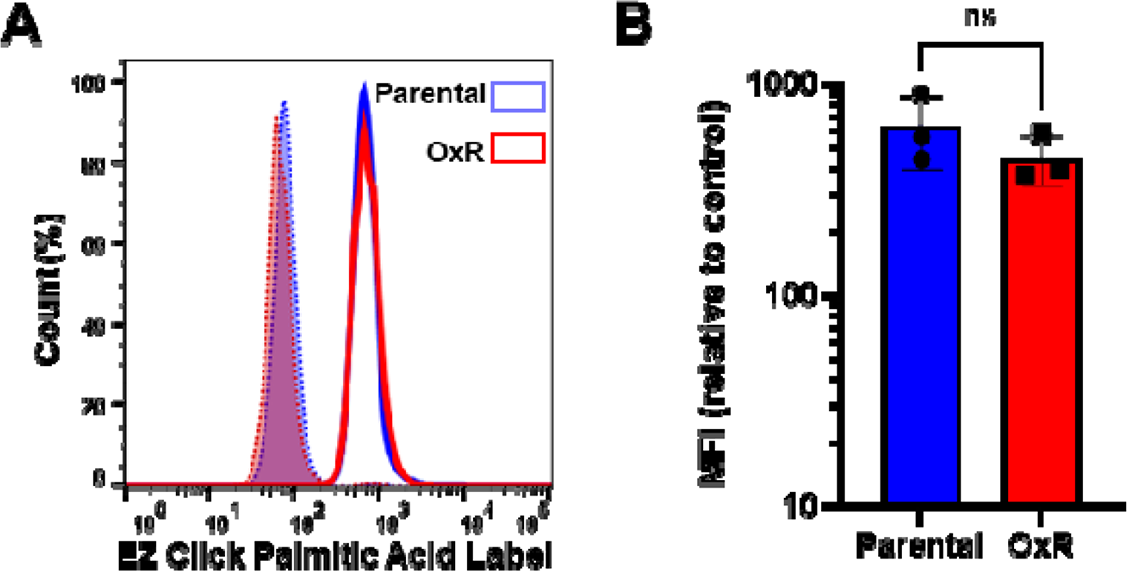
Total palmitoylation remains unchanged between SW620 parental and OxR cells. **(A)** Parental and OxR SW620 cells were labeled with EZClick^TM^ Palmitic Acid/ Fluorescent Azide staining kit and analyzed via flow cytometry to determine total protein palmitoylation between cell lines. Blue histograms represent parental SW620 cells and red histograms are SW620 OxR cells. Shaded histograms are background controls for each cell line (Palmitic Acid (**-**)/ Fluorescent Azide (**+**). **(B)** Quantification of median fluorescence intensity (MFI) shows no significant change in total palmitoylation between cellular phenotypes (unpaired two-tailed t-test). Data are presented as mean±SD. N=3. Significance was measured using an unpaired two-tailed t-test.

**Figure 7-figure supplement 1.**
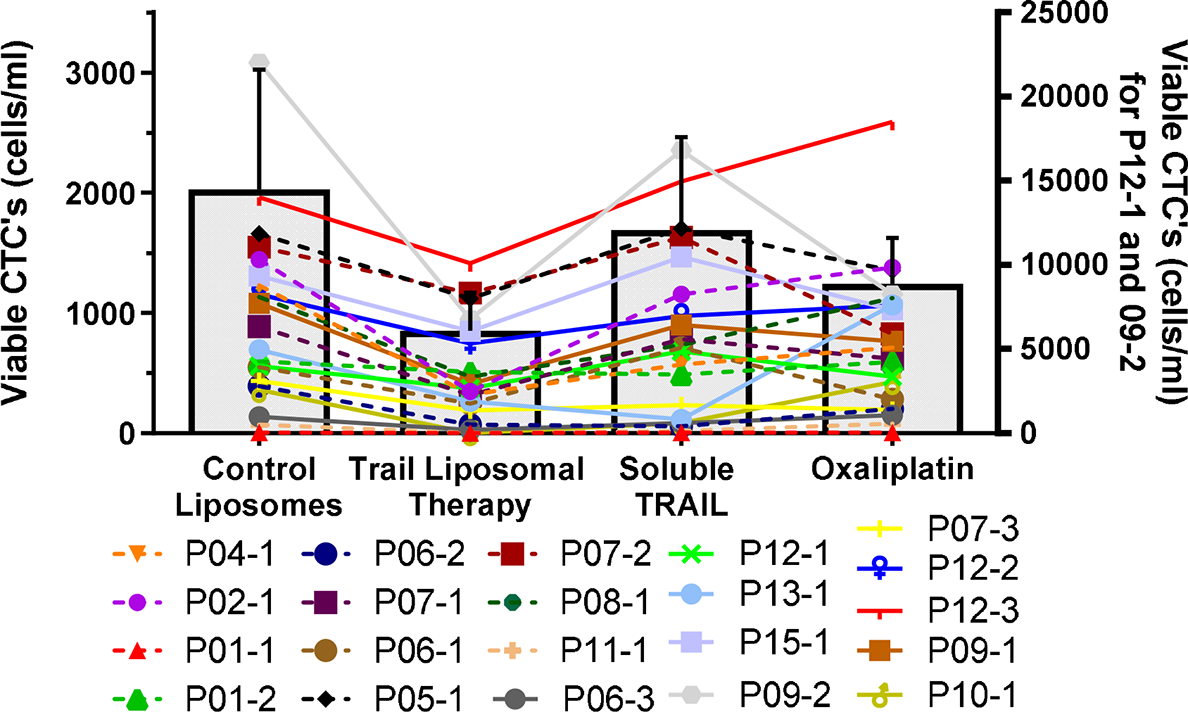
Absolute numbers of viable CTCs per ml of blood following TRAIL liposomal therapy and control treatments. Bars represent mean of all patient samples and error bars represent SEM. Samples 12-1 and 09-2 showed very large CTC concentrations and were plotted using the alternative scale shown on the right. All other samples (and average) were plotted using the scale shown on the left.

**Figure 7-figure supplement 2.**
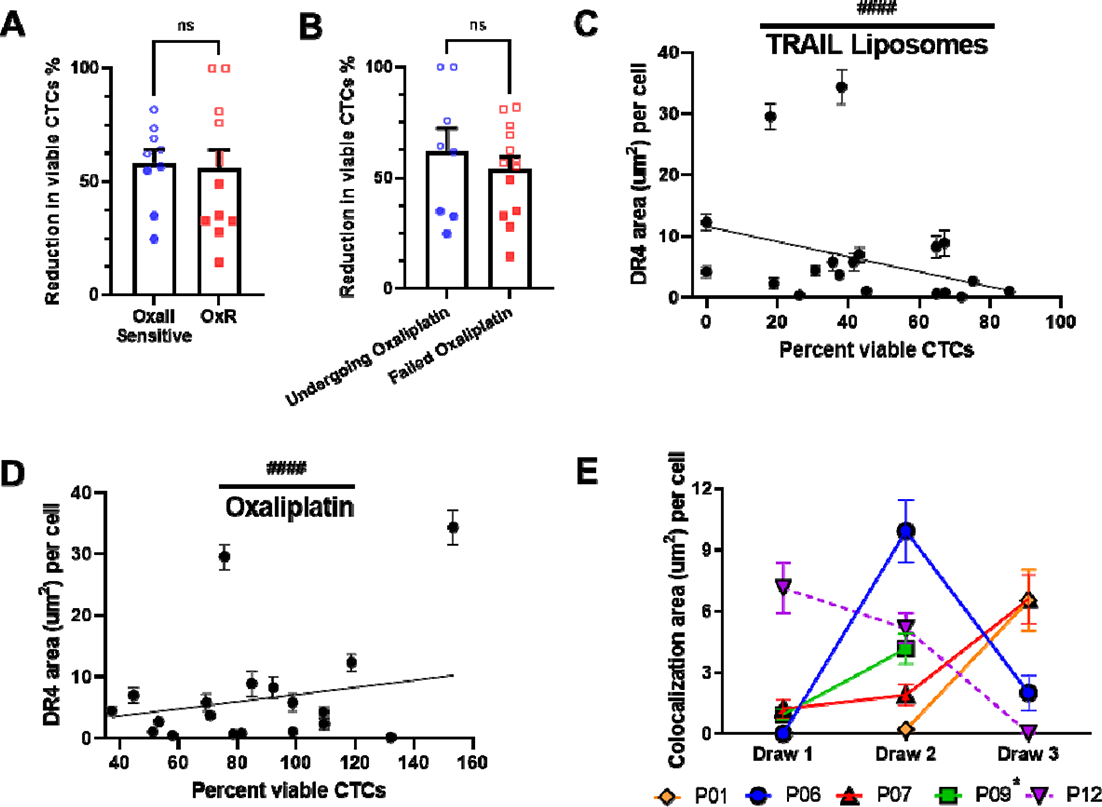
TRAIL liposomes are effective in oxaliplatin sensitive and refractory patients. **(A)** Patients were categorized as either oxaliplatin-sensitive (viability <80%, N=9) or oxaliplatin resistant (viability >80%, N=12) to compare changes in the reduction of viable CTCs (unpaired two-tailed t-test). **(B)** Patients undergoing oxaliplatin chemotherapy at the time of blood draw (N=8) showed insignificant changes in viable CTC reduction compared to patients that have previously failed oxaliplatin (N=13) (unpaired two-tailed t-test). **(C)** DR4 area of patient CTCs plotted against the percentage of viable CTCs following TRAIL liposome treatments. Each point corresponds with one patient draw. ^####^p<0.0001 (simple linear regression to confirm significant deviation from zero). **(G)** DR4 area of patient CTCs plotted against the normalized percentage of viable CTCs following oxaliplatin treatment. Each point corresponds with one patient draw. ^####^p<0.0001 (simple linear regression to confirm significant deviation from zero). **(E)** Lipid raft/DR4 analysis of repeat patients, analyzing the changes in DR4 colocalization over the course of therapy. *Patient 9 died after draw 2, precluding further blood collection. For all graphs, data are presented as mean± SEM.

**Supplementary File 1.**
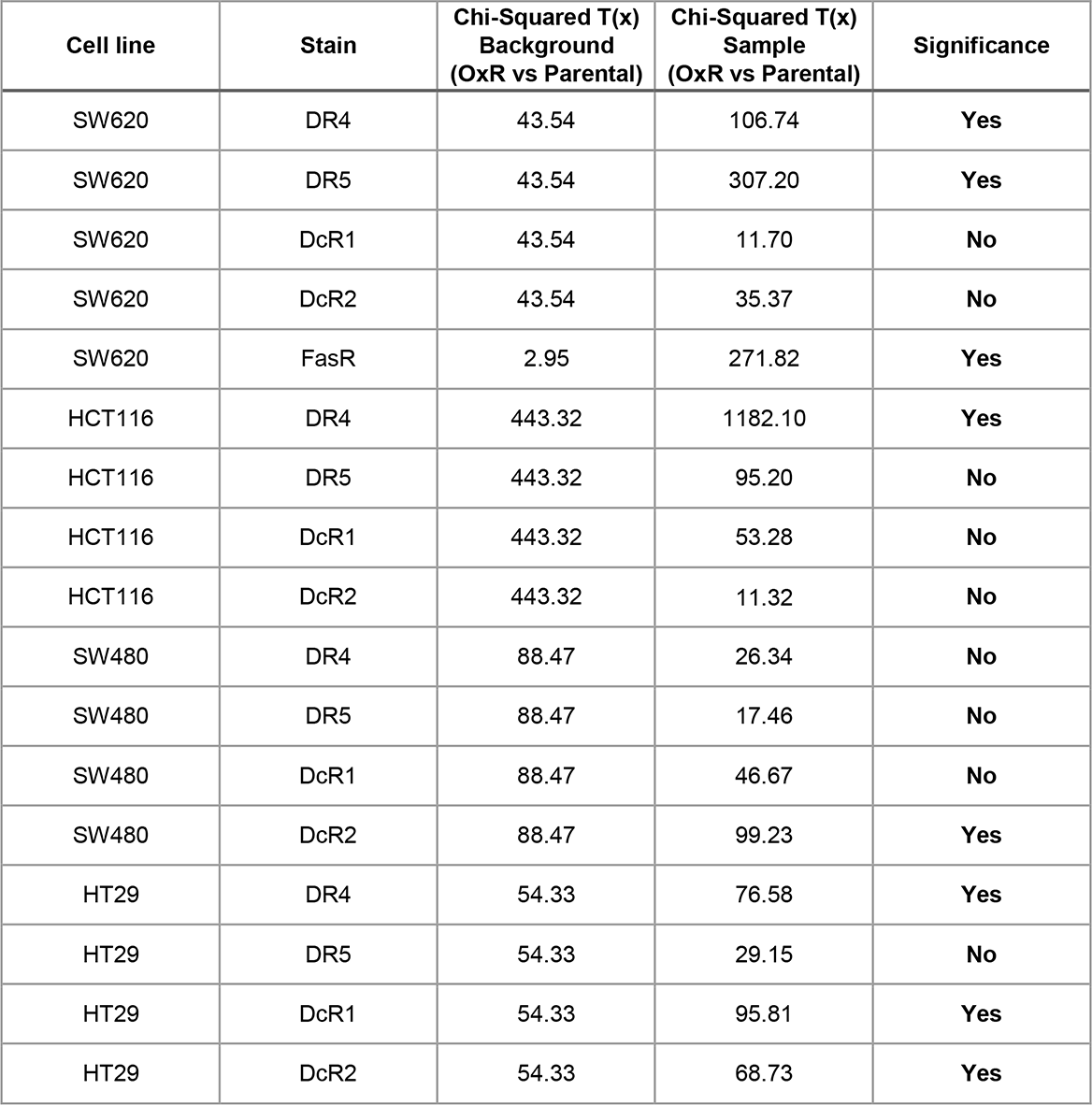
. Statistical reporting of Chi-Squared T(x) values for comparing distribution differences in flow cytometry staining. Significance was determined if Chi-Squared T(x) sample > Chi-Squared T(x) background.

## References

1. Siegel RL, Miller KD, Jemal A. Cancer statistics, 2019. CA Cancer J Clin. 2019;69:7–34.

2. Survival Rates for Colorectal Cancer [Internet]. [cited 2020 Apr 3]. Available from: https://www.cancer.org/cancer/colon-rectal-cancer/detection-diagnosis-staging/survival-rates.html

3. Mehlen P, Puisieux A. Metastasis: a question of life or death. Nat Rev Cancer. 2006;6:449–58.

4. Werner J, Heinemann V. Standards and Challenges of Care for Colorectal Cancer Today. Visc Med. 2016;32:156–7.

5. Rodriguez-Bigas MA, Lin EH, Crane CH. Stage IV Colorectal Cancer. Holl-Frei Cancer Med 6th Ed [Internet]. 2003 [cited 2019 Apr 9]; Available from: https://www.ncbi.nlm.nih.gov/books/NBK13267/

6. Briffa R, Langdon SP, Grech G, J.Harrison D. Acquired and Intrinsic Resistance to Colorectal Cancer Treatment. Colorectal Cancer - Diagn Screen Manag [Internet]. IntechOpen; 2017 [cited 2020 Apr 9]; Available from: https://www.intechopen.com/books/colorectal-cancer-diagnosis-screening-and-management/acquired-and-intrinsic-resistance-to-colorectal-cancer-treatment

7. Martinez-Balibrea E, Martínez-Cardús A, Ginés A, Porras VR de, Moutinho C, Layos L, et al. Tumor-Related Molecular Mechanisms of Oxaliplatin Resistance. Mol Cancer Ther. 2015;14:1767–76.

8. Sussman RT, Ricci MS, Hart LS, Sun SY, El-Deiry WS. Chemotherapy-resistant side-population of colon cancer cells has a higher sensitivity to TRAIL than the Non-SP, a higher expression of c-Myc and TRAIL-receptor DR4. Cancer Biol Ther. 2007;6:1486–91.

9. Van der Jeught K, Xu H-C, Li Y-J, Lu X-B, Ji G. Drug resistance and new therapies in colorectal cancer. World J Gastroenterol. 2018;24:3834–48.

10. Combès E, Andrade AF, Tosi D, Michaud H-A, Coquel F, Garambois V, et al. Inhibition of Ataxia-Telangiectasia Mutated and RAD3-Related (ATR) Overcomes Oxaliplatin Resistance and Promotes Antitumor Immunity in Colorectal Cancer. Cancer Res. 2019;79:2933–46.

11. Ruiz de Porras V, Bystrup S, Martínez-Cardús A, Pluvinet R, Sumoy L, Howells L, et al. Curcumin mediates oxaliplatin-acquired resistance reversion in colorectal cancer cell lines through modulation of CXC-Chemokine/NF-κB signalling pathway. Sci Rep [Internet]. 2016 [cited 2019 May 29];6. Available from: https://www.ncbi.nlm.nih.gov/pmc/articles/PMC4835769/

12. Cuello M, Ettenberg SA, Nau MM, Lipkowitz S. Synergistic induction of apoptosis by the combination of trail and chemotherapy in chemoresistant ovarian cancer cells. Gynecol Oncol. 2001;81:380–90.

13. von Karstedt S, Montinaro A, Walczak H. Exploring the TRAILs less travelled: TRAIL in cancer biology and therapy. Nat Rev Cancer. Nature Publishing Group; 2017;17:352–66.

14. Simons K, Toomre D. Lipid rafts and signal transduction. Nat Rev Mol Cell Biol. 2000;1:31–9.

15. Greenlee JD, Subramanian T, Liu K, King MR. Rafting Down the Metastatic Cascade: The Role of Lipid Rafts in Cancer Metastasis, Cell Death, and Clinical Outcomes. Cancer Res. American Association for Cancer Research; 2021;81:5–17.

16. de Laurentiis A, Donovan L, Arcaro A. Lipid Rafts and Caveolae in Signaling by Growth Factor Receptors. Open Biochem J. 2007;1:12–32.

17. Marconi M, Ascione B, Ciarlo L, Vona R, Garofalo T, Sorice M, et al. Constitutive localization of DR4 in lipid rafts is mandatory for TRAIL-induced apoptosis in B-cell hematologic malignancies. Cell Death Dis. Nature Publishing Group; 2013;4:e863–e863.

18. Mollinedo F, Gajate C. Lipid rafts as signaling hubs in cancer cell survival/death and invasion: implications in tumor progression and therapy. J Lipid Res. American Society for Biochemistry and Molecular Biology; 2020;jlr.TR119000439.

19. George KS, Wu S. Lipid Raft: A Floating Island Of Death or Survival. Toxicol Appl Pharmacol. 2012;259:311–9.

20. Naval J, de Miguel D, Gallego-Lleyda A, Anel A, Martinez-Lostao L. Importance of TRAIL Molecular Anatomy in Receptor Oligomerization and Signaling. Implications for Cancer Therapy. Cancers [Internet]. 2019 [cited 2020 Feb 27];11. Available from: https://www.ncbi.nlm.nih.gov/pmc/articles/PMC6521207/

21. Nagane M, Pan G, Weddle JJ, Dixit VM, Cavenee WK, Huang HJ. Increased death receptor 5 expression by chemotherapeutic agents in human gliomas causes synergistic cytotoxicity with tumor necrosis factor-related apoptosis-inducing ligand in vitro and in vivo. Cancer Res. 2000;60:847–53.

22. Gibson SB, Oyer R, Spalding AC, Anderson SM, Johnson GL. Increased Expression of Death Receptors 4 and 5 Synergizes the Apoptosis Response to Combined Treatment with Etoposide and TRAIL. Mol Cell Biol. 2000;20:205–12.

23. Baritaki S, Huerta-Yepez S, Sakai T, Spandidos DA, Bonavida B. Chemotherapeutic drugs sensitize cancer cells to TRAIL-mediated apoptosis: up-regulation of DR5 and inhibition of Yin Yang 1. Mol Cancer Ther. American Association for Cancer Research; 2007;6:1387– 99.

24. El Fajoui Z, Toscano F, Jacquemin G, Abello J, Scoazec J-Y, Micheau O, et al. Oxaliplatin sensitizes human colon cancer cells to TRAIL through JNK-dependent phosphorylation of Bcl-xL. Gastroenterology. 2011;141:663–73.

25. Xu L, Qu X, Zhang Y, Hu X, Yang X, Hou K, et al. Oxaliplatin enhances TRAIL-induced apoptosis in gastric cancer cells by CBL-regulated death receptor redistribution in lipid rafts. FEBS Lett. 2009;583:943–8.

26. Dallas NA, Xia L, Fan F, Gray MJ, Gaur P, van Buren G, et al. Chemoresistant Colorectal Cancer Cells, the Cancer Stem Cell Phenotype and Increased Sensitivity to Insulin-like Growth Factor Receptor-1 Inhibition. Cancer Res. 2009;69:1951–7.

27. Yang AD, Fan F, Camp ER, Buren G van, Liu W, Somcio R, et al. Chronic Oxaliplatin Resistance Induces Epithelial-to-Mesenchymal Transition in Colorectal Cancer Cell Lines. Clin Cancer Res. 2006;12:4147–53.

28. Tanaka S, Hosokawa M, Yonezawa T, Hayashi W, Ueda K, Iwakawa S. Induction of epithelial-mesenchymal transition and down-regulation of miR-200c and miR-141 in oxaliplatin-resistant colorectal cancer cells. Biol Pharm Bull. 2015;38:435–40.

29. Garrido C, Galluzzi L, Brunet M, Puig PE, Didelot C, Kroemer G. Mechanisms of cytochrome c release from mitochondria. Cell Death Differ. Nature Publishing Group; 2006;13:1423–33.

30. Hitomi J, Katayama T, Eguchi Y, Kudo T, Taniguchi M, Koyama Y, et al. Involvement of caspase-4 in endoplasmic reticulum stress-induced apoptosis and Abeta-induced cell death. J Cell Biol. 2004;165:347–56.

31. Ozören N, El-Deiry WS. Cell surface Death Receptor signaling in normal and cancer cells. Semin Cancer Biol. 2003;13:135–47.

32. Sandra F, Hendarmin L, Nakamura S. Osteoprotegerin (OPG) binds with tumor necrosis factor-related apoptosis-inducing ligand (TRAIL): suppression of TRAIL-induced apoptosis in ameloblastomas. Oral Oncol. 2006;42:415–20.

33. Si W, Shen J, Zheng H, Fan W. The role and mechanisms of action of microRNAs in cancer drug resistance. Clin Epigenetics. 2019;11:25.

34. Just C, Knief J, Lazar-Karsten P, Petrova E, Hummel R, Röcken C, et al. MicroRNAs as Potential Biomarkers for Chemoresistance in Adenocarcinomas of the Esophagogastric Junction [Internet]. J. Oncol. Hindawi; 2019 [cited 2020 Apr 17]. page e4903152. Available from: https://www.hindawi.com/journals/jo/2019/4903152/

35. Toscano F, Fajoui ZE, Gay F, Lalaoui N, Parmentier B, Chayvialle J-A, et al. p53-Mediated upregulation of DcR1 impairs oxaliplatin/TRAIL-induced synergistic anti-tumour potential in colon cancer cells. Oncogene. Nature Publishing Group; 2008;27:4161–71.

36. Suprynowicz FA, Disbrow GL, Krawczyk E, Simic V, Lantzky K, Schlegel R. HPV-16 E5 oncoprotein upregulates lipid raft components caveolin-1 and ganglioside GM1 at the plasma membrane of cervical cells. Oncogene. Nature Publishing Group; 2008;27:1071–8.

37. Xiao W, Ishdorj G, Sun J, Johnston JB, Gibson SB. Death receptor 4 is preferentially recruited to lipid rafts in chronic lymphocytic leukemia cells contributing to tumor necrosis related apoptosis inducing ligand-induced synergistic apoptotic responses. Leuk Lymphoma. Taylor & Francis; 2011;52:1290–301.

38. Ujlaky-Nagy L, Nagy P, Szöllő i J, Vereb G. Flow Cytometric FRET Analysis of Protein Interactions. Methods Mol Biol Clifton NJ. 2018;1678:393–419.

39. Neves AR, Nunes C, Reis S. Resveratrol induces ordered domains formation in biomembranes: Implication for its pleiotropic action. Biochim Biophys Acta BBA - Biomembr. 2016;1858:12–8.

40. Rossin A, Derouet M, Abdel-Sater F, Hueber A-O. Palmitoylation of the TRAIL receptor DR4 confers an efficient TRAIL-induced cell death signalling. Biochem J. 2009;419:185– 92, 2 p following 192.

41. Draper JM, Smith CD. Palmitoyl acyltransferase assays and inhibitors (Review). Mol Membr Biol. 2009;26:5–13.

42. Stuckey DW, Shah K. TRAIL on Trial: Preclinical advances for cancer therapy. Trends Mol Med [Internet]. 2013 [cited 2017 Dec 7];19. Available from: https://www.ncbi.nlm.nih.gov/pmc/articles/PMC3880796/

43. Ortiz-Otero N, Marshall JR, Lash B, King MR. Chemotherapy-induced release of circulating-tumor cells into the bloodstream in collective migration units with cancer-associated fibroblasts in metastatic cancer patients. BMC Cancer. 2020;20:873.

44. Mitchell MJ, Wayne E, Rana K, Schaffer CB, King MR. TRAIL-coated leukocytes that kill cancer cells in the circulation. Proc Natl Acad Sci. 2014;111:930–5.

45. Jyotsana N, Zhang Z, Himmel LE, Yu F, King MR. Minimal dosing of leukocyte targeting TRAIL decreases triple-negative breast cancer metastasis following tumor resection. Sci Adv. American Association for the Advancement of Science; 2019;5:eaaw4197.

46. Mitchell MJ, King MR. Fluid shear stress sensitizes cancer cells to receptor-mediated apoptosis via trimeric death receptors. New J Phys. 2013;15:015008.

47. Mühlethaler-Mottet A, Flahaut M, Bourloud KB, Nardou K, Coulon A, Liberman J, et al. Individual caspase-10 isoforms play distinct and opposing roles in the initiation of death receptor-mediated tumour cell apoptosis. Cell Death Dis. Nature Publishing Group; 2011;2:e125–e125.

48. Sprick MR, Rieser E, Stahl H, Grosse-Wilde A, Weigand MA, Walczak H. Caspase-10 is recruited to and activated at the native TRAIL and CD95 death-inducing signalling complexes in a FADD-dependent manner but can not functionally substitute caspase-8. EMBO J. 2002;21:4520–30.

49. Horn S, Hughes MA, Schilling R, Sticht C, Tenev T, Ploesser M, et al. Caspase-10 Negatively Regulates Caspase-8-Mediated Cell Death, Switching the Response to CD95L in Favor of NF-κB Activation and Cell Survival. Cell Rep. 2017;19:785–97.

50. . Gasiulė S, Dreize N, Kaupinis A, Ražanskas R, Čiupas L, Stankevičius V, et al. Molecular Insights into miRNA-Driven Resistance to 5-Fluorouracil and Oxaliplatin Chemotherapy: miR-23b Modulates the Epithelial–Mesenchymal Transition of Colorectal Cancer Cells. J Clin Med [Internet]. 2019 [cited 2020 Apr 17];8. Available from: https://www.ncbi.nlm.nih.gov/pmc/articles/PMC6947029

51. Evert J, Pathak S, Sun X-F, Zhang H. A Study on Effect of Oxaliplatin in MicroRNA Expression in Human Colon Cancer. J Cancer. 2018;9:2046–53.

52. Delmas D, Rébé C, Micheau O, Athias A, Gambert P, Grazide S, et al. Redistribution of CD95, DR4 and DR5 in rafts accounts for the synergistic toxicity of resveratrol and death receptor ligands in colon carcinoma cells. Oncogene. 2004;23:8979–86.

53. Ouyang W, Yang C, Zhang S, Liu Y, Yang B, Zhang J, et al. Absence of death receptor translocation into lipid rafts in acquired TRAIL-resistant NSCLC cells. Int J Oncol. 2013;42:699–711.

54. Song JH, Tse MCL, Bellail A, Phuphanich S, Khuri F, Kneteman NM, et al. Lipid Rafts and Nonrafts Mediate Tumor Necrosis Factor–Related Apoptosis-Inducing Ligand–Induced Apoptotic and Nonapoptotic Signals in Non–Small Cell Lung Carcinoma Cells. Cancer Res. American Association for Cancer Research; 2007;67:6946–55.

55. Chakrabandhu K, Hérincs Z, Huault S, Dost B, Peng L, Conchonaud F, et al. Palmitoylation is required for efficient Fas cell death signaling. EMBO J. 2007;26:209–20.

56. . Zhang X-L, Ding H-H, Xu T, Liu M, Ma C, Wu S-L, et al. Palmitoylation of δ-catenin promotes kinesin-mediated membrane trafficking of Nav1.6 in sensory neurons to promote neuropathic pain. Sci Signal [Internet]. American Association for the Advancement of Science; 2018 [cited 2020 Apr 17];11. Available from: https://stke.sciencemag.org/content/11/523/eaar4394

57. . Tseng W, Leong X, Engleman E. Orthotopic Mouse Model of Colorectal Cancer. J Vis Exp JoVE [Internet]. 2007 [cited 2018 Aug 20]; Available from: https://www.ncbi.nlm.nih.gov/pmc/articles/PMC2557075/

58. Bolte S, Cordelières FP. A guided tour into subcellular colocalization analysis in light microscopy. J Microsc. 2006;224:213–32.

