## Supplementary figures and images for "Oxaliplatin Resistance in Colorectal Cancer Enhances TRAIL Sensitivity Via Death Receptor 4 Upregulation and Lipid Raft Localization"

### Figure 1-figure supplement 1

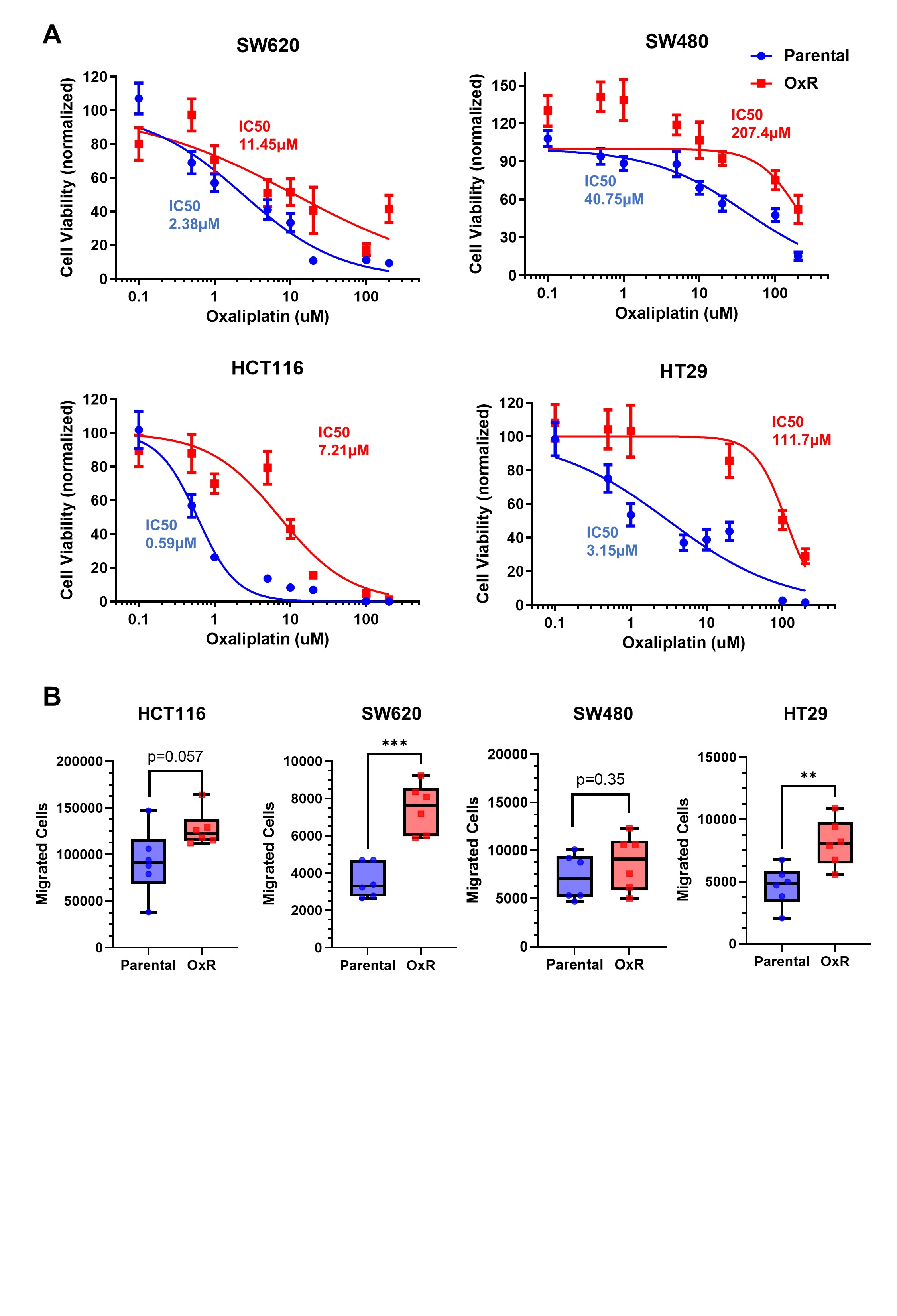

### Figure 1-figure supplement 2

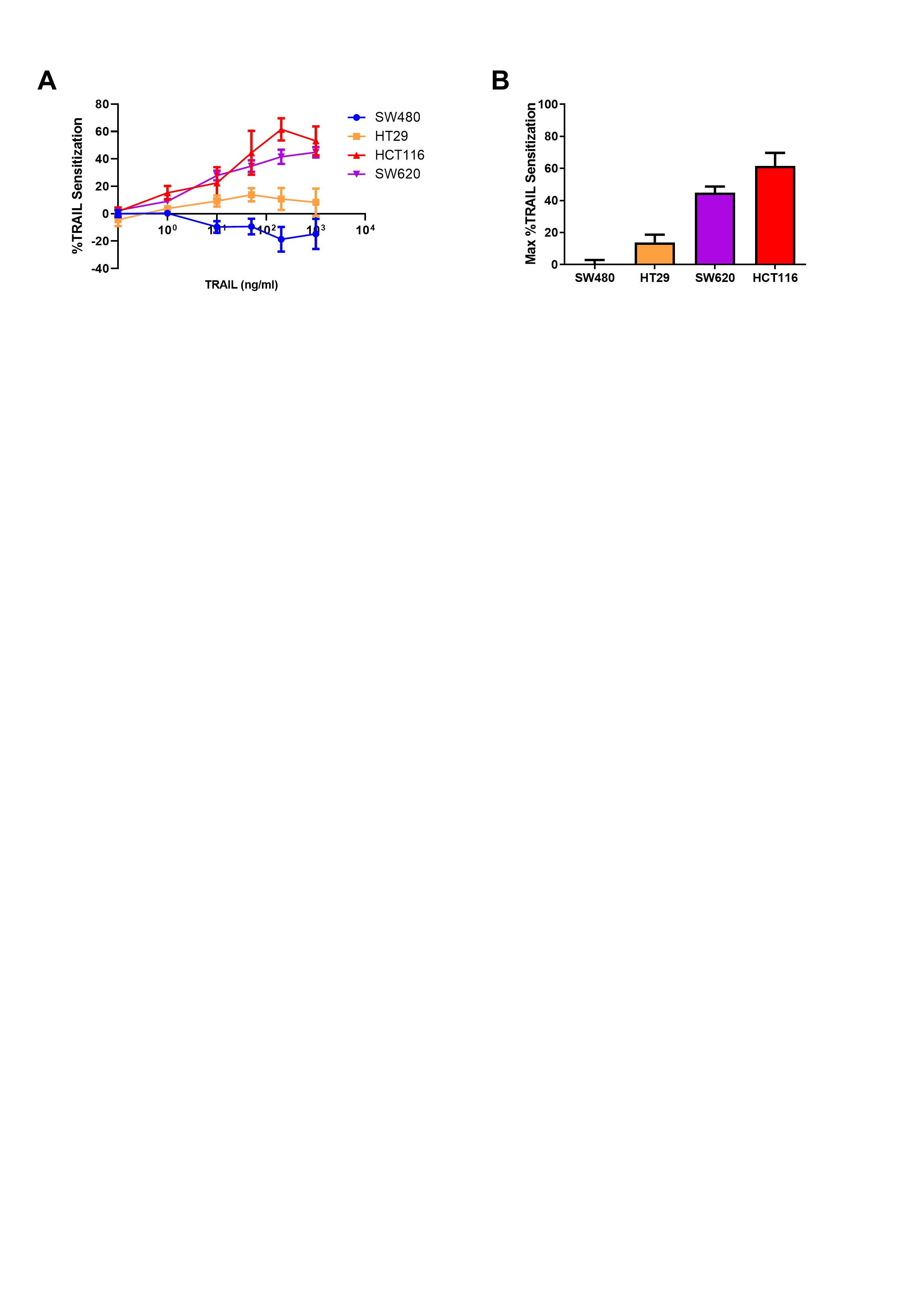

### Figure 1-figure supplement 3

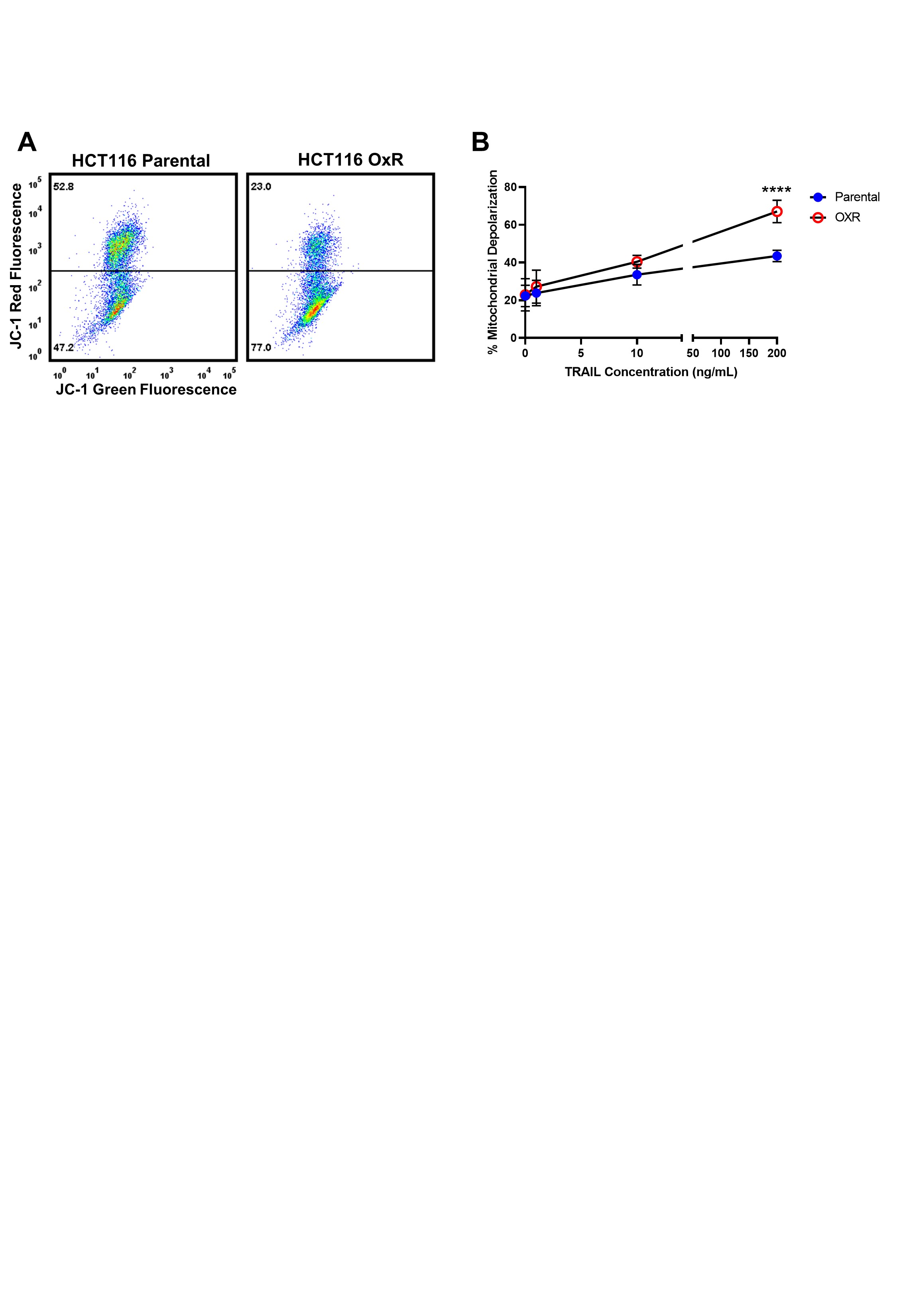

### Figure 2-figure supplement 1

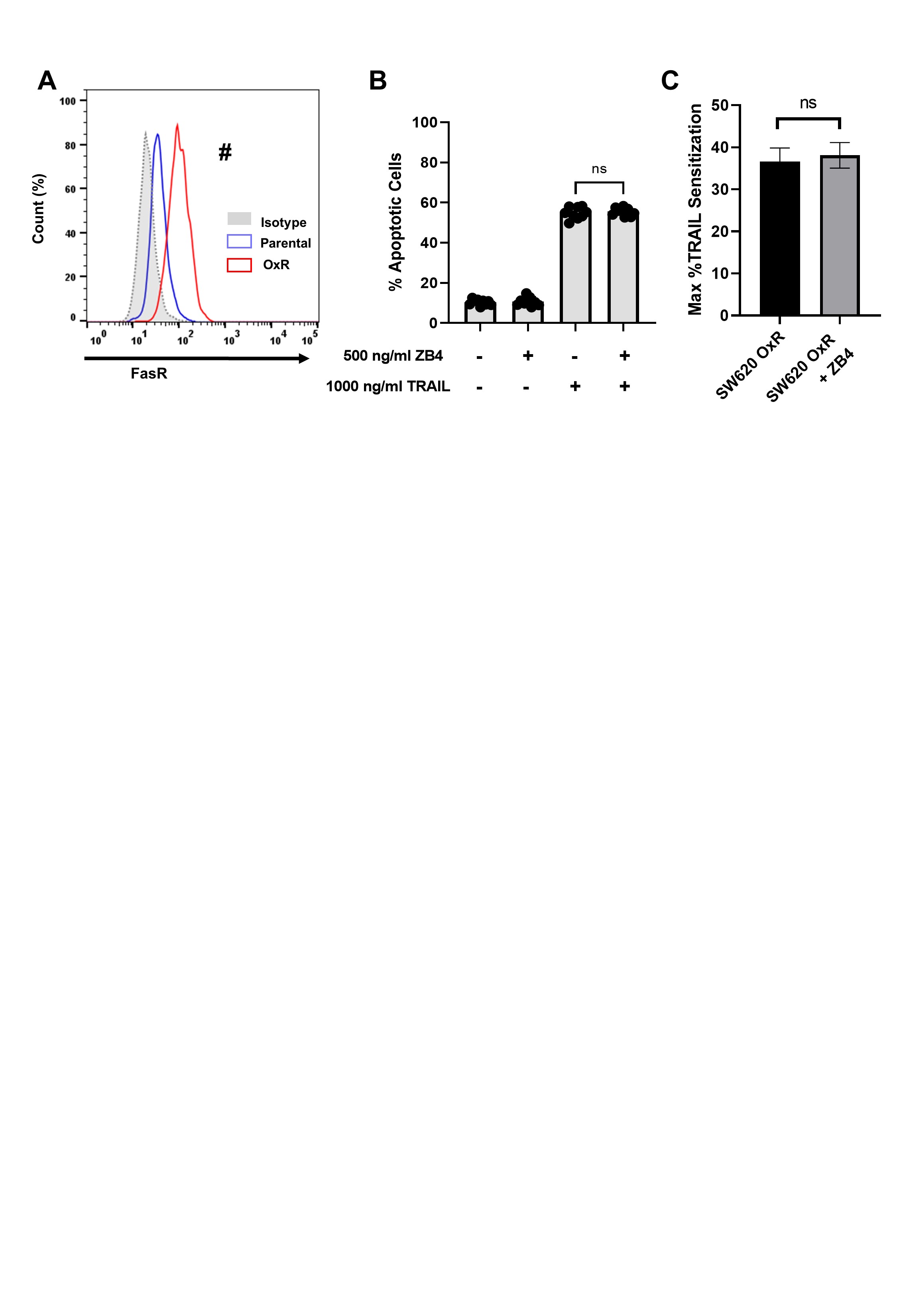

### Figure 3-figure supplement 1

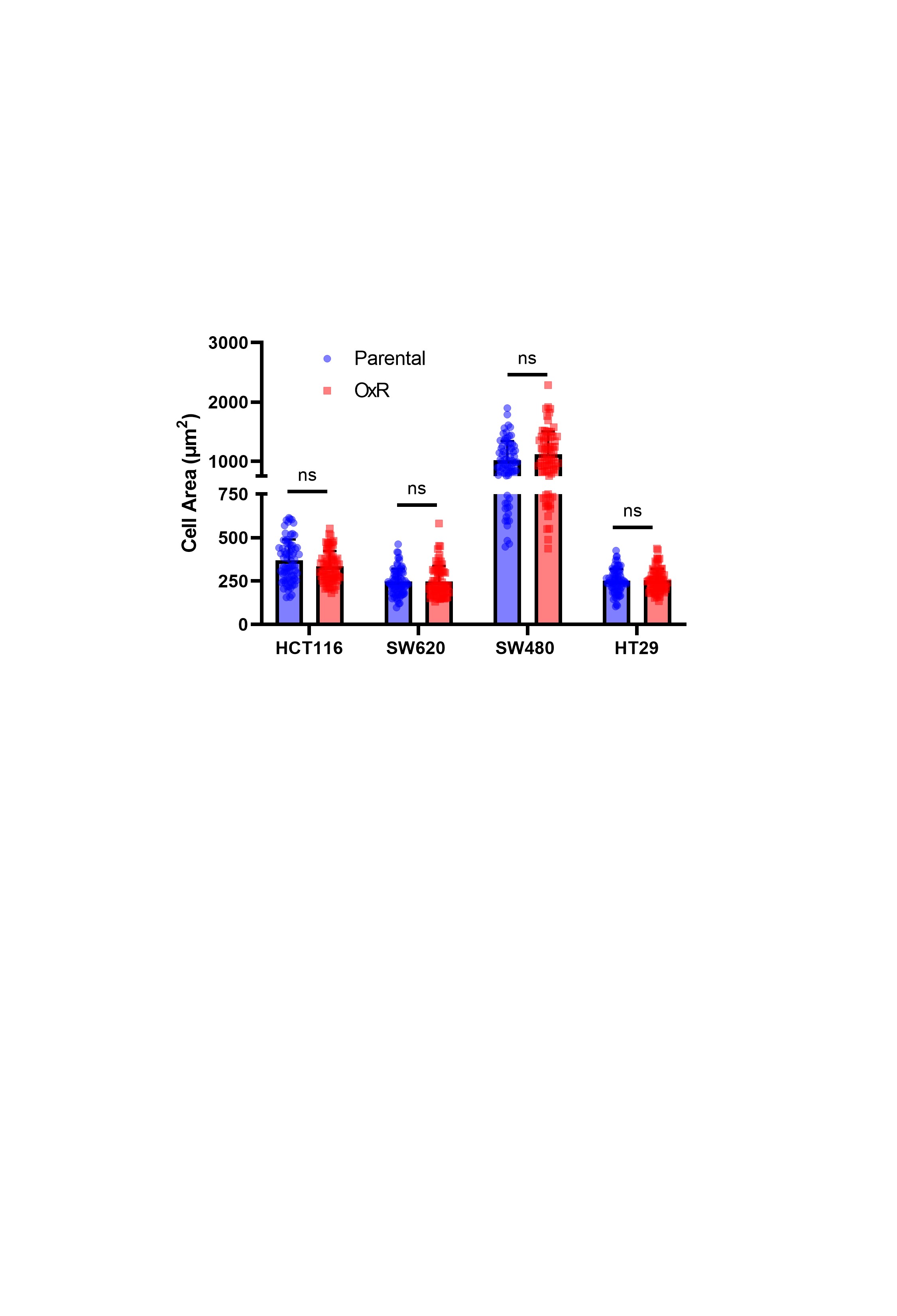

### Figure 3-figure supplement 2

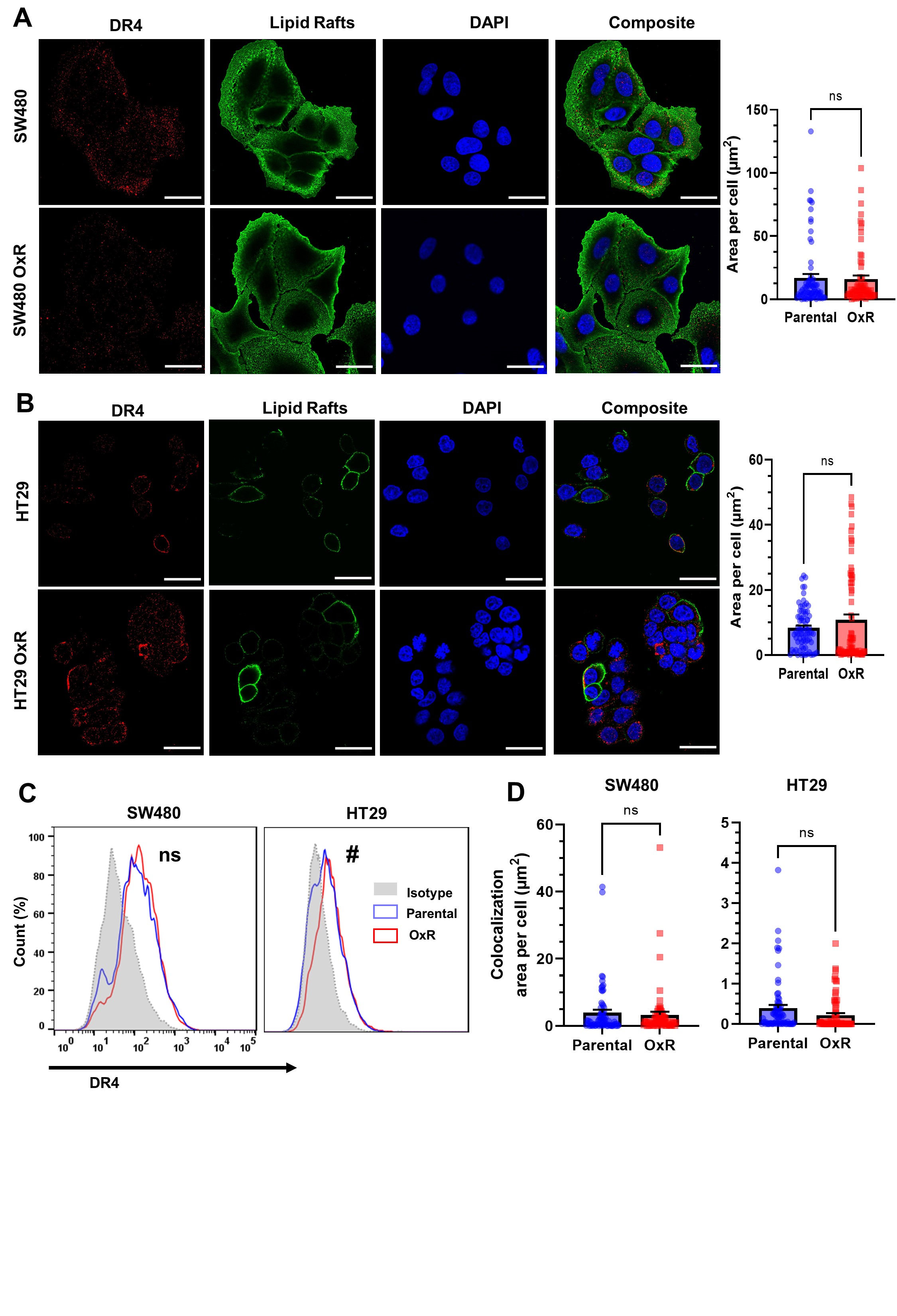

### Figure 3-figure supplement 3

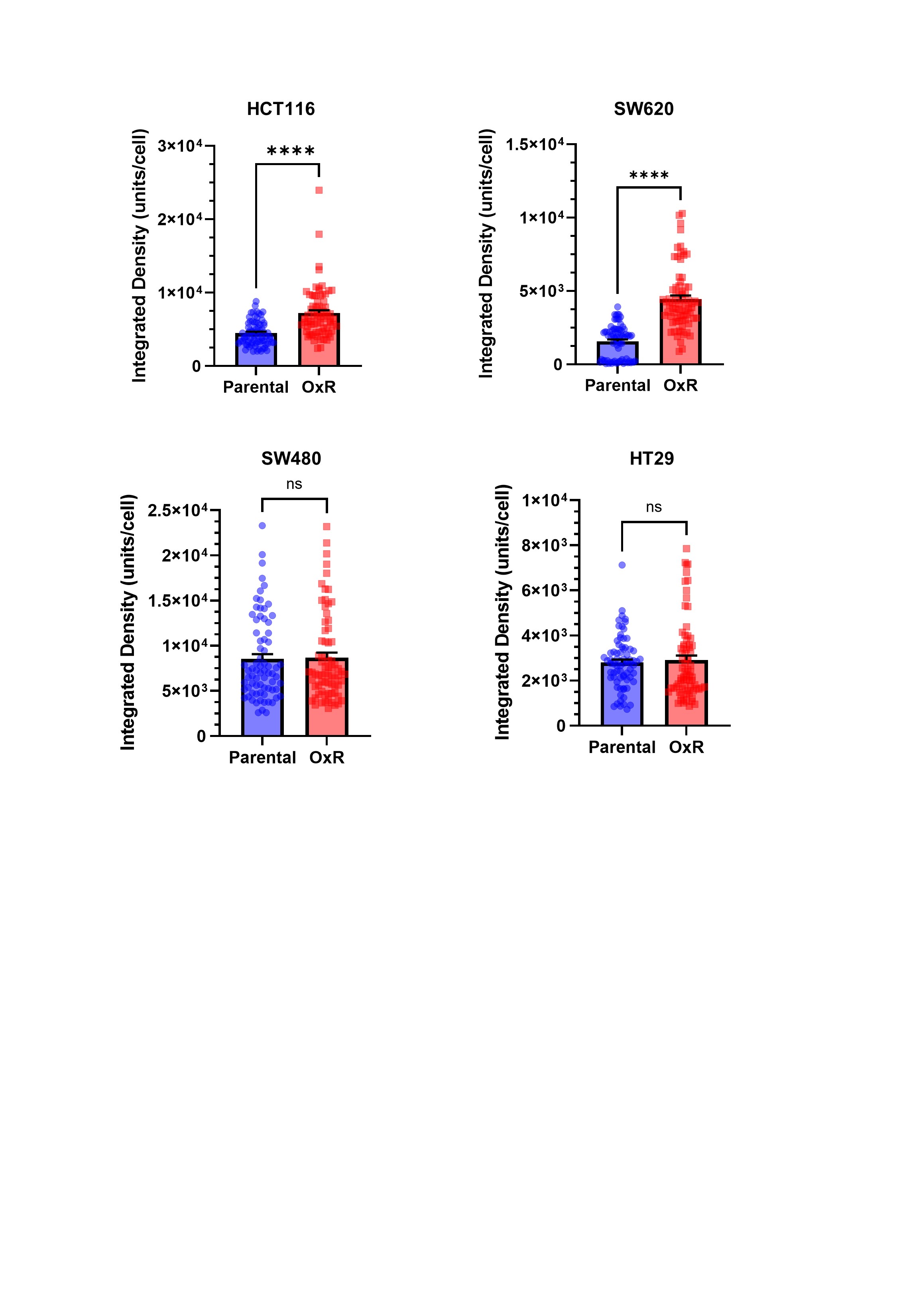

### Figure 3-figure supplement 4

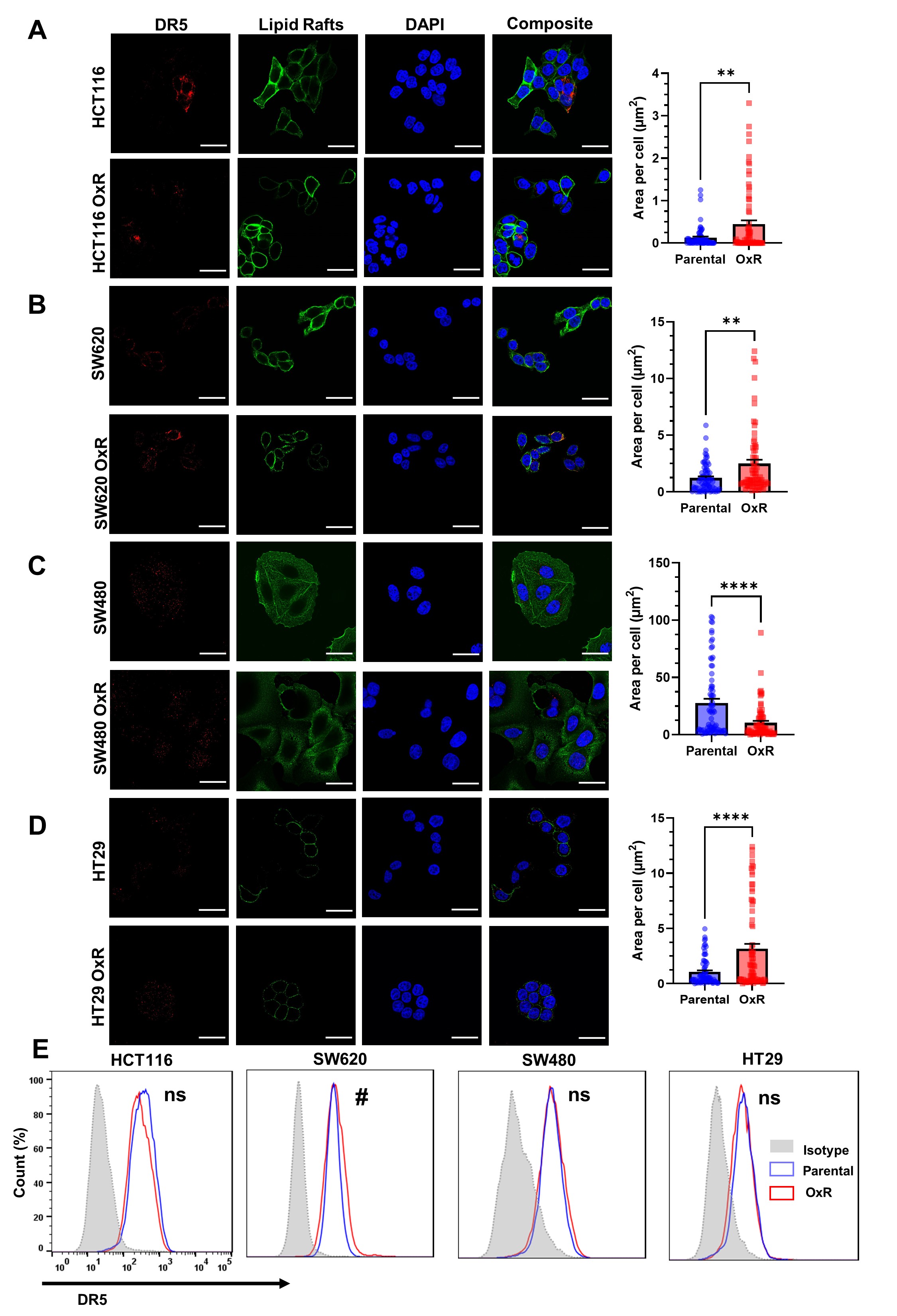

### Figure 3-figure supplement 5

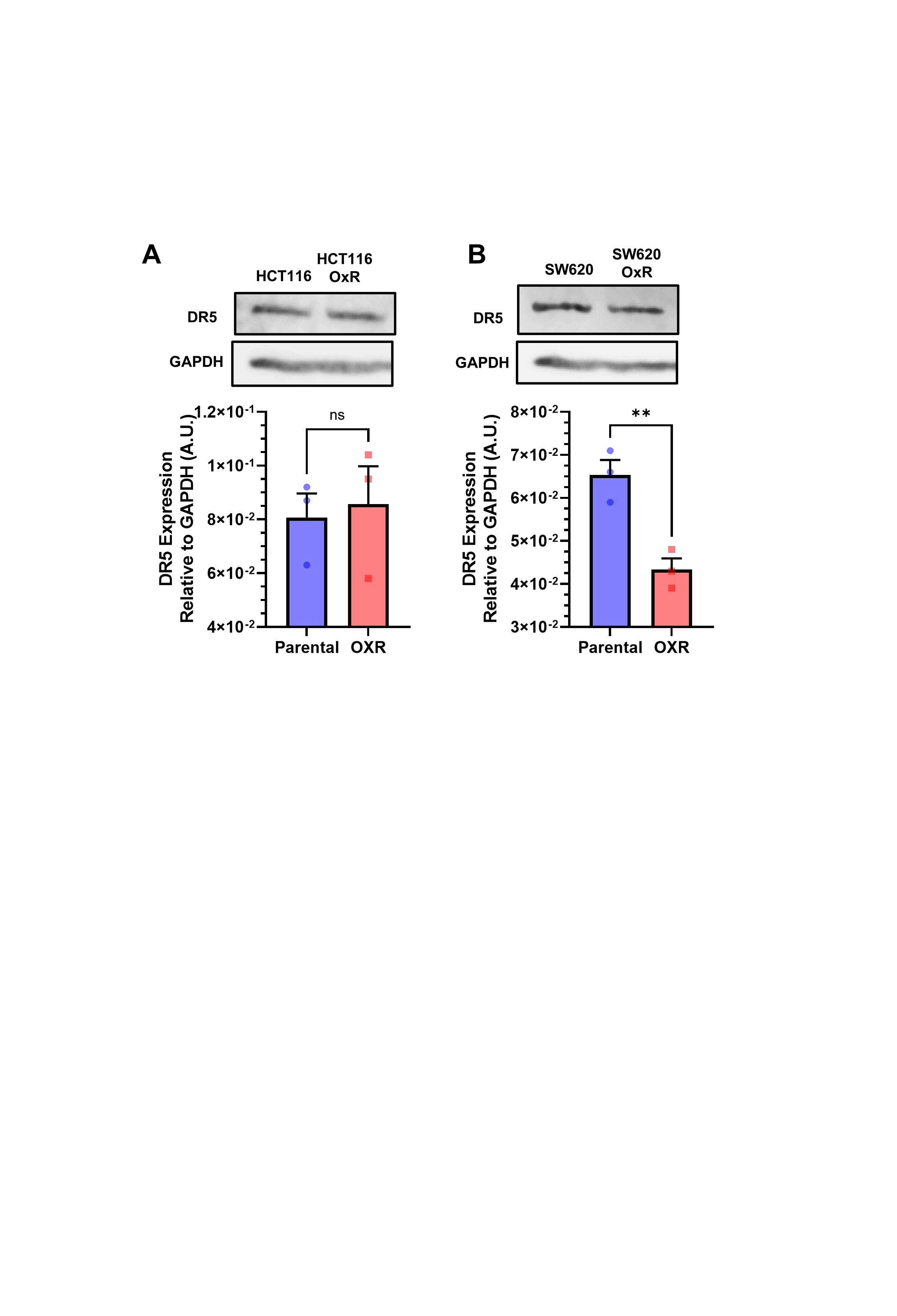

### Figure 3-figure supplement 6

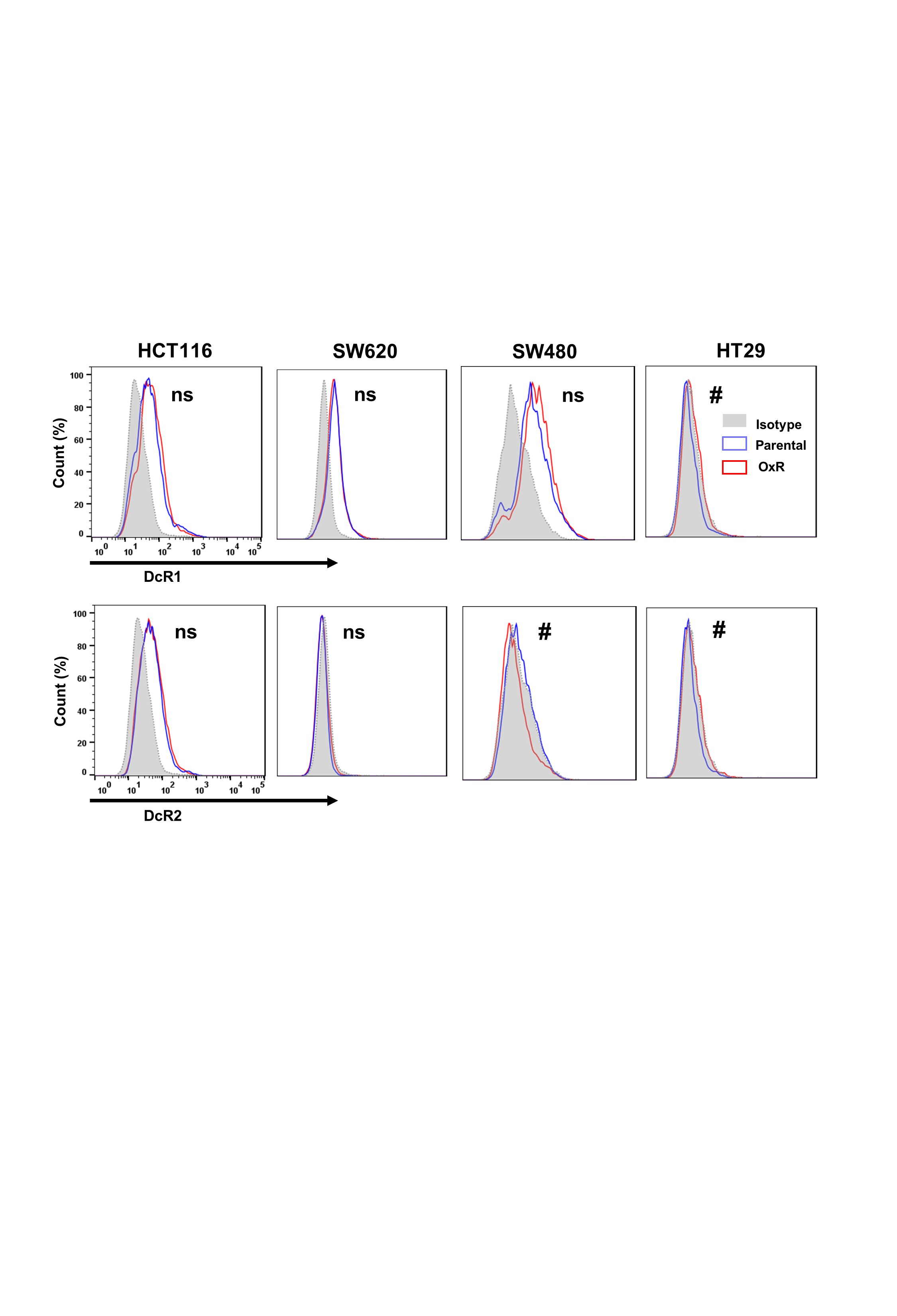

### Figure 3-figure supplement 7

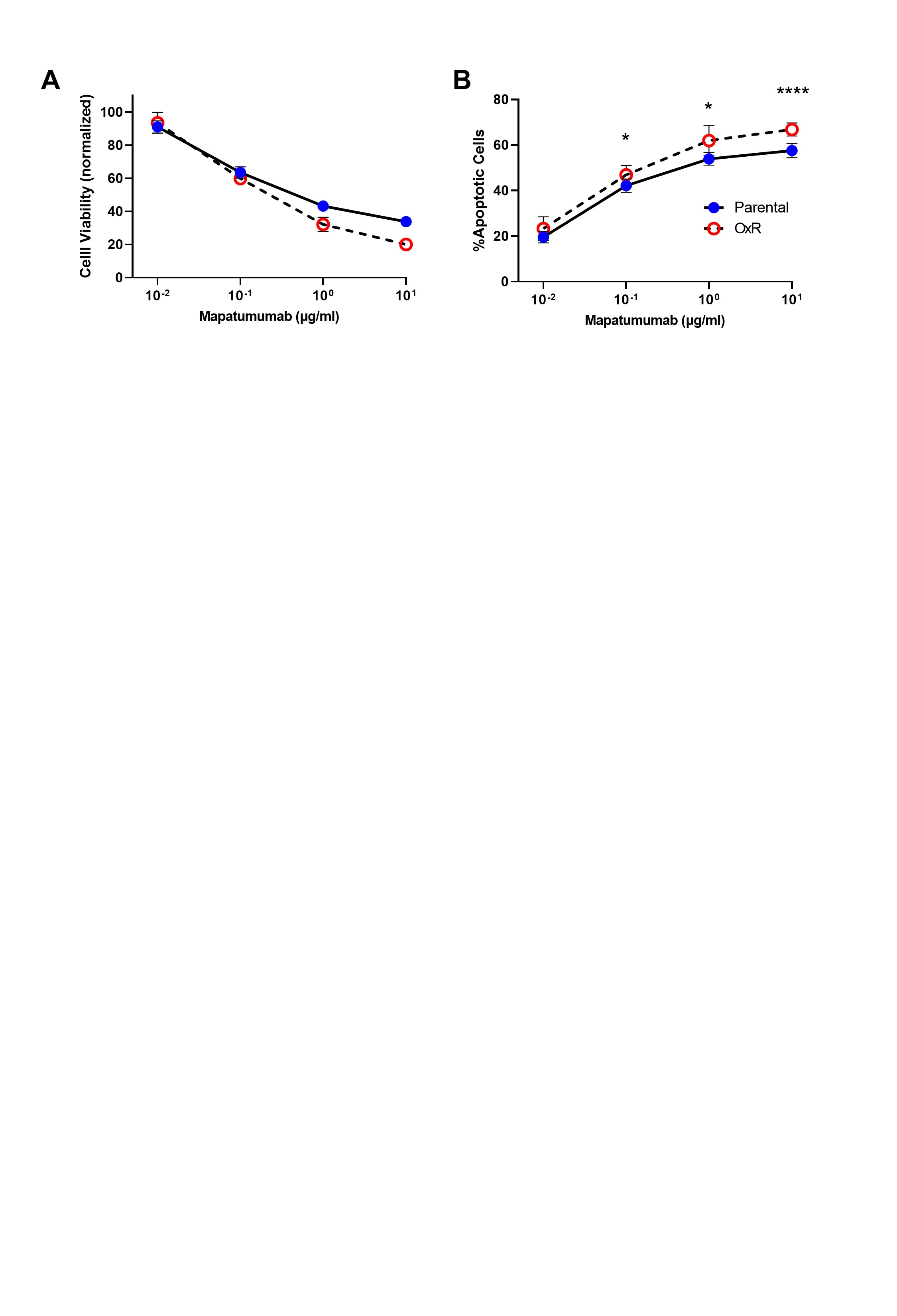

### Figure 4-figure supplement 1

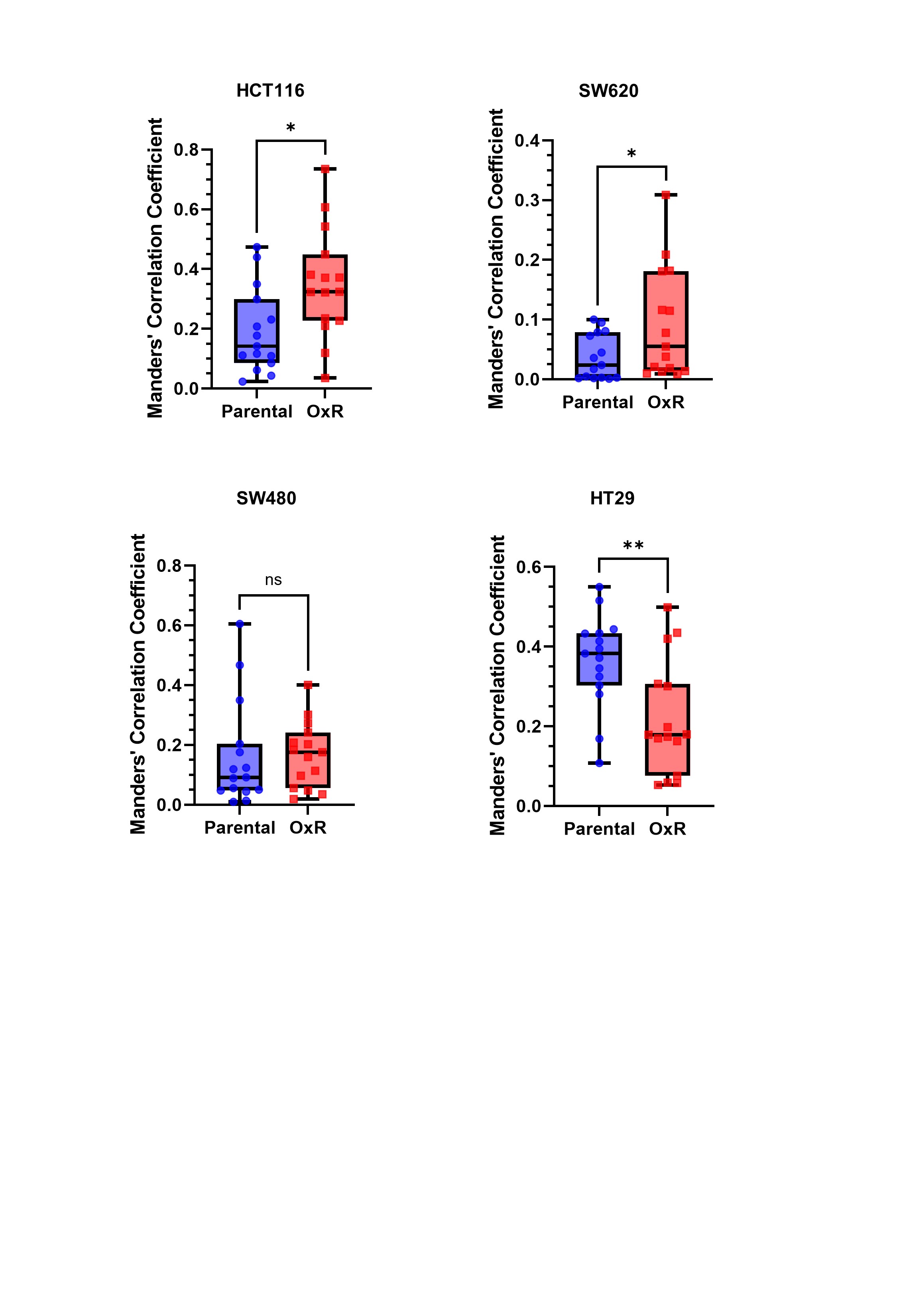

### Figure 4-figure supplement 2

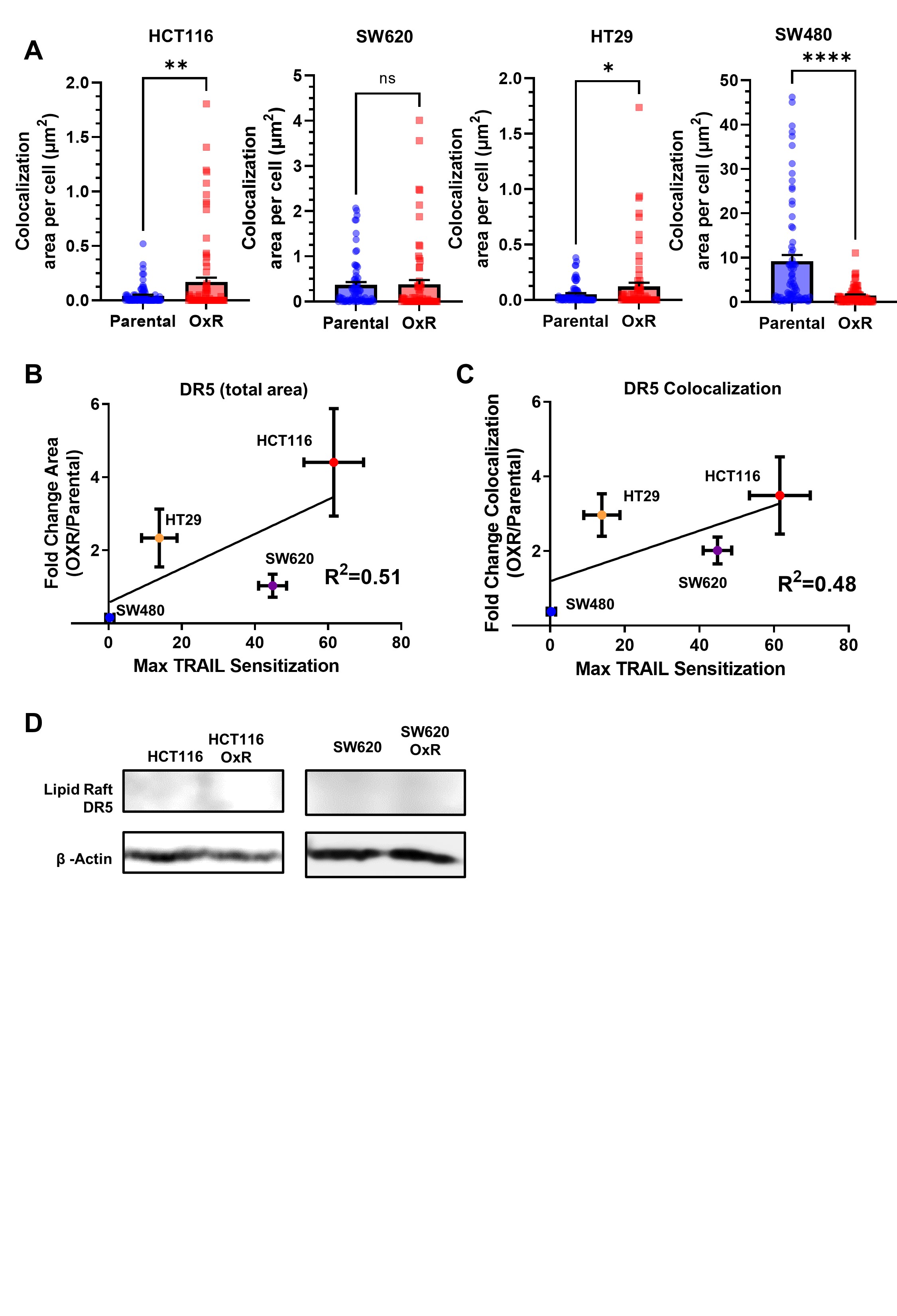

### Figure 4-figure supplement 3

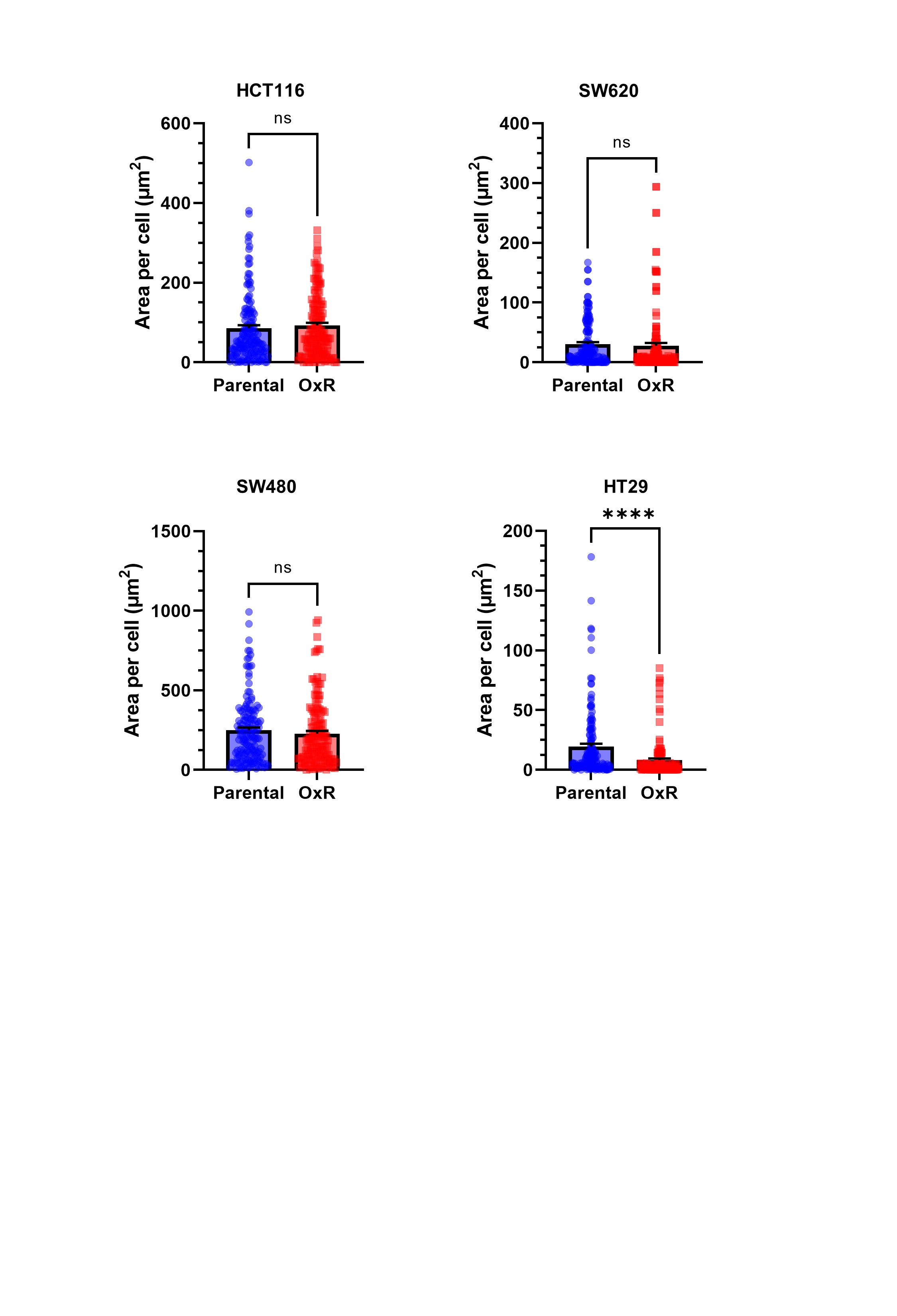

### Figure 5-figure supplement 1

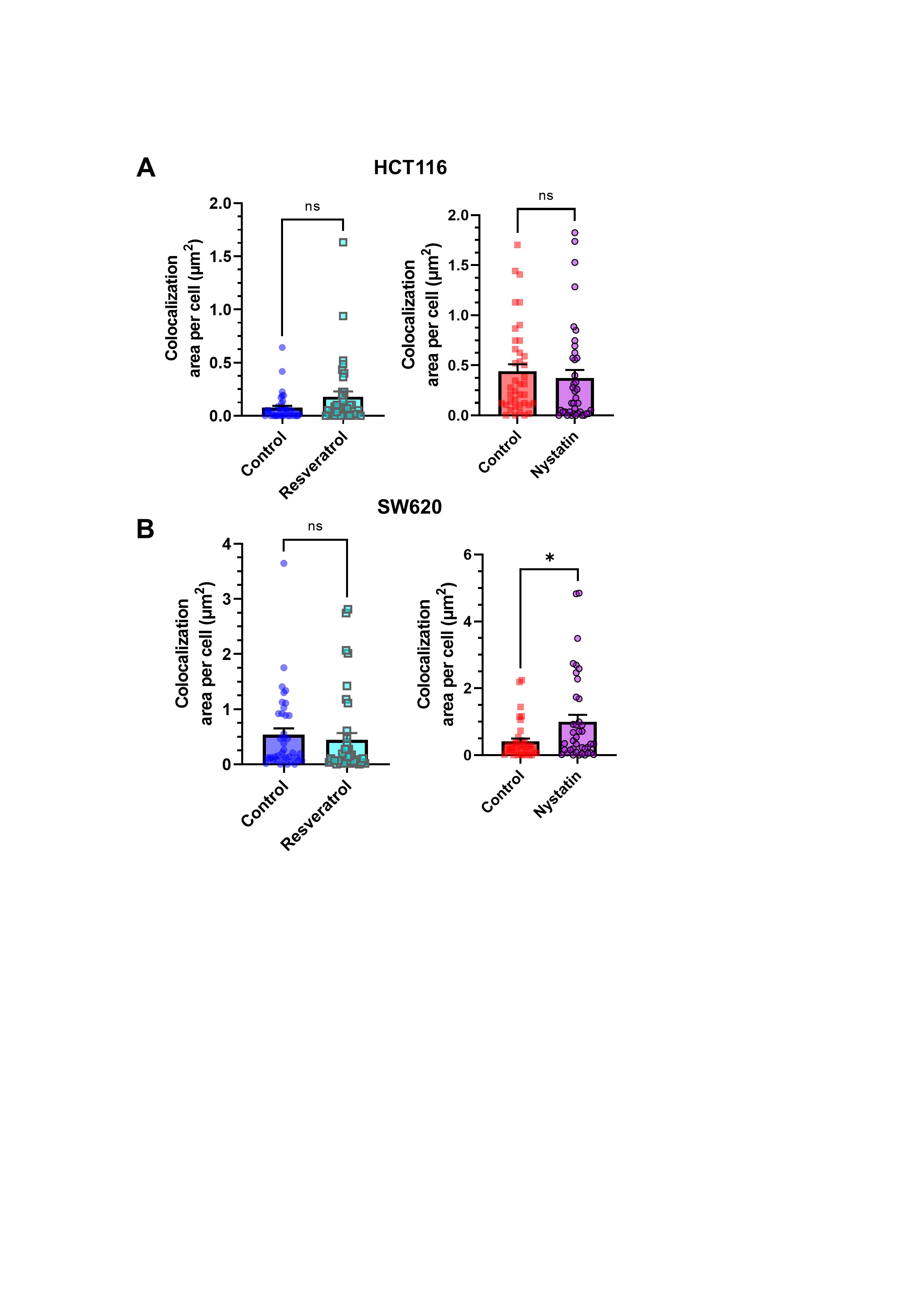

### Figure 6-figure supplement 1

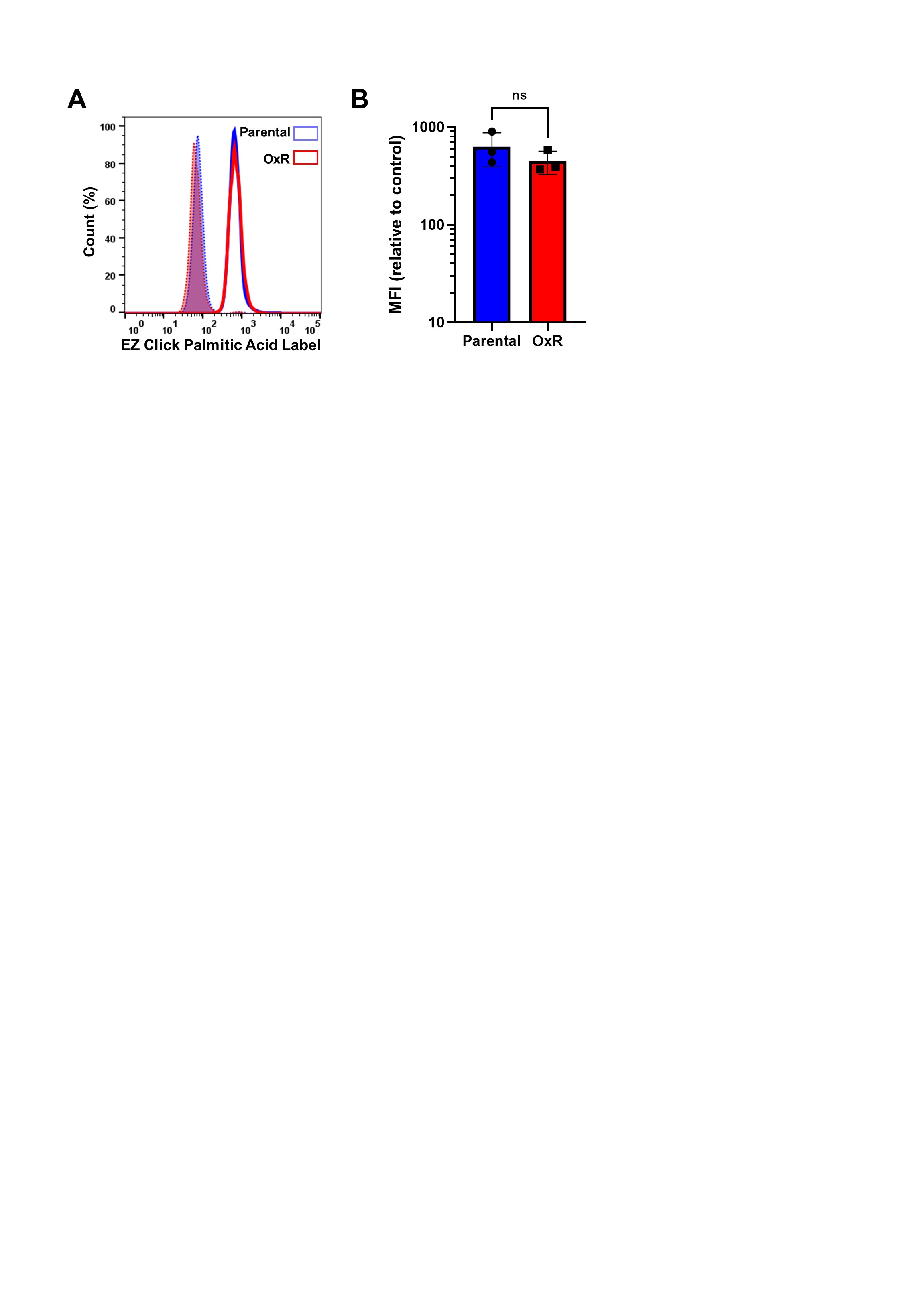

### Figure 7-figure supplement 1

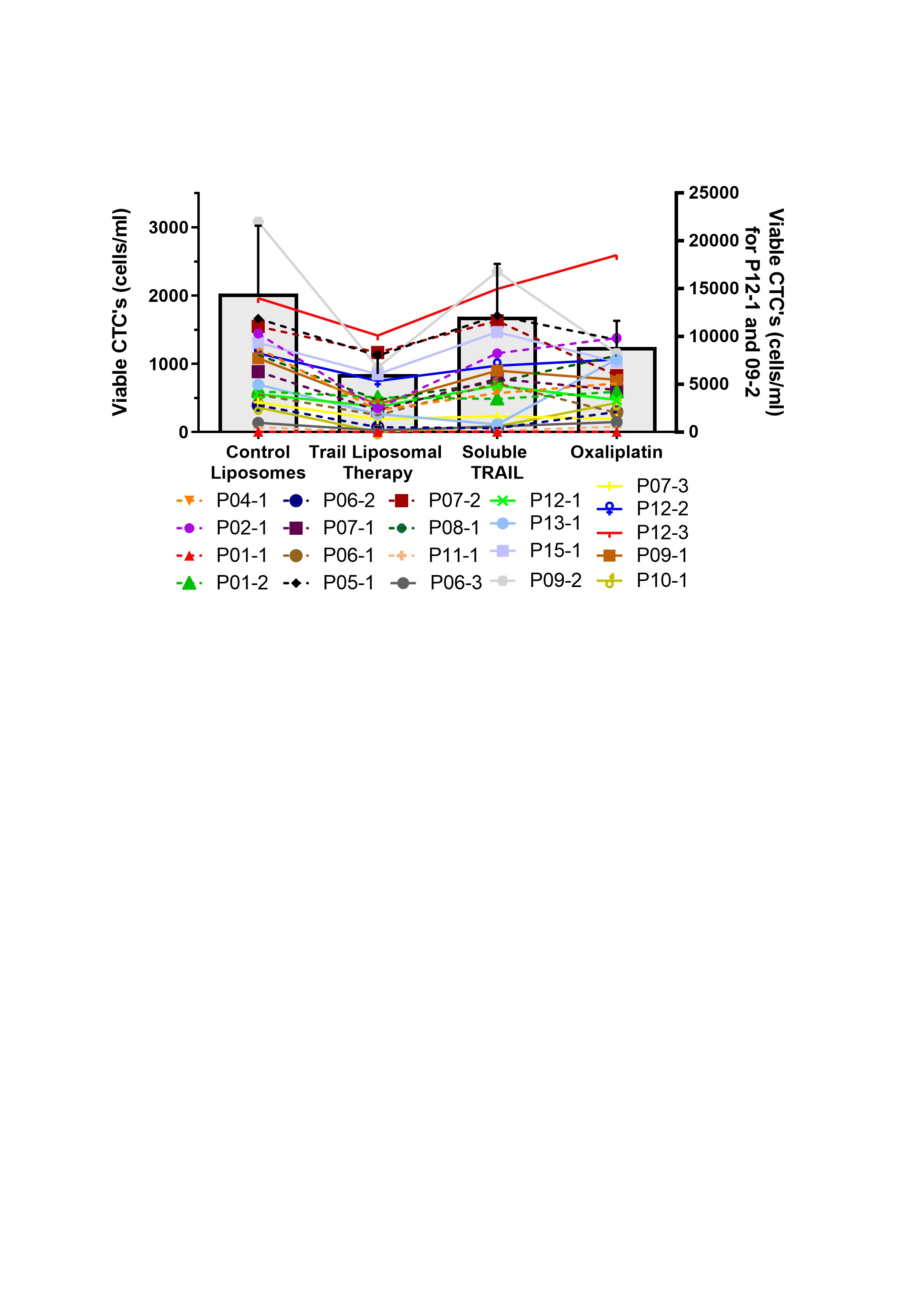

### Figure 7-figure supplement 2

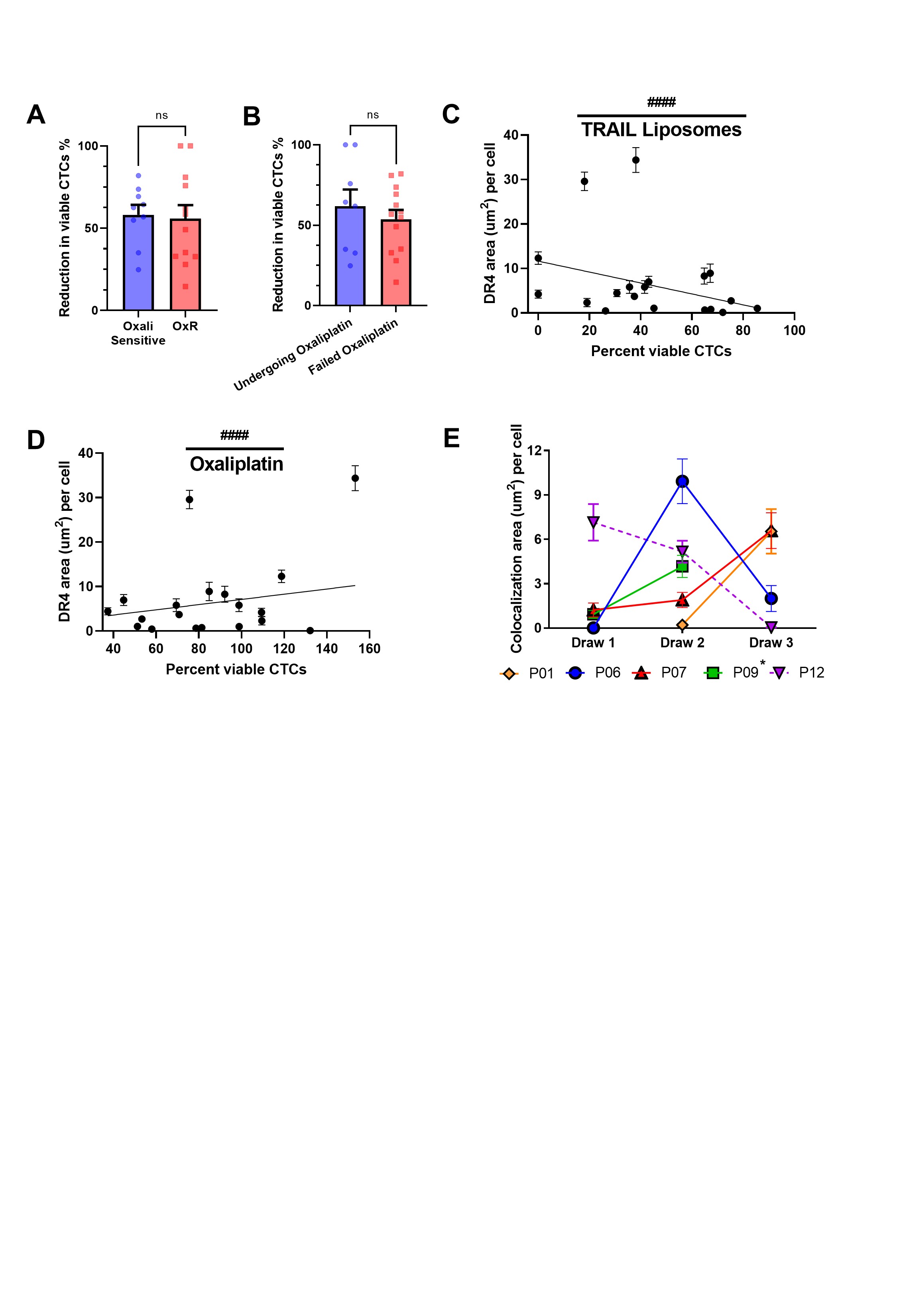

### Supplementary Table 1

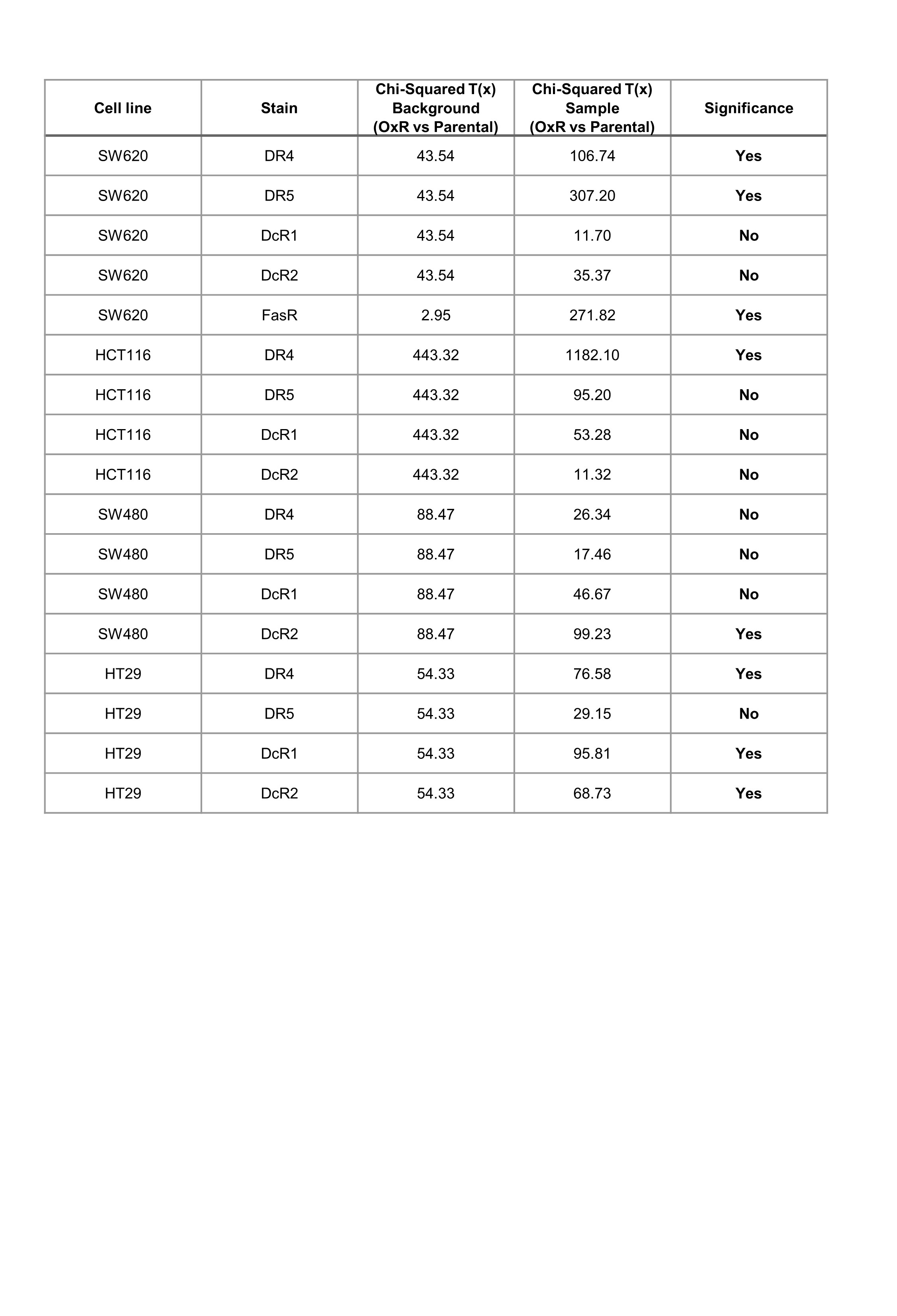
